# Computational redesign of a PETase for plastic biodegradation by the GRAPE strategy

**DOI:** 10.1101/787069

**Authors:** Yinglu Cui, Yanchun Chen, Xinyue Liu, Saijun Dong, Yu’e Tian, Yuxin Qiao, Ruchira Mitra, Jing Han, Chunli Li, Xu Han, Weidong Liu, Quan Chen, Wenbin Du, Shuangyan Tang, Hua Xiang, Haiyan Liu, Bian Wu

**Affiliations:** CAS Key Laboratory of Microbial Physiological and Metabolic Engineering, State Key Laboratory of Microbial Resources, Institute of Microbiology, Chinese Academy of Sciences, Beijing, China; University of Chinese Academy of Sciences, Beijing, China; Industrial Enzymes National Engineering Laboratory, Tianjin Institute of Industrial Biotechnology, Chinese Academy of Sciences, Tianjin, China; School of Life Sciences, Hefei National Laboratory for Physical Sciences at the Microscale, University of Science and Technology of China, Hefei, China

**Author notes:** Equal contribution (Yinglu Cui, Yanchun Chen).

## Abstract

The excessive use of plastics has been accompanied by severe ecologically damaging effects. The recent discovery of a PETase from *Ideonella sakaiensis* that decomposes poly(ethylene terephthalate) (PET) under mild conditions provides an attractive avenue for the biodegradation of plastics. However, the inherent instability of the enzyme limits its practical utilization. Here, we devised a novel computational strategy (greedy accumulated strategy for protein engineering, GRAPE). A systematic clustering analysis combined with greedy accumulation of beneficial mutations in a computationally derived library enabled the design of a variant, DuraPETase, which exhibits an apparent melting temperature that is drastically elevated by 31 °C and strikingly enhanced degradation performance toward semicrystalline PET films (23%) at mild temperatures (over two orders of magnitude improvement). The mechanism underlying the robust promotion of enzyme performance has been demonstrated via a crystal structure and molecular dynamics simulations. This work shows the capabilities of computational enzyme design to circumvent antagonistic epistatic effects and provides a valuable tool for further understanding and advancing polyester hydrolysis in the natural environment.

## Main Text

Earth’s ecosystem is approaching a planetary-scale transformation as a result of human influence^1^. Currently, a wide variety of petroleum-based synthetic polymers are produced worldwide to the extent of a 20-fold increase in annual production over the five decades since the 1960s, and a slowdown of this trend is not expected^1^. Accumulated marine and terrestrial pollution problems and the negative impacts of microplastic exposure on human health have caused escalating public and governmental concerns^3, 4^. With the excessive use of plastics and increasing pressure on the capacity for plastic waste disposal, the demand for eco-friendly waste management has assumed dramatic importance in the last few years^5, 6^.

Poly(ethylene terephthalate) (PET) is one of the most widely used man-made synthetic plastics worldwide, with an annual manufacturing capacity of over 30 million tons^7^. Its excellent durability, however, has now become an environmental detriment^8^. The aromatic terephthalate building blocks in crystalline PET largely reduce the chain mobility and lead to restricted accessibility of the ester linkages^9^. Therefore, the degradation activities of natural hydrolases have been limited by extremely low turnover rates towards PET^10^. In the past decade, there have been several foundational studies identifying enzymes from thermophilic organisms that exhibit detectable PET degrading activity^11-14^. However, these enzymes require considerably high operational temperatures near the PET glass transition temperature (∼76 °C), severely hampering their utility in environmental remediation. Recently, a major breakthrough was achieved by Yoshida et al., who reported a new PETase from *Ideonella sakaiensis* 201-F6 (*Is*PETase) with the highest activity among PET-degrading enzymes at ambient conditions^15^, which indicates a promising application for *in situ* disposal of hard-to-collect plastic disintegrates with low energy consumption. Although *Is*PETase has caught much attention for its intriguing capability, the enzyme exhibits poor durability: most of its activity is lost within 24 h of incubation at 37 °C^16^. Apart from this, *Is*PETase exhibits only reasonable activity toward low-crystallinity PET (1.9%), whereas most PET applications, including commercial containers, employ high-crystallinity PET with 30%-50% crystallinity^17^.

Not content with the finite repertoire of naturally occurring enzymes, a number of research groups have studied the engineering of *Is*PETase, which has been recently summarized by Taniguchi^18^. Lately reported crystal structures of PET-degrading enzymes allow for methods that exploit rational design to improve the PET degradation activity^19-25^. Successful single point mutations afford 1.2- to 3.1-fold higher affinity for PET^22-26^, and very recently, Kim’s group has succeeded in creating an *Is*PETase^S121E/D186H/R280A^ variant with enhanced thermal stability (by 8.81 °C) and 14-fold higher PET degradation at 40 °C^16^. Despite these attempts, there remains a key challenge to whether protein engineering achieves improved enzymatic stability and effective catalytic conversion of high-crystallinity PET by allowing large jumps along the sequence fitness landscape. There is usually a pathway whereby some new functions could be acquired by individually beneficial mutations; however, when the desired function is beyond what a single mutation or double mutations can accomplish, the number of possible paths grows exponentially as the mutations accumulate, and most paths result in downhill or even unfolded proteins^27^. Since a majority of protein engineering studies involve simple uphill walks, the main demand lies in identifying an efficient path of accumulated mutations to achieve the desired protein performance^28^.

The last few years have witnessed impressive progresses in dealing with multidimensional space by a number of metaheuristic methods^29^. Inspired by the widespread greedy algorithm applications in artificial intelligence, we introduce a novel strategy, termed GRAPE, to effectively tackle the evolutionary hard problem by computational design that enhances the probability of discovering the adaptive routes to improved fitness. The computationally redesigned *Is*PETase (DuraPETase) derived from this campaign exhibited an apparent melting temperature increased by 31 °C with good performance toward semicrystalline PET and vastly improved long-term survival under mild conditions. The enzymatic PET degradation was enhanced by over 300-fold at 37 °C for 10 days. Ameliorated biodegradation of other plastics, such as poly(butylene terephthalate) (PBT) and poly(ethylene 2,6-naphthalenedicarboxylate) (PEN), suggested DuraPETase a broad ability to degrade semiaromatic polyesters. To evaluate the underlying molecular mechanism for enhanced performance, a three-dimensional crystal structure of the variant was determined; this structure highlighted a fine-tuned ‘aromatic tunnel’ flanked by synergistic hydrophobic interactions.

## Results

### Computational redesign of a robust DuraPETase using the GRAPE strategy

The GRAPE strategy introduced here uses a greedy strategy for global optimization of mutations in each cluster to create functional variations and select the fittest variants to direct the search to higher elevations on the fitness landscape. The initial step in the GRAPE strategy (Scheme 1) consists of computational predictions of potentially stabilizing mutations along the whole protein sequence. We previously developed the ABACUS^30^ software for *de novo* protein design based on a statistical energy function. Here, this algorithm was applied together with three complementary algorithms, FoldX (force field-based energy function)^31^, Rosetta_ddg (force field-based energy function)^32^ and Consensus Analysis (phylogeny-based method)^33^, to improve the protein stability. Mutations with an ABACUS energy < −3 A.e.u., a folding free energy (ΔΔG_fold_) < −5 kJ/mol or a consensus score > 2.0 were selected (Table S1 and Figure S1). A total of 253 unique predicted mutations were obtained as potentially stabilized candidates after filtering, and ABACUS, FoldX, Rosetta_ddg, and consensus analysis algorithms provided 85, 63, 65, and 54 of these 253 mutations, respectively, with some overlap. In the second step, the potentially stabilized candidates were inspected for biophysical pitfalls, such as the introduction of internal cavities, loss of hydrogen-bonding interactions, or exposure of hydrophobic residues at the surface of the enzyme, providing a sublibrary of 85 candidates. After experimental validation, 21 well-expressed mutants displayed increased stability (> 1.5 °C increase in the apparent *T*_*m*_) (Figures S2 and S3). However, owing to the ubiquity of epistatic effects^34-36^, these positive mutations may not cooperate to reach the target function. Indeed, an *Is*PETase variant containing all 21 beneficial mutations is completely inactive.

Hence, the next step, which systematically accumulates the beneficial variants in the well-defined library, is crucial for the GRAPE strategy. Searching in the high-dimensional space involves an incomprehensibly large number of possible pathways among which only an infinitesimal fraction can escape from one local optima trap to reach a better solution. The proposed GRAPE strategy combines the advantages of greedy and clustering algorithms to provide a viable solution to minimize experimental efforts but maximize the exploration of epistatic effects in terms of additivity and/or synergism between sets of mutations at each branching point. In the GRAPE strategy, the K-means algorithm^37^, which has been proven to be a very powerful tool for data mining problems and has been adopted to perform knowledge discovery in bioinformatics research^38^, was applied to cluster the stabilizing mutations into several groups. Variants were characterized based on their positions, efficacies, and presumed effects. Subsequently, the best individual of each population served as the parent to attract offspring to its region of the fitness landscape. Each individual in the cluster was crossed with the current global best one. The greedy algorithm accepts the newly generated individual only when its fitness is better than that of the parent. Figure 1A shows the stipulation that each individual stays in its historically best position and moves forward toward the global best position.

**Figure 1.**
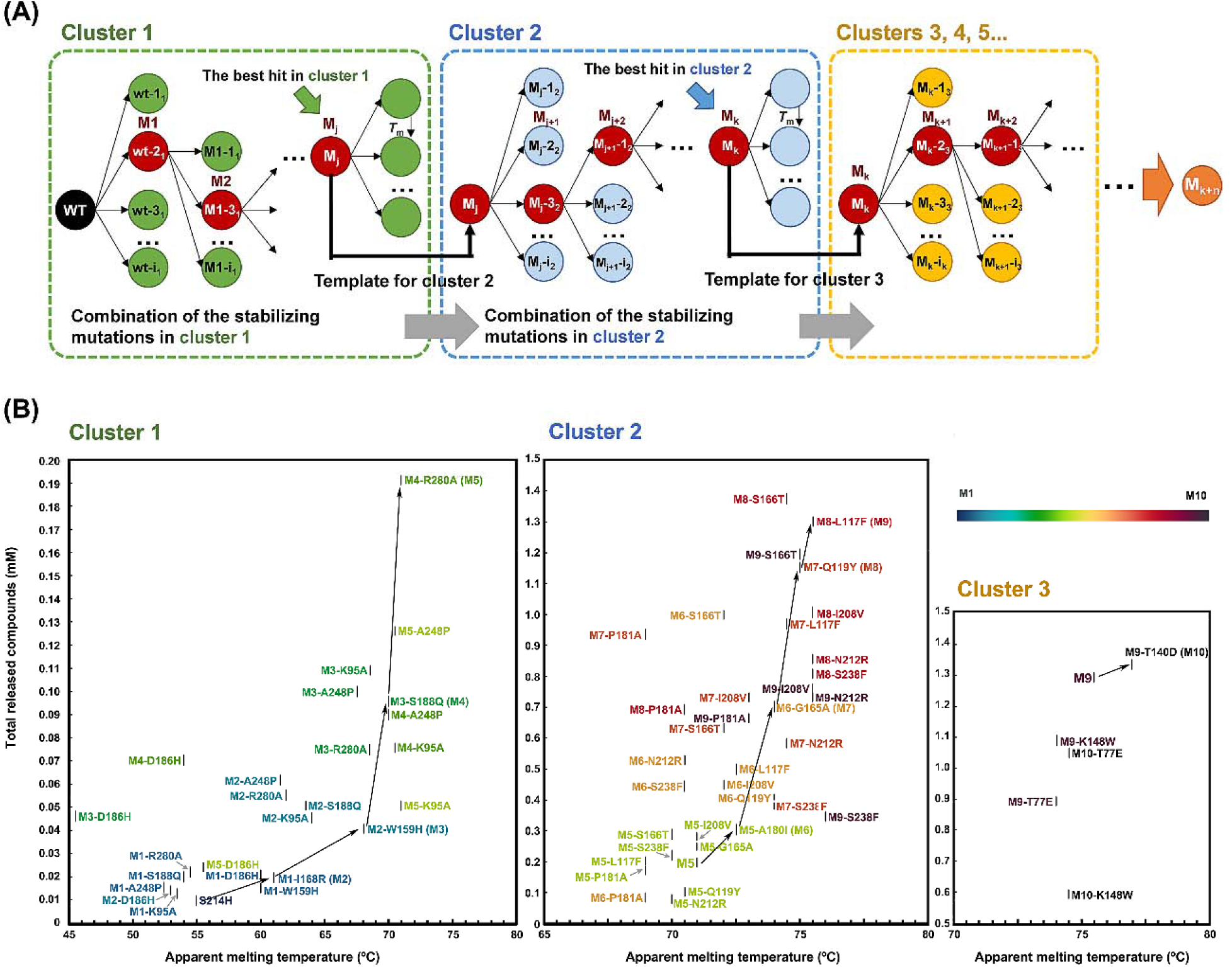
Flowchart of greedy accumulation. (A) Beneficial mutations in each cluster were crossed with the best hit of the current population until the remaining mutations in the cluster have been traversed or the *T*_*m*_ values of the combined variants decrease. If the variant showed high thermostability but an enzyme activity reduction of > 50%, the combined variant was not adopted. For example, the best hit in cluster 1 (M_j_) served as the template for further cycles of accumulation in cluster 2, whereas the best hit in cluster 2 (M_k_) was regarded as the parent for cycles in the next cluster. The best hit in each stage was sequentially referred to as M_1_, M_2_, …, M_j_, M_j+1_, …, M_k_, M_k+1_, …, M_k+n_. The red circles represent the mutants chosen from a combination stage that were used as the template for further accumulation (black arrows). (B) Enhancing the thermostability and activity of IsPETase by greedy accumulation. The accumulated mutations in each cluster are listed in Table 1.

Accordingly, the 21 identified stabilizing mutations were clustered into three groups (Table 1). S214H in the 1^st^ cluster was considered the starting point for the initial round of greedy accumulation. The exploitation process was then continued in a multihierarchical manner until all variants in the cluster were traversed or no further improvements were found (Figure 1B). Whether a combination stage was positive or deleterious was conditional on the basis of two parameters, i.e., thermostability and enzyme activity. If the combined variant showed high thermostability but seriously reduced activity, it was advisable to make a compromise. After five cycles of the accumulations, the best hit (S214H-I168R-W159H-S188Q-R280A, M5) in the 1^st^ cluster was obtained, which had a *T*_*m*_ value of 71 °C. Further combination of the remaining mutations in the 1^st^ cluster in M5 failed to increase the *T*_*m*_ value. Subsequently, the M5 variant was used as the template for crossover of mutations in the 2^nd^ cluster, leading to the best hit variant S214H-I168R-W159H-S188Q-R280A-A180I-G165A-Q119Y-L117F (*Is*PETase-M9) with a *T*_*m*_ value of 75 °C. Although the combination of the remaining mutations in the 2^nd^ cluster slightly increased the *T*_*m*_ values, degradation activities were largely reduced compared to that of M9 (Figure S4). In addition, the introduction of the S187W mutation to the best hit in each round resulted in extremely decreased expression. Therefore, M9 was set as the parent for the accumulation of other sites in the 3^rd^ cluster. The most thermostable variant, S214H-I168R-W159H-S188Q-R280A-A180I-G165A-Q119Y-L117F-T140D (*Is*PETase-M10, referred to as DuraPETase), originating from the 3^rd^ cluster, exhibited dramatically enhanced thermostability (Δ*T*_*m*_ = 31 °C). Before addressing the above challenge, we also explored a stepwise combination of the most stabilizing variants (Δ*T*_*m*_ > 7 °C). The stability of the S214H-D186H-I168R mutant was dramatically decreased, and the introduction of the P181A mutation largely reduced the enzymatic activity. Progressively addressing these multiple sites in the GRAPE scheme with multilayer upward branches allows negative trade-offs to be determined and quantified and thus finds effective ways to circumvent apparent dead ends.

**Table 1.**
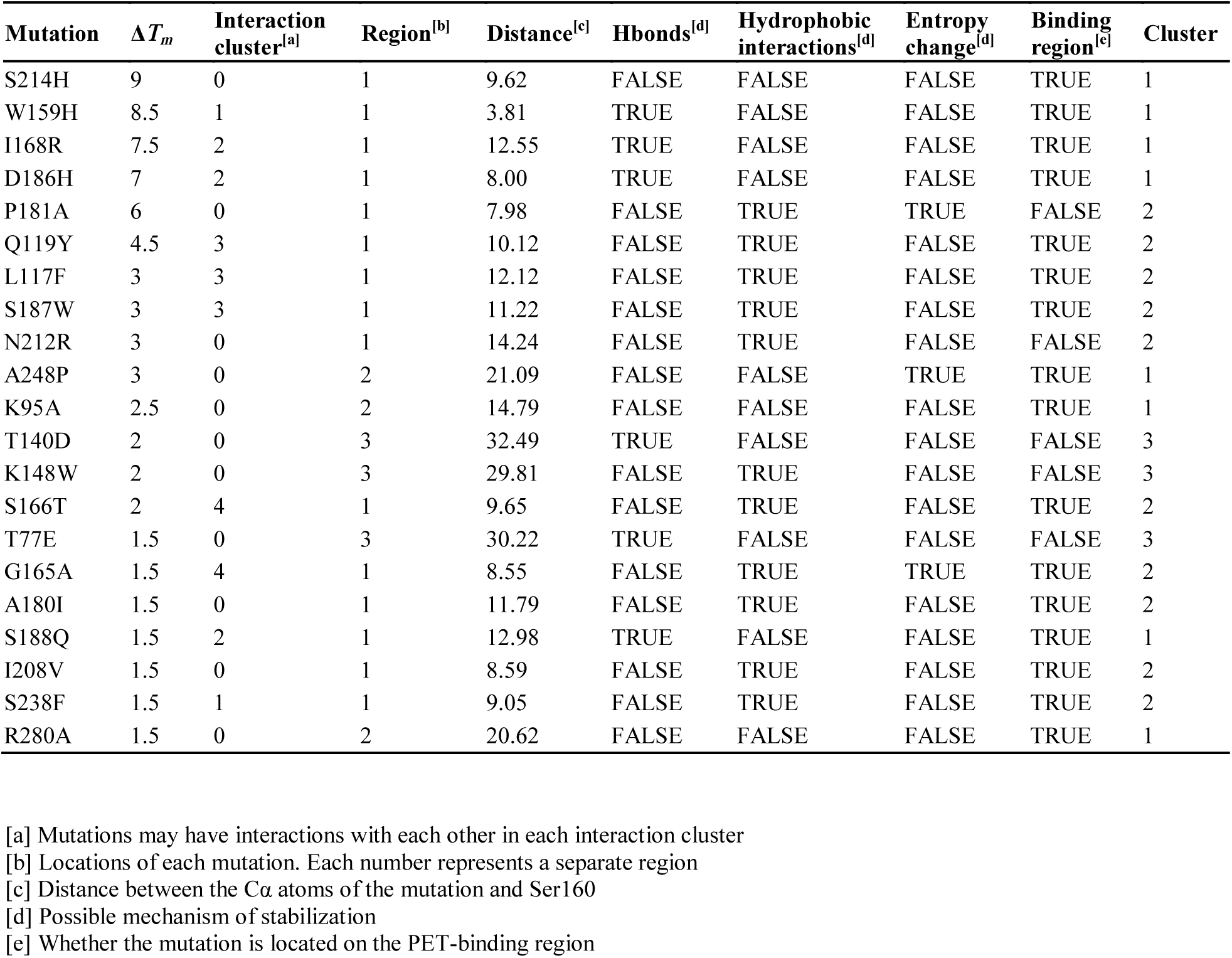
Clustering of the stabilizing point mutations based on their positions, efficacies and presumed effects.

### Enhanced PET degradation activity toward semicrystalline PET film

With the significantly enhanced performance of DuraPETase, we can explore the anticipated PET digestion of semicrystalline PET film (23%) (Figure S5). DuraPETase and the wild-type enzyme were incubated at various temperatures ranging from 37 °C to 60 °C (Figure 2A). The degradation activity profiles largely reflect the difference in inherent robustness between the designed enzyme and the wild type. *Is*PETase lost complete activity within 24 h at 37 °C, and the concentration of the degradation product reached 12 μM and did not increase thereafter, which is consistent with the results in Kim’s report^16^. When the temperature increased, the activity of *Is*PETase dropped sharply within 1 h due to thermal-induced denaturation. As expected, the enhanced thermostability of the DuraPETase coincided with remarkable improvement in long-term survival at moderate temperatures, and the enzyme even withstood elevated temperatures up to 60 °C for 3 days of incubation. The total released compound concentrations reached up to 3.1 mM when the incubation time was extended to 10 days at 37 °C, which was a 300-fold increase over the concentrations of compounds released by *Is*PETase. (Figure 2B).

**Figure 2.**
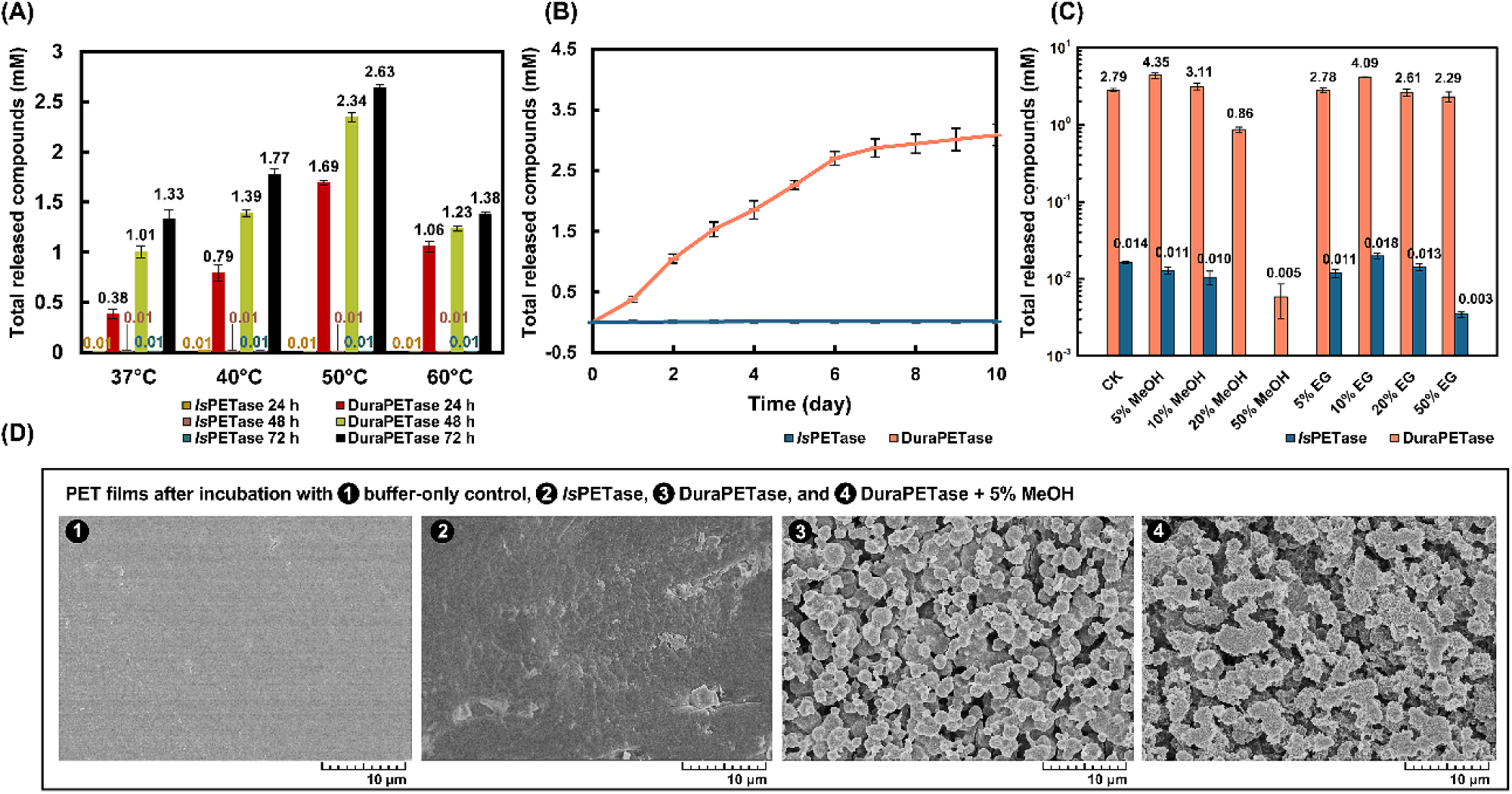
(A) Degrading product release of PET film (23% crystallinity) at different temperatures after 3 days of incubation with *Is*PETase and DuraPETase; The enzymes converted PET to terephthalic acid (TPA) and mono(2-hydroxyethyl)-TPA (MHET), with trace amounts of bis(2-hydroxyethyl)-TPA (BHET). (B) The total product release after incubation with *Is*PETase and DuraPETase for 10 days at 37 °C. (C) Effects of MeOH and EG solvents on the enzymatic activity of DuraPETase. (D) SEM images for buffer-only PET film control (➊), PET films after incubation with *Is*PETase (➋), DuraPETase (➌), and PET film after incubation with DuraPETase and 5% MeOH (➍). All SEM images were taken after 10 days of incubation at an enzyme loading of 0.01 mg/mL in glycine-NaOH buffer (pH 9.0) or with a buffer-only control. Error bars in enzyme assays (A), (B) and (C) represent the s.d. values obtained in triplicate experiments.

The high performance of DuraPETase suggested that its tolerance to organic solvents, especially ethylene glycol (EG) would be substantially enhanced, which can decrease product inhibition during the depolymerizing process. As shown in Figure 2C, DuraPETase exhibited a marked increase in tolerance to methanol (MeOH) and EG. The degradation activity was enhanced by 1.6-fold and 1.1-fold at 5% and 10% (v/v) MeOH, respectively, and approximately 30% of residual activity was maintained at 20% (v/v) MeOH. When incubated with EG solvent, DuraPETase maintained most of its activity at 5%, 20% and 50% (v/v) EG and even increased its degradation activity up to 1.5-fold when incubated with 10% (v/v) EG. This promoted performance of DuraPETase in co-solvents enhances the potential of this enzyme for auxiliary biodegradation in glycolysis and alcoholysis applications.

Polymer hydrolysis was also confirmed by scanning electron microscopy (SEM) images of the film surfaces following enzymatic hydrolysis for 10 days (Figure 2D). The buffer-only control sample exhibited a smooth and uniform surface without characteristic surface defects. The digested PET sample treated with *Is*PETase showed visual modifications with some irregular grooves of different sizes. Remarkably, severe erosion occurred with the formation of highly porous foam structures on the film surface after 10 days of degradation by DuraPETase, indicating that the biodegradation behavior was not limited to the amorphous area. The hole sizes after degradation were increased when 5% MeOH was added, and there were many spiny fragments in the corroded area. The film surface visually became nonuniform and partially broke into pieces when touched (Figure S6).

### Complete degradation of microplastics by DuraPETase

Pollution of the marine environment by micro- and even nanoplastics has become ubiquitous. Their persistence continues to increase, as they are extremely difficult to recognize and collect due to their reduced visibility. The high degradation ability of DuraPETase allows us to explore whether complete degradation of nanoplastics or even microplastics can be achieved. The total released product determined by HPLC indicated complete degradation of nanoplastics (⍰ = 50-100 nm, 0.21 g/L) by DuraPETase within 1 h at 37 °C (Figure 3C). While this observation is encouraging, much slower degradation of PET microplastic was observed during the incubation period. Figure 3E shows photographs of microplastic samples (⍰= 10-50 μm, 0.29 g/L) biodegraded over a period of 20 days. The light transmittance of microplastic solvent exhibited a slow increase. After 20 days of incubation, the microplastics degraded by DuraPETase almost disappeared with respect to microplastics added to a buffer-only control, which maintained a white color. Although the *Is*PETase-degraded sample was slightly clarified, its optical transparency was significantly lower than that of samples incubated with DuraPETase. In addition, HPLC analysis of the degradation product concentration reached 1.26 mM, which also indicated > 80% degradation of microplastics by DuraPETase.

**Figure 3.**
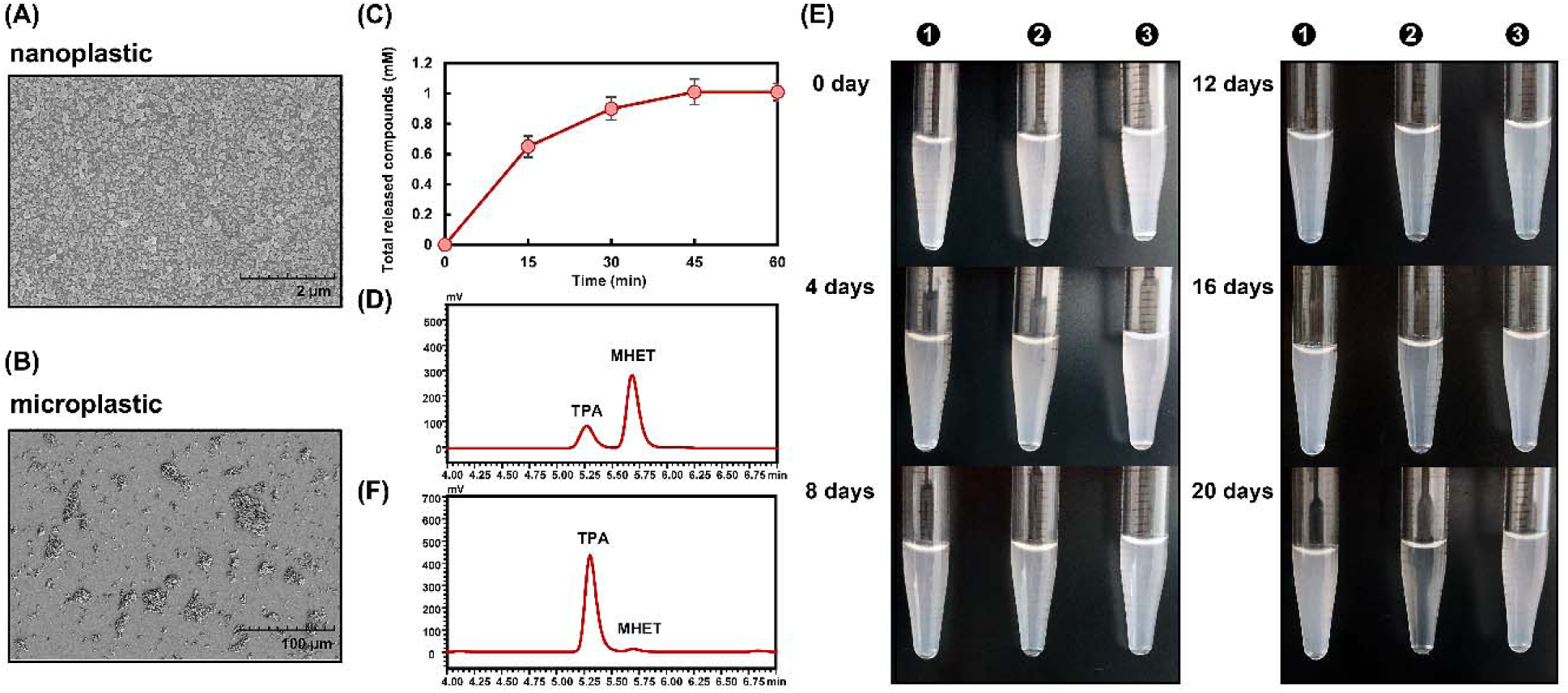
(A) SEM image of PET nanoplastics before incubation. (B) SEM image of PET microplastics before incubation. (C) Hydrolytic activity of DuraPETase toward PET nanoplastics. PET nanoparticles (200 µL, 0.53 g/L) were incubated with 10 μL of enzyme (stock concentration 0.1 mg/mL) in 290 μL of 50 mM glycine-NaOH buffer (pH 9.0) at 37 °C for 1 h.(D) High-performance liquid chromatography chromatogram of the BHET, MHET and TPA products released from the nanoplastics. (E) Time course of PET microplastic degradation. Photographs of optical transparency changes of microplastics after incubation with *Is*PETase (➊), DuraPETase (➋), buffer-only control (➌).The PET microparticles (200 µL, 1.43 g/L) were incubated with 50 μL of enzyme (stock concentration 0.5 mg/mL) in 750 μL of 50 mM glycine-NaOH buffer (pH 9.0) at 37 °C in 0 and ➊, and ➋ 50 μL of *Is*PETase and DuraPETase were replenished into the solvents after 10 days, respectively. (F) High-performance liquid chromatography chromatogram of the MHET and TPA products released from the microplastics after 20 days.

### Extending the potential applications of DuraPETase

To extend the potential applications of DuraPETase, we also examined the use of this enzyme for the degradation of other semiaromatic polyesters, including PEN and PBT. PEN is similar to PET in chemical structure but contains naphthyl rings, which produce much stiffer molecular chains than the phenyl rings of PET. Thus, PEN exhibits a high *T*_g_ (120 °C) and good tensile and barrier properties^39^. PBT exhibits higher tensile, flexural and dielectric strengths and faster, more economical molding characteristics than many thermosets. Although the degradation product of PBT is TPA, the longer aliphatic chain in butanediol than in EG endows the PBT’s excellent resistance to a broad range of chemicals at room temperature (*T*_g_ ≈ 50 °C)^40^.

Biodegradation of PBT was barely accomplished by *Is*PETase due to the notably high specificity of the enzyme toward aromatic groups rather than aliphatic chains (Figure 4). However, a gradual increase in degradation product release was found for DuraPETase at 37 °C, although the degradation activity was substantially reduced with respect to the degradation of PET. SEM analysis also supported this conclusion. In contrast to the slow degradation of PBT, significant enhancement of PEN degradation by DuraPETase was observed. Over 10 days of incubation, DuraPETase produced a maximum product concentration of 48 μM with PEN film, presumably reflecting the high hydrolysis specificity of DuraPETase toward semiaromatic polyesters due to the redesigned hydrophobic active cleft, even though the degradation activity was still lower than that targeting PET. As the introduction of the naphthalene ring into the main chain stiffens the polymer chains and largely enhances their dielectric and mechanical properties, PEN holds potential for food packaging, high-performance industrial fiber, and flexible printed circuit applications. Different types of such high-performance plastic would eventually be dispersed into the environment. These results therefore inspire further engineering to improve the depolymerization of new classes of semiaromatic polyesters.

**Figure 4.**
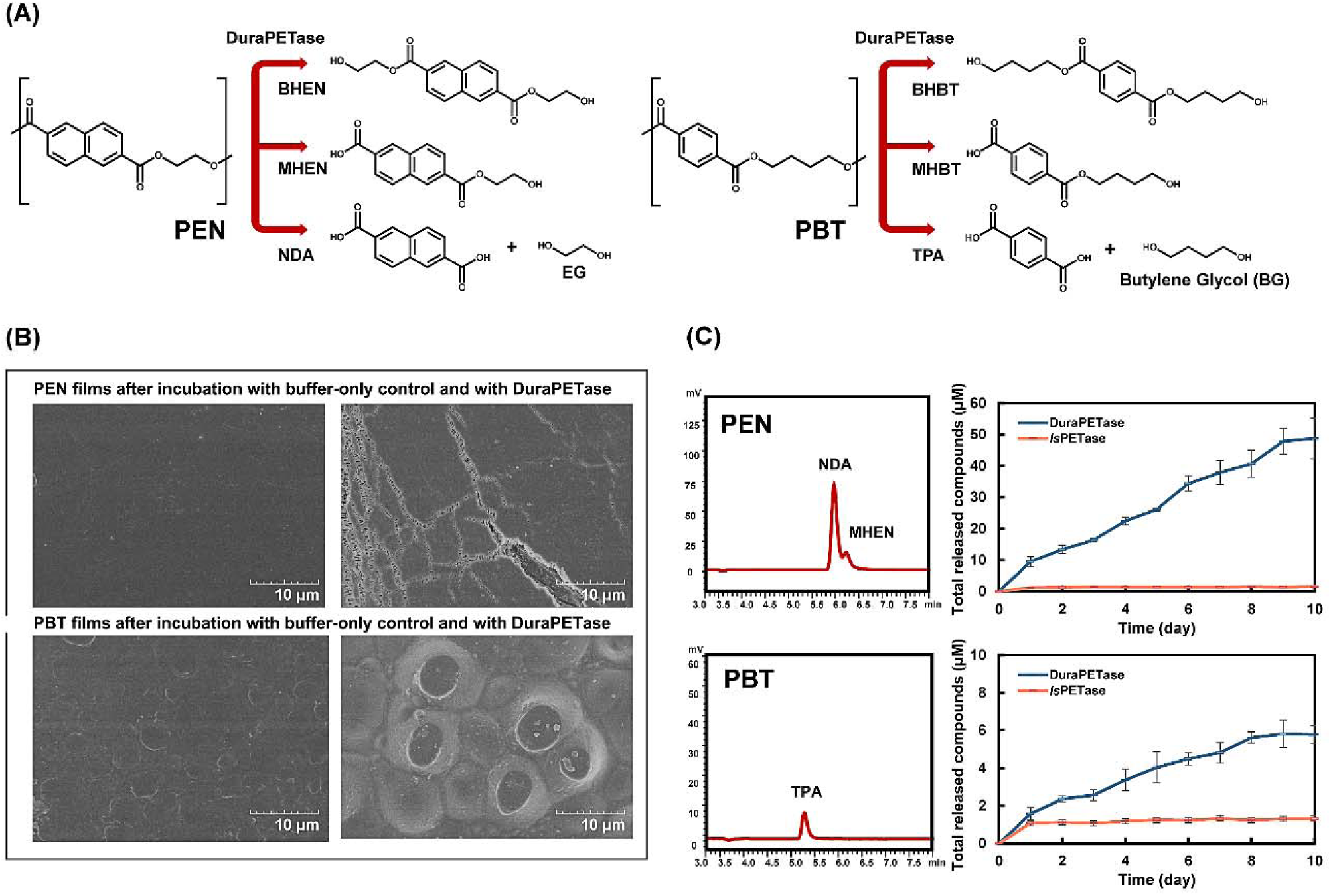
(A) PEN and PBT degradation. DuraPETase catalyzes the hydrolytic cleavage of PEN to produce 2,6-*naphthalenedicarbSoxylic* acid (NDA), bis(2-hydroxyethyl)-NDA (BHEN), and mono(2-hydroxyethyl)-NDA (MHEN) and converts PBT to bis(4-hydroxybutyl)-TPA (BHBT), mono(4-hydroxybutyl)-TPA (MHBT), and TPA. (B) SEM images of PEN and PBT films after incubation with buffer only control (left) and with DuraPETase (right). All SEM images were taken after 10 days of incubation at an enzyme loading of 0.01 mg/mL in glycine-NaOH buffer (pH 9.0) or with a buffer-only control at 37 °C. (C) High-performance liquid chromatography chromatogram of the products released from the PEN and PBT films. Error bars represent s.d. values obtained in triplicate experiments.

### Mechanism underlying the improved properties of DuraPETase

To further evaluate the molecular basis for the improved properties of the obtained variant, we determined the crystal structure of DuraPETase at 1.63 Å resolution (PDB ID: 6KY5), which revealed a conserved α/β hydrolase fold, with a core consisting of seven α-helices and nine β-sheets. The DuraPETase-PET binding mode predicted by molecular docking was similar to the mode observed in the crystal structure of *Is*PETase bound to 1-(2-hydroxyethyl) 4-methyl terephthalate (HEMT)^21^. The distance between the carbonyl carbon of the PET polymer and the C_α_ atom of Ser160 was 2.75 Å, which was suggested to be within the relevant reactive distance for nucleophilic attack. The oxyanion of the tetrahedral intermediate is stabilized by an oxyanion hole involving backbone NH groups of Tyr87 and Met161. In addition to the interaction information observed crystallographically, we explored MD simulations to analyze potential effects of the mutations on global structure dynamics.

According to the crystal structure and MD simulations, key aspects of the molecular mechanism by which the mutations improve the protein robustness were proposed as follows: introduction of new electrostatic interactions (T140D, W159H, I168R and S188Q), improvement of hydrophobic packing in the protein surface and interior (L117F, Q119Y, A180I, S214H and R280A), and reduction in the conformational entropy of a local coil region (G165A) (Figure S7).

For the T140D mutation, the substitution of aspartate for threonine in T140D mutation conferred new hydrogen bonds with the hydroxyl group of Ser142 in the crystal structure, which maintained for 93.18% of the simulation time. The N_ε_ atom of W159H formed new hydrogen-bonding interactions with the backbone oxygen atom of H237 in the crystal structure. However, this W159H mutation was observed to restore the active site by forming a new hydrogen bond with Ser160 in the simulations. The original catalytic residue His237 was concomitantly flipped “up” out of the catalytic triad to play an aromatic stabilization role with PET (Figure 5B). This reconstruction of the catalytic triad was also suggested by Austin et al.^20^. For I168R and S188Q mutations, new salt bridge interactions were formed between I168R and D186 in the crystal structure. The guanidine group is also suggested to donate new hydrogen bonds to the amide oxygen atom of S188Q in the MD simulations for 10.36% of trajectories to stabilize the enzyme structure, which may explain the negative epistatic interactions when D186H was added to S214H-I168R-W159H (M3) and S214H-I168R-W159H-S188Q (M4).

**Figure 5.**
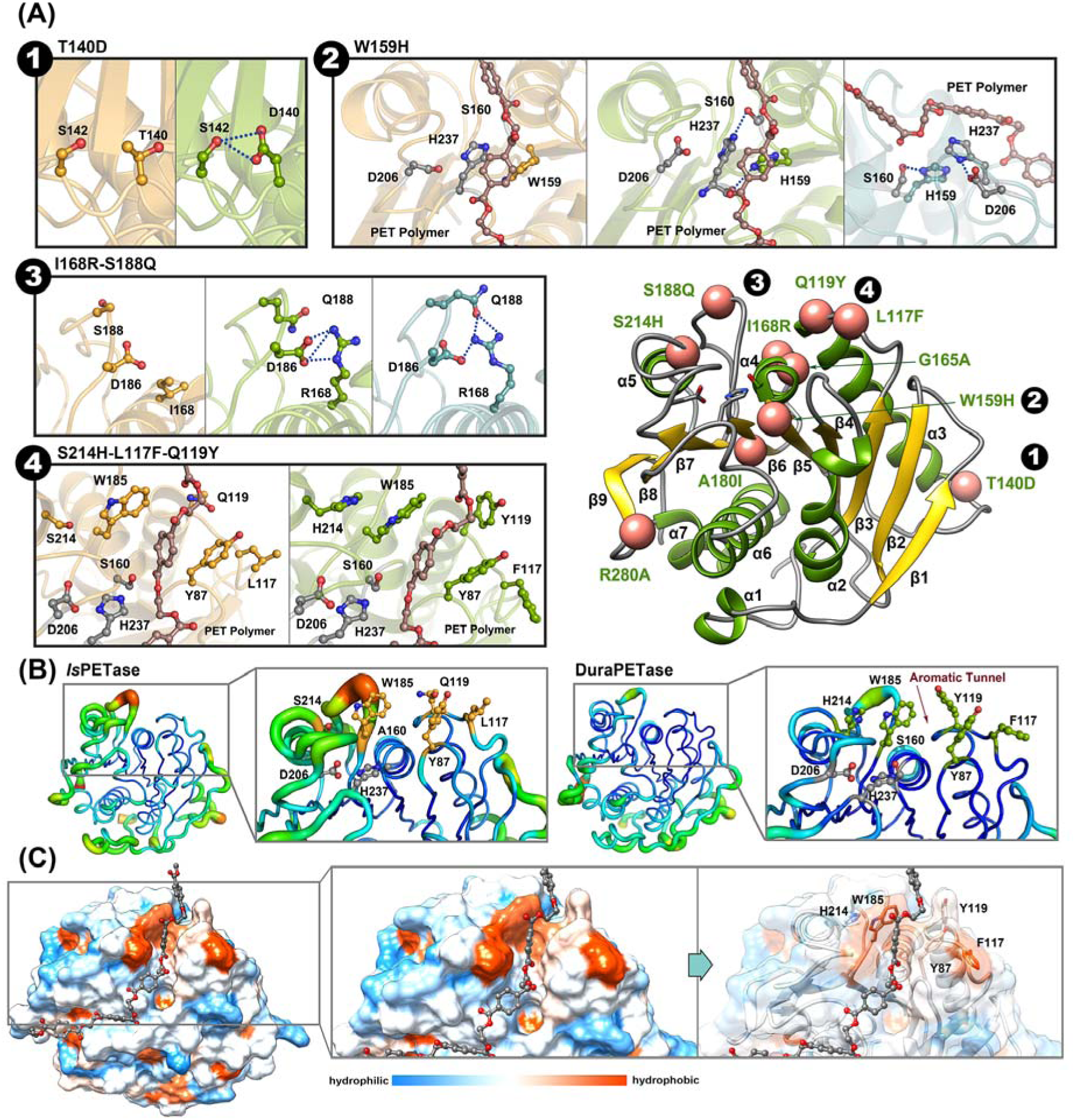
Structural features of DuraPETase. (A) Structural effects of ➊T140D, ➋ W159H, ➌ I168R and ➍ S188Q, and 0 S214H, L117F and Q119Y mutations in the DuraPETase crystal structure docked with PET molecule (green) or in MD simulations (blue) compared to those of the *Is*PETase crystal structure docked with PET molecule (bright orange). Key residues proximal to the stabilizing mutations are shown in ball and stick representations. The catalytic triad is colored in gray. Right bottom panel: Location of the stabilizing mutations in the crystal structure of DuraPETase (PDB ID: 6KY5). The DuraPETase structure is shown as a cartoon, while the Cα atoms of the stabilizing mutations are shown as coral atoms. (B) B factors of the X-ray structures of *Is*PETase (PDB ID: 5XH3) and DuraPETase, indicating global changes in protein flexibility, especially for the active site region. (C) Hydrophobic surface of the DuraPETase crystal structure and that docked with PET molecule.

In addition to electrostatic effects, hydrophobic interactions have a marked influence on enzymatic performance (Figure S7). It is noteworthy that the benzene ring of the PET molecule stands at an active-site crevice that consists of five mutually contacting phenylalanine, tyrosine, tryptophan, and histidine side chains in DuraPETase-PET binding mode. These key residues may provide an “induced-fit” arrangement of the active site cleft. Residues Tyr87 and Trp185 were observed to form continuous π-π interactions with the aromatic motif of the PET polymer for 54.50% and 31.96% of the entire trajectory, respectively. Simultaneously, L117F and Q119Y were located in proximity to the active site and promoted “T-shaped” π-π interactions with Tyr87 (the oxyanion hole-forming residue) for 71.42% and 68.62% of the trajectory, respectively. The stability of the oxyanion hole may be enhanced, and the tetrahedral intermediate can be further stabilized. Another substitution in the active site, S214H was suggested to prevent the wobbling of Trp185, and it formed an offset parallel π-π stacking interaction with Trp185 for 60.48% of the simulation time. Therefore, the crystal structure and MD simulations suggested a well-organized hydrophobic domain involving S214H, Trp185, Tyr87, L117F, and Q119Y, which we termed as ‘aromatic tunnel’ flanker, to finely tune the active-site cleft suitable for binding the aromatic motif of PET. Although the flexibility of the active site region has been largely reduced (Figure 5B), the hydrophobic substitutions may significantly enhance the binding affinity for PET polymer (Figure 5C), and further enable DuraPETase to hydrolyze ester bonds along semicrystalline PET polymer chains via targeting neighboring aromatic motifs in the polymer chain backbone.

## Discussion

Durability, the greatest asset of plastic, has now become a lingering curse that results in plastic remaining in our environment for hundreds of years^8^. Even when physically broken, plastics never truly leave the environment but are present as micro- and nanoplastics that are choking marine life and propagating up the food chain. The biodegradation of plastics under ambient conditions, especially for uncollectable microplastics, is highly desirable to enhance changes in this scenario. To this end, the seminal discovery of *Is*PETase immediately aroused immense interest from numerous research groups in investigating the mechanistic basis of its catalytic mechanism and in increasing the efficiency and stability of this exciting enzyme. Although combining random mutagenesis with high-throughput screening has proven to be a successful strategy for the modification of enzyme properties, a long-sought alternative to screening-based approaches is reliable *in silico* design of performance-enhancing mutations, especially for the degradation of insoluble solid synthetic polymers. Over the last 20 years, *in silico* design based on energy calculations has advanced greatly from fairly simple to increasingly accurate and versatile methods^46-47^ and has had a particularly positive impact in the area of protein skeleton optimization. However, the accuracy based on energy functions is still suboptimal because of several factors, including insufficient conformational sampling of the static structure, imbalances in the force fields, and intrinsic problems with existing data sets^28^. Although the drawbacks can be mitigated by using hybrid methods that incorporate complementary statistical-based approaches such as ABACUS, most stability strategies focus on single-point-mutation or simple stepwise combination processes, resulting in increased prediction errors upon application to multiple-point mutants. Whenever epistatic effects are present, the predictions are prone to frustrate optimization processes.

The GRAPE strategy proposed in this study represents a step forward in the computational enzyme design because of its capability to reduce the risk of combining mutations with antagonistic effects and independency of specific structure. In most cases the combination process fails to immediately find efficient pathways when coupled mutations have negative epistatic interactions. However, simultaneously making all possible paths creates an exponentially larger library. A systematic clustering and greedy combination strategy that has been successfully utilized in machine learning therefore provides a way to tap into the possible beneficial pathways in a well-defined library by reducing the dimensionality of the data. The computationally redesigned DuraPETase enzyme described herein has substantially improved operational robustness and semicrystalline PET degradation activity, which are important properties for its potential use in real-case applications such as surface modification of PET fibers in the textile industry and *in situ* disposal of microplastics. Both the crystal structure and MD simulations suggest an underlying ‘aromatic tunnel’ mechanism to explain the promoted enzymatic performance via a structural view, indicating a significant effect of the synergy between various beneficial mutations on stability and activity enhancement. Furthermore, Zimmermann et al. has hypothesized combinatorial exo- and endo-type or endo-type-only degradation mechanisms for different PET microstructures when they are incubated with TfCut2 near the PET glass transition temperature^48^. Encouraged by the impressive performance of DuraPETase, researchers are expected to investigate the degradation mechanism of scrap PET based on this robust enzyme at mild temperatures.

In summary, this work shows that a collection of subtle variations identified by *in silico* approaches provides clues regarding how to design a PET hydrolase to improve its incorporation of semicrystalline aromatic polyesters. We believe that the proposed GRAPE strategy constitutes a significant advance in enzyme design methodology that is complementary to traditional computational design strategies, which often focus on exquisitely tuned energy evaluations of single mutations. The variant DuraPETase, designed here by the GRAPE strategy, serves as a useful catalyst for efficient PET degradation at moderate temperatures and opens up avenues for research in decreasing environmental microplastic accumulation and understanding PET microstructure alterations upon enzymatic erosion. Despite the aforementioned achievements, complete degradation of plastic waste still presents a number of challenges. There is an urgent need to further research this topic via studies of issues such as coupling DuraPETase with downstream enzymes, degrading the TPA and EG product, and scaling up the combination degradation system for industrial applications.

## Acknowledgments

This work is supported by the National Key R&D Program of China (Grant No. 2018YFA0901600), the National Natural Science Foundation of China (Grant Nos. 31601412, 31822002, 31961133016), Beijing Municipal Natural Science Foundation (8194074), and the Biological Resources Program (KFJ-BRP-009) of the Chinese Academy of Sciences.

## Author contributions

YLC, QC performed the computational work, YLC, YCC, XYL, SJD and YET performed biochemical and biocatalytic experiments, YXQ, RM, JH, CLL performed polymer preparations and analysis, YCC, XH and WL determined the crystal structure of DuraPETase, WBD, SYT, HX, HYL and BW provided supervision and input on experimental design. YLC and BW drafted the manuscript, which was revised and approved by all authors. YLC and YCC contributed equally to this work.

## Competing interests

Authors declare no competing interests.

## Data and materials availability

All data is available in the main text or the supplementary materials.

## Supplementary Materials

**Scheme 1.**
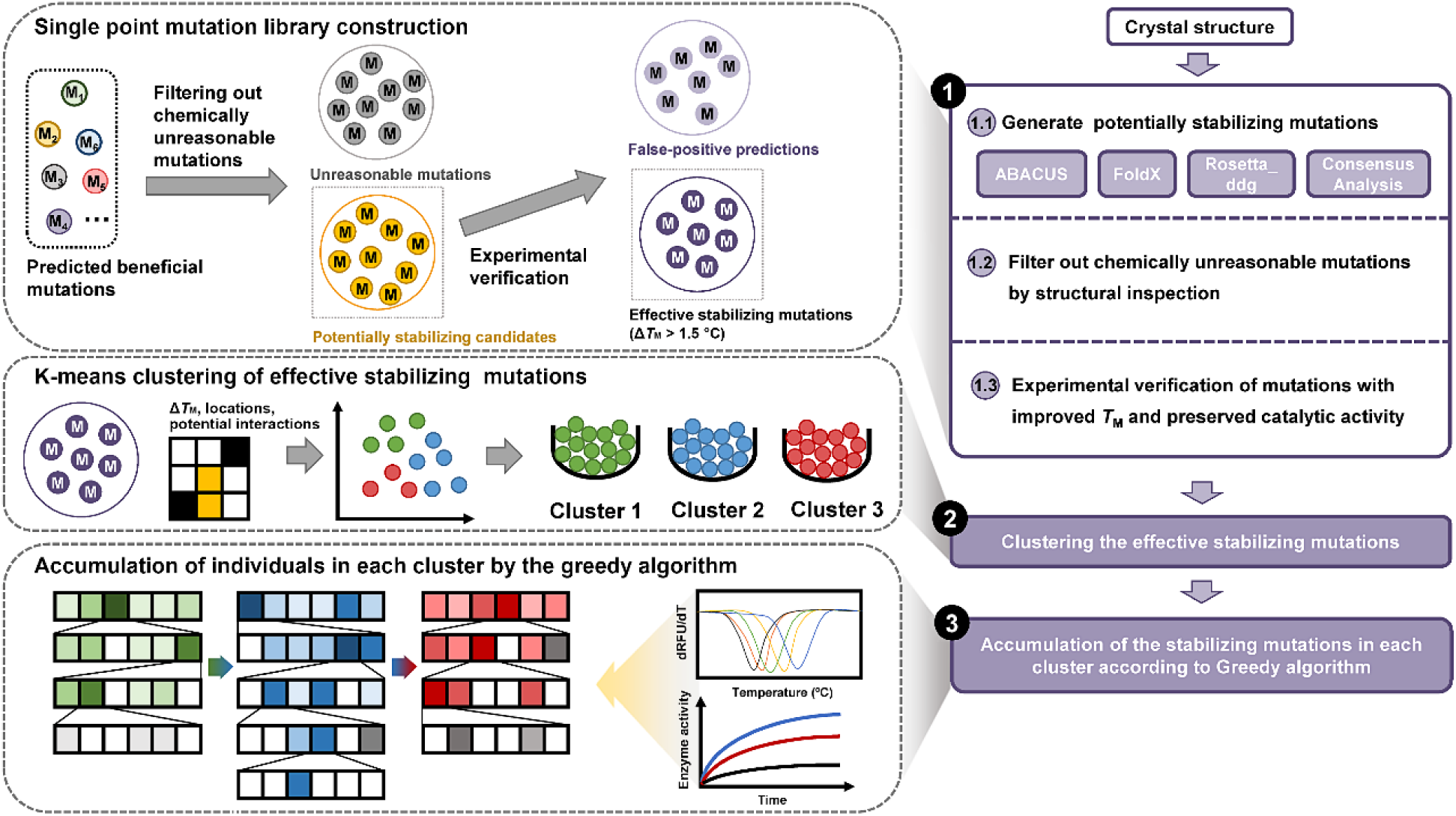
Schematic representation of the GRAPE strategy. In step 1, stabilizing mutations are generated with multiple algorithms. The computational designs with typical known pitfalls are eliminated. Then, the remaining designs are selected for experimental validation. Step 2 characterizes the variants according to their positions, efficacies and presumed effects. Accumulation of the mutations in each cluster according to the greedy algorithm is performed in Step 3. Details regarding Step 3 are demonstrated in Figure 1(A).

