## Supplementary material for "Computational redesign of a PETase for plastic biodegradation by the GRAPE strategy": SI

**This PDF file includes:**

Materials and Methods

Tables S1-S3

Figures S1-S8

References (S1-S16)

Materials and Methods

Materials

BHET, TPA and PET granular were purchased from Sigma-Aldrich Co. (St. Louis, MO, USA). MHET was obtained as a mixture with water by hydrolyzing BHET with KOH at 75 °C for 60 min. PEN powders were purchased from Xi’an Ruixi Biological Technology Co., Ltd. (Xi’an, China). PBT powders were purchased from Hezhong Bio-Chemical Co., Ltd. (Wuhan, China). All other chemicals were purchased from Sigma-Aldrich Co. in the highest available purity and were used without further purification.

PET nanoparticles and microparticles were prepared from PET granulates as described previously^1^. For nanoparticles, 0.05 g PET granulates was dissolved in 10 mL of 1,1,1,3,3,3-hexafluoro-2-propanol and filled into a burette. This solution was added dropwise with vigorous stirring by means of an UltraTurrax® (FA25-D, **FLUKO, China**) at 19,000 rpm into 35 mL ice double distilled water at a volume flow of 1 mL/min. For microparticles, 0.05 g PET granulates was dissolved in 10 mL of 1,1,1,3,3,3-hexafluoro-2-propanol. This solution was added dropwise with vigorous stirring at 10,000 rpm into 10 mL ice double distilled water at a volume flow of 1 mL/min. The precipitated polymer was filtered off with a pleated filter (597^1^/_2_ FOLDING FILTER, Schleicher und Schuell GmbH). The organic solvent was then removed from the mixture using a rotary evaporator (Rotavapor RE-2000A, Yuhua, China). After dewatering, the nanoparticles and microparticles were weighted, and the final concentrations of PET nanoparticles and microparticles were calculated as 0.53 g/L and 1.43 g/L, respectively. PET nanoparticle and microparticle sizes were determined by SEM.

The semiaromatic polymer films were prepared from the PEN and PBT powders and PET granulates. The PET granulates (0.1 g) were dissolved in 1 mL of 1,1,1,3,3,3-hexafluoro-2-propanol. Then, 1 mL solution was plated in a glass culture dish and extracted with 300 µL of acetonitrile to obtain the PET semicrystalline films. The PEN powders (0.1 g) were dissolved in 1 mL of 1,1,1,3,3,3-hexafluoro-2-propanol. Then, 1 mL solution was plated in a glass culture dish and extracted with 300 µL of acetonitrile to obtain PEN films. The PBT powders (0.1 g) were dissolved in 1 mL of 1,1,1,3,3,3-hexafluoro-2-propanol. Then, 1 mL solution was plated in a glass culture dish to obtain PBT films. The semiaromatic polymer films were cut into a circular form 8 mm in diameter.

Cloning

The gene encoding PETase from *Ideonella sakaiensis* strain 201-F6 (GenBank accession number: GAP38373.1) was commercially synthesized with codon optimization for expression in *Escherichia coli* cells (GenScript, Nanjing, China). The nucleotide sequence corresponding to the signal peptide was removed from the synthetic DNA. The synthesized gene was then cloned into pET-21b at *Nde*I and *Xho*I digestion sites and transformed in *E. coli* C41(DE3).

Site-directed mutagenesis

Single point mutations were commercially constructed with the wild-type PETase plasmid as a template (GenScript, Nanjing, China). Variants during the accumulation step were constructed using a QuickChange site-directed mutagenesis kit (Agilent Technologies, Santa Clara, CA, USA). The sequences of the mutagenesis oligonucleotides are listed in Supplementary Table 1. The PCR products were incubated with DpnI (New England Biolabs, Ipswich, MA, USA) to digest the original DNA template and then separately transformed into *E. coli* TOP10. The introduced mutations were confirmed by sequencing.

Protein purification

The pET21b-PETase plasmid containing either wild type or variants was transformed into *E. coli* C41 (DE3) cells that were grown in 2YT medium at 37 °C to an OD_600_ of ~1.0 and then induced by 1 mM isopropyl β-D-thiogalactopyranoside (IPTG) at 16 °C for 20 h. Cells were harvested by centrifugation at 10,000 × g for 10 min; resuspended in a lysis buffer containing 50 mM Na_2_HPO_4_, 100 mM NaCl and 30 mM imidazole (pH 7.5); and then disrupted by ultrasonication. The cell extract was obtained after removing precipitates by centrifugation (13,000 × g, 1 h, 4 °C) and filtration (0.22-μm filter, Millex). The supernatant was then applied to a 5-mL HisTrap HP column (GE Healthcare, Milwaukee, WI, USA). After washing unbound proteins, the target protein was eluted at 300 mM imidazole, and then the buffer was exchanged for 50 mM Na_2_HPO_4_, 100 mM NaCl (pH 7.5) using a HiPrep 26/10 Desalting column (GE Healthcare, USA). The purified enzyme was stored at 4 °C.

Crystallization, data collection, structure determination and refinement

All crystallization experiments were conducted at 22 °C using the sitting-drop vapor-diffusion method. In general, 1 μL DuraPETase containing solution (25 mM Tris-HCl, 150 mM NaCl, pH 7.5; 20 mg/mL) was mixed with 1 μL of reservoir solution in 48-well Cryschem Plates and equilibrated against 100 μL of the reservoir solution. The optimized crystallization condition of DuraPETase was 1.9 M (NH_4_)_2_SO_4_. Within 5 to 6 days, the crystal sizes were suitable for X-ray diffraction.

All of the X-ray diffraction data sets were collected at National Synchrotron Radiation Research Center (NSRRC) in Hsinchu, Taiwan. The diffraction images were processed using HKL2000. Further refinement was performed using the programs Refmac5^2^ and Coot^3^. Briefly, 5% of randomly selected reflections were set aside for calculating R_free_ as a monitor. DuraPETase structure was determined by MR method with Phaser using the refined PETase structure as a search model. The 2*F*_o_-*F*_c_ difference Fourier maps showed clear electron densities for most amino acid residues. The data collection and refinement statistics are summarized in Table S3.

Determination of apparent melting temperatures

A fluorescence-based thermal stability assay was used to determine apparent melting temperatures. Protein solution (20 µL) in the buffer (50 mM Na_2_HPO_4_, 100 mM NaCl, pH 7.5) was mixed with 5 µL 100-fold diluted SYPRO Orange dye (Molecular Probes, Life Technologies, USA) in a thin-walled 96-well PCR plate. The plate was sealed with optical-quality sealing tape and heated in a CFX 96 real-time polymerase chain reaction (PCR) system (BioRad, Hercules, CA, USA) from 25 to 90 °C at a heating rate of 1.625 °C/min. Fluorescence changes were monitored with a charge-coupled device (CCD) camera. The wavelengths for excitation and emission were 490 and 575 nm, respectively. To obtain the temperature midpoint for the protein unfolding transition, a Boltzmann model was used to fit the fluorescence data obtained by the CCD detector:

$I=(A+\frac{(B-A)}{1+e^{\frac{(T_{m}-T)}{C}}})$	(1)

where *A* and *B* are pretransitional and posttransitional fluorescence intensities. *C* is a slope factor. *I* is the fluorescence intensity at temperature *T*^4^. Data points after the fluorescence intensity maximum were excluded from fitting.

HPLC analysis

HPLC analysis was performed on an LC-2030C HT system (SHIMADZU, Japan) equipped with a UniSil® 5-120 C18 Aq column (4.6 × 250 mm, 5 μm, Nano-Micro Technology Co, Ltd, Suzhou, China). The mobile phase was composed of 70% buffer A (0.1% formic acid in distilled water) and 30% buffer B (acetonitrile). The flow rate was 0.8 mL/min. Separation was performed at 25 °C with detection at 260 nm.

Enzyme assays for nanoparticles

To evaluate the performance of 85 promising single point variants, we used PET nanoparticles as the substrate for fast screening. The PET nanoparticles (200 uL) were added in 290 μL of 50 mM glycine-NaOH buffer (pH 9.0) with 10 μL of enzyme (stock concentration 0.1 mg/mL). The reaction mixture was incubated at 37 °C for 1 h. The reaction was terminated by heat treatment (100 °C, 10 min). The supernatant obtained by centrifugation (18,000 × g, 5 min) was then analyzed by HPLC.

Enzyme assays for PET film

The PET film (⌀8 mm) was soaked with 10 μL of enzyme (stock concentration 0.5 mg/mL) in 490 μL buffer containing 50 mM glycine-NaOH (pH 9.0). For enzyme assays in the combination stage, the reaction mixture was incubated at 37 °C for 48 h. The reaction was terminated by heat treatment (100 °C, 10 min). The supernatant obtained by centrifugation (18,000 × g, 5 min) was then analyzed by HPLC.

Durability of DuraPETase for PET film degradation

The PET film (⌀8 mm) was soaked with 10 µL of enzymes (stock concentration 0.5 mg/mL) in 490 μL buffer containing 50 mM glycine-NaOH (pH 9.0) at different temperatures (37 °C, 40 °C, 50 °C and 60 °C) for 10 days. The reaction was terminated by heat treatment (100 °C, 10 min). The supernatant obtained by centrifugation (18,000 × g, 5 min) was then analyzed by HPLC.

Enzyme assays for microparticles

PET microparticles (200 μL) were added in 750 μl of 50 mM glycine-NaOH buffer (pH 9.0) with 50 µL of enzyme (stock concentration 0.5 mg/mL). The reaction mixture was incubated at 37 °C for 20 days, and 50 μL of *Is*PETase and DuraPETase were replenished into the solvents after 10 days, respectively.

Enzyme assays for semiaromatic polymer films

The PEN and PBT films (⌀8 mm) were soaked with 10 μL of DuraPETase and *Is*PETase (stock concentration 0.5 mg/mL) in 490 μL buffer containing 50 mM glycine-NaOH (pH 9.0). The reaction mixture was incubated at 37 °C for 10 days. The reaction was terminated by heat treatment (100 °C, 10 min). The supernatant obtained by centrifugation (18,000 × g, 5 min) was then analyzed by HPLC.

Effects of organic solvents on enzyme activity

The effects of organic solvents on DuraPETase and *Is*PETase were determined by incubating the enzymes with a series of concentrations of methanol and ethylene glycol (0-50%) with PET film (⌀8 mm) in buffer containing 50 mM glycine-NaOH (pH 9.0) at 37 °C for 10 days. The concentration of the enzymes used for all of the enzyme assays were 0.01 mg/mL in 500 μL of the solvent.

Scanning electron microscopy

After cultivation, the films were washed with 1% SDS, distilled water, and then ethanol. The morphology of PET films before and after enzyme exposure was examined by SU8010 SEM (Hitachi, Tokyo, Japan) at an accelerating voltage of 5 kV. Samples were sputter-coated with platinum in an ion sputter (E1045, Hitachi, Japan).

Differential Scanning Calorimetry (DSC)

The crystallinity of PET, PEN, and PBT films were analyzed by differential scanning calorimetry instrument (DSC Q2000). The following protocol was used for each sample. A sample of 2-5 mg in an aluminum pan was cooled from room temperature to 0 °C. The pan was then heated from 0 to 280 °C at 10 °C min^-1^ and then maintained at 280 °C for 1 min under a nitrogen atmosphere. Subsequently, the sample was quenched to 0 °C at 10 °C min^-1^. The heat of fusion ΔH_m_ and cold crystallization ΔH_c_ were determined by integrating areas (J g^-1^) under peaks. The percent crystallinity was calculated using the following equation:

$\% crystallinity= \left[ \frac{\Delta H_{m}-\Delta H_{c}}{\Delta H_{m}^{\circ}} \right]\times100$	(2)

where ΔH_m_° is the heat of fusion for a 100% crystalline polymer, which is estimated to be 140.1 J g^-1 16^, 103.4 J g^-1 5^ and 145.5 J g^-1 6^ for PET, PEN and PBT, respectively.

Energy Calculations

Based on the crystal structure of *Is*PETase (PDB ID: 5XH3^7^), energy calculations with ABACUS^8^, FoldX^9^ and Rosetta_ddg^10^ were performed. All positions of the protein sequence were mutated *in silico* to all proteinogenic amino acids except cysteine. The relative folding free energy changes (∆∆G^Fold^) predicted by the FoldX and Rosetta_ddg algorithms were calculated using Eq. 3 as follows:

$\Delta\Delta G^{Fold}=\Delta G_{mutation}^{Fold}-\Delta G_{WT}^{Fold}$	(3)

In Eq. 1, the ΔG^Fold^ represents the free energy difference between the folded and unfolded structures. For FoldX, standard settings were used, and each calculation was repeated five times to obtain better averaging. We used the settings described by Kellogg et al. ^10^ for Rosetta_ddg (options -ddg::local_opt_only true -ddg::opt_radius 8.0 -ddg::weight_file soft_rep_design -ddg::iterations 50 -ddg::min_cst false -ddg::mean true -ddg::min false -ddg::sc_min_only false -ddg::ramp_repulsive false). Any substitution with predicted ∆∆G^Fold^ < -5 kJ mol^-1^ was selected. For ABACUS, the automatic design protocol was used. The total ABACUS energy consists of the statistical energy function (SEF) and van der Waals energy terms. After calculation, the predicted mutations with relative energy changes less than -3 ABACUS energy units (A. e. u.) were retained as potentially stabilizing mutations for further investigation.

Consensus Analysis

For consensus analysis, an amino acid sequence alignment of 9 already-known PET hydrolases from thermophilic organisms (Figure S1) was performed by multiple sequence alignment. Sequences were acquired from databases integrated into the NCBI (<https://www.ncbi.nlm.nih.gov/>) and UniProt (http://www.uniprot.org/). An amino acid sequence HMM search was performed using the HMMER (http://hmmer.org/) webpage. Then, the alignment was used to construct a profile HMM with the “hmmbuild” function. For visualization, the HMM logo was created via the Skylign online tool^11^ (http://skylign.org/; Figure S1). Based on the relative entropy or Kullback-Leibler distance, the frequencies of each letter in the target position were calculated.

Clustering

K-means clustering is one of the most commonly used algorithms^12^, which uses k prototype vectors (i.e., centers or centroids of k clusters) to characterize data and minimizes a sum-of-squares cost function to find these prototypes with a coordinate descent optimization method. The algorithm uses Euclidean distance measure to compute distances between instances and clusters. Given a set of individuals (x_1_, x_2_, …, x_n_), k-means clustering aims to partition the individuals into k (≤ n) sets, which is known as the function given as follows:

$\arg min\sum_{i=1}^{k} \sum_{x\in S_{i}}^{k_{i}} \left\| x-\mu_{i} \right\|^{2}$	(4)

where $\left\| x-\mu_{i} \right\|$ is the Euclidean distance between *x*_i_ and *μ*_i_, *k*_i_ is the number of individuals in i^th^ cluster and *k* is the number of clusters. The variants were characterized by a set of parameters, including Δ*T*_M_ improvements, potential effects of the mutations, the interactions between the variants, the location of the variants and the distances between the Cα atoms of the variants and the catalytic triads. We implemented the WEKA program^13^ for discriminating individuals.

Molecular Docking and MD Simulations

A model substrate consisting of three consecutive units of PET, a head PET residue (like a N-terminal amino acid residue) and a tail PET residue (like a C-terminal amino acid residue) to cap the polymer chain in two ends was generated. Molecular docking was performed by YASARA. The ligand was bound to Subsite I and II by global docking. After global docking, the docking model with the highest binding energy and distance of PET carbonyl carbon and Ser160 Cα atom below 4 Å was selected. The complexes were then subjected to 999 runs of local docking, yielding the final docked binding mode.

The PETase-PET complex was simulated in AMBER 16^14^ using the ff14SB force field. The force field parameters of PET molecular refer to the Amber contributed parameters database for material systems. To keep all systems neutral, Cl^-^ ions were added based on a coulomb potential grid. Systems were then solvated with the TIP3P water model in a truncated octahedron box with a 10-Å distance around the solute (Jorgensen, Chandrasekhar, Madura, Impey, & Klein, 1983) and minimized over 5,000 steps of steepest descent minimization followed by 7,000 steps of conjugate gradient minimization. Subsequently, the systems were heated from 0 K to 310 K by Langevin dynamics with collision frequency 1 ps^-1^, equilibrated over 500 ps, and simulated for 100 ns with a time step of 2 fs. Short-range interactions were cut off at 12 Å, and bonds involving hydrogen were held fixed using SHAKE.

**Table S1.** Experimentally characterized point mutations of *Is*PETase predicted by ABACUS, FoldX, Rosetta_ddg and Consensus analysis. Protein variants that are significantly more thermostable (Δ*T*_m_ > 1.5 ºC) are labeled in blue.

| **ABACUS** | | | **FoldX** | | | **Rosetta_ddg** | | | **Consensus Analysis** | | |
| --- | --- | --- | --- | --- | --- | --- | --- | --- | --- | --- | --- |
| **Mutations** | **ABACUS Energy** | **Δ*T*_m_** | **Mutations** | **ΔΔ*G*_fold_ (kJ/mol)** | **Δ*T*_m_** | **Mutations** | **ΔΔ*G*_fold_ (kJ/mol)** | **Δ*T*_m_** | **Mutations** | **Consensus Score** | **Δ*T*_m_** |
| **N37P** | **-3.35** | **-0.5** | **A40P** | **-6.82** | **-0.5** | **N37P** | **-6.11** | **-0.5** | **A74T** | **2.45** | **-3.5** |
| **G66A** | **-3.94** | **-0.5** | **A47W** | **-6.78** | **-0.5** | **A47W** | **-7.33** | **-0.5** | **S121D** | **2.58** | **-1.5** |
| **A82V** | **-5.80** | **-6.5** | **T51P** | **-8.62** | **-1.5** | **S54W** | **-15.82** | **-1.5** | **A152S** | **2.00** | **0** |
| **Q91P** | **-3.28** | **-8.5** | **S54W** | **-11.09** | **-1.5** | **T77P** | **-9.23** | **-1** | **G155A** | **2.89** | **+0.5** |
| **K95A** | **-3.68** | **+2.5** | **T77E** | **-5.98** | **+1.5** | **R100F** | **-24.24** | **-3** | **W159H** | **4.57** | **+8.5** |
| **Q119Y** | **-3.84** | **+4.5** | **R90D** | **-6.40** | **-1** | **H104F** | **-5.05** | **-1.5** | **A180I** | **2.48** | **+1.5** |
| **R123V** | **-5.28** | **-6** | **S92P** | **-6.23** | **-2** | **D112R** | **-16.99** | **-3** | **Q182L** | **2.12** | **-2.5** |
| **T140D** | **-5.49** | **+2** | **R100F** | **-10.21** | **-3** | **Q119Y** | **-6.65** | **+4.5** | **A183T** | **2.22** | **-5** |
| **M154F** | **-3.80** | **0** | **H104F** | **-7.07** | **-1.5** | **S125R** | **-7.70** | **0** | **D186H** | **2.26** | **+7** |
| **W159H** | **-6.55** | **+8.5** | **T113P** | **-16.07** | **-3.5** | **Q133Y** | **-9.08** | **-3.5** | **T189K** | **3.26** | **-1.5** |
| **G165A** | **-3.09** | **+1.5** | **T116P** | **-9.04** | **+0.5** | **G139N** | **-10.90** | **+1** | **S214H** | **4.34** | **+9** |
| **S166T** | **-3.51** | **+2** | **L117F** | **-5.78** | **+3** | **K148W** | **-17.33** | **+2** | **I218F** | **3.72** | **-8.5** |
| **P181A** | **-3.90** | **+6** | **Q133Y** | **-5.27** | **-3.5** | **A152S** | **-5.39** | **0** | **S223P** | **2.17** | **-6.5** |
| **Q182L** | **-3.05** | **-2.5** | **V134L** | **-5.27** | **+1** | **I168R** | **-5.16** | **+7.5** | **S238F** | **3.70** | **+1.5** |
| **A183T** | **-3.93** | **-5** | **S136Q** | **-6.28** | **-0.5** | **S175W** | **-21.18** | **0** | **A240P** | **3.45** | **+0.5** |
| **I208V** | **-3.17** | **+1.5** | **S141P** | **-8.70** | **+1** | **D186W** | **-20.29** | **-1** | **K253Y** | **3.19** | **+0.5** |
| **N225V** | **-5.33** | **-6** | **G147T** | **-5.01** | **+1** | **S187W** | **-8.11** | **+3** | **T270Q** | **3.73** | **-2.5** |
| **Q228V** | **-4.10** | **-8.5** | **T151P** | **-5.61** | **+1** | **S193R** | **-6.45** | **-0.5** | **A272L** | **2.64** | **0** |
| **G254V** | **-7.93** | **-4** | **S175W** | **-6.53** | **0** | **T198V** | **-5.22** | **0** | **E274P** | **2.74** | **+1** |
| **R280A** | **-3.23** | **+1.5** | **D186H** | **-6.78** | **+7** | **E204K** | **-8.12** | **-0.5** | **S290P** | **2.74** | **+0.5** |
|  | **-** |  | **S187P** | **-7.53** | **-1.5** | **P210K** | **-55.50** | **-1.5** |  |  |  |
|  |  |  | **S188Q** | **-5.00** | **+1.5** | **P217L** | **-8.20** | **-0.5** |  |  |  |
|  |  |  | **S193R** | **-6.36** | **-0.5** | **E231R** | **-12.17** | **-4.5** |  |  |  |
|  |  |  | **T198V** | **-6.82** | **0** | **S238F** | **-8.79** | **+1.5** |  |  |  |
|  |  |  | **N212R** | **-5.89** | **+3** | **N246W** | **-16.13** | **-6** |  |  |  |
|  |  |  | **A215P** | **-8.12** | **-6** | **T286V** | **-5.56** | **-4.5** |  |  |  |
|  |  |  | **D220E** | **-5.39** | **-0.5** |  |  |  |  |  |  |
|  |  |  | **A226P** | **-8.66** | **-1** |  |  |  |  |  |  |
|  |  |  | **E231M** | **-15.86** | **+0.5** |  |  |  |  |  |  |
|  |  |  | **A240V** | **-5.90** | **+0.5** |  |  |  |  |  |  |
|  |  |  | **N241L** | **-12.39** | **-1.5** |  |  |  |  |  |  |
|  |  |  | **N246W** | **-7.45** | **-6** |  |  |  |  |  |  |
|  |  |  | **A248P** | **-7.03** | **+3** |  |  |  |  |  |  |
|  |  |  | **K259F** | **-7.87** | **-2** |  |  |  |  |  |  |
|  |  |  | **T266P** | **-9.46** | **-2.5** |  |  |  |  |  |  |
|  |  |  | **T279P** | **-5.36** | **+0.5** |  |  |  |  |  |  |
|  |  |  | **R280A** | **-5.41** | **+1.5** |  |  |  |  |  |  |

**Table S2.** Primers used for cloning and site-directed mutagenesis

| Mutation | Primer (5'→3') | |
| --- | --- | --- |
| T77E | Forward | CAACCAATGCAGGCGGCGAGGTTGGCGCGATTGCAAT |
|  | Reverse | ATTGCAATCGCGCCAACCTCGCCGCCTGCATTGGTTG |
| K95A | Forward | AAGCAGCATTGCGTGGTGGGGTCCG |
|  | Reverse | CGGACCCCACCACGCAATGCTGCTT |
| L117F | Forward | GAACAGCACTTTTGACCAGCCCAGCA |
|  | Reverse | TGCTGGGCTGGTCAAAAGTGCTGTTC |
| Q119Y | Forward | CAGCACTCTAGACTACCCCAGCAGCC |
|  | Reverse | GGCTGCTGGGGTAGTCTAGAGTGCTG |
| T140D | Forward | GCGAGCTTGAACGGGGACAGCAGTAGCCCGAT |
|  | Reverse | ATCGGGCTACTGCTGTCCCCGTTCAAGCTCGC |
| K148W | Forward | GCAGTAGCCCGATTTACGGATGGGTCGATACTGCCC |
|  | Reverse | GGGCAGTATCGACCCATCCGTAAATCGGGCTACTGC |
| W159H | Forward | CATGGGTGTGATGGGCCATTCAATGGGGGGCGGCG |
|  | Reverse | CGCCGCCCCCCATTGAATGGCCCATCACACCCATG |
| G165A | Forward | ATGGGGGGCGGCGCTTCACTTCGTAG |
|  | Reverse | CTACGAAGTGAAGCGCCGCCCCCCAT |
| I168R | Forward | GCGGTTCACTTCGTAGCGCCGCGAAC |
|  | Reverse | GCGGTTCACTTCGTAGCGCCGCGAAC |
| A180I | Forward | GAGTTTAAAAGCAGCGATACCGCAGGCGCCATGG |
|  | Reverse | CCATGGCGCCTGCGGTATCGCTGCTTTTAAACTC |
| P181A | Forward | AAGCAGCGGCAGCGCAGGCGCCATGG |
|  | Reverse | CCATGGCGCCTGCGCTGCCGCTGCTT |
| D186H | Forward | CGCAGGCGCCATGGCACTCTTCAACCAAC |
|  | Reverse | GTTGGTTGAAGAGTGCCATGGCGCCTGCG |
| S188Q | Forward | CCGCAGGCGCCATGGGACTCTCAGACCAACTTCAGCAG |
|  | Reverse | CTGCTGAAGTTGGTCTGAGAGTCCCATGGCGCCTGCGG |
| I208V | Forward | GAGAATGATAGCGTTGCACCGGTGAA |
|  | Reverse | TTCACCGGTGCAACGCTATCATTCTC |
| N212R | Forward | GATAGCATTGCACCGGTGCGCAGCCATGCGC |
|  | Reverse | GCGCATGGCTGCGCACCGGTGCAATGCTATC |
| S238F | Forward | CGGCGGTAGCCACTTTTGTGCCAACTCTG |
|  | Reverse | CAGAGTTGGCACAAAAGTGGCTACCGCCG |
| A248P | Forward | AACAGCAACCAGCCACTGATCGGAAAA |
|  | Reverse | TTTTCCGATCAGTGGCTGGTTGCTGTT |
| R280A | Forward | CCCAACAGCACAGCCGTGTCGGATTT |
|  | Reverse | AAATCCGACACGGCTGTGCTGTTGGG |
| L117F-Q119Y | Forward | GAACAGCACTTTCGACTACCCCAGCA |
|  | Reverse | TGCTGGGGTAGTCGAAAGTGCTGTTC |
| G165A-S166T | Forward | GGGGCGGCGCTACACTTCGTAGCGCC |
|  | Reverse | GGCGCTACGAAGTGTAGCGCCGCCCC |
| S166T-I168R | Forward | GGGGCGGCGGTACACTTCGTAGCGCC |
|  | Reverse | GGCGCTACGAAGTGTACCGCCGCCCC |
| A180I-P181A | Forward | AAAAGCAGCGATTGCGCAGGCGCCAT |
|  | Reverse | ATGGCGCCTGCGCAATCGCTGCTTTT |
| D186H-S188Q | Forward | CGCAGGCGCCATGGCACTCTCAGACCAAC |
|  | Reverse | GTTGGTCTGAGAGTGCCATGGCGCCTGCG |
| S187W-S188Q | Forward | GGCGCCATGGGACTGGTCACAAAACT |
|  | Reverse | AGTTTTGTGACCAGTCCCATGGCGCC |

**Table S3.** Crystallographic data collection and refinement statistics of DuraPETase

| Data collection |  |
| --- | --- |
| Space group | *P2_1_2_1_2* |
| *a, b, c* [Å] | 87.55, 129.79, 51.76 |
| *α* /*β* /*γ* (°) | 90.00/90.00/90.00 |
| Resolution (Å) | 25-1.63  (1.69-1.63) |
| Unique reflections | 71809 (7155) |
| Redundancy | 9.0 (7.1) |
| Completeness (%) | 96.6 (97.0) |
| Average I/σ (I) | 24.4 (3.48) |
| CC 1/2* | (0.900) |
| Rmeans | 0.067 (0.566) |
| Rpim | 0.021 (0.208) |
| Refinement |  |
| No. of reflections | 71722 (7100) |
| R_work_ (95% data) | 0.168 (0.218) |
| R_free_ (5% data) | 0.182 (0.255) |
| Rmsd bonds (Å) | 0.006 |
| Rmsd angles (°) | 0.863 |
| Most favored (%) | 98.3 |
| Allowed (%) | 1.7 |
| Disallowed (%) | 0.0 |
| No. of non-H atoms / average B [Å^2^] |  |
| Protein | 3906/23.76 |
| Water | 519/35.84 |
| Ligand/Ion | 85/49.03 |
| PDB ID code | 6KY5 |

Values in parentheses are for the outermost resolution shells.

CC 1/2*= percentage of correlation between intensities from random half-datasets^1^.

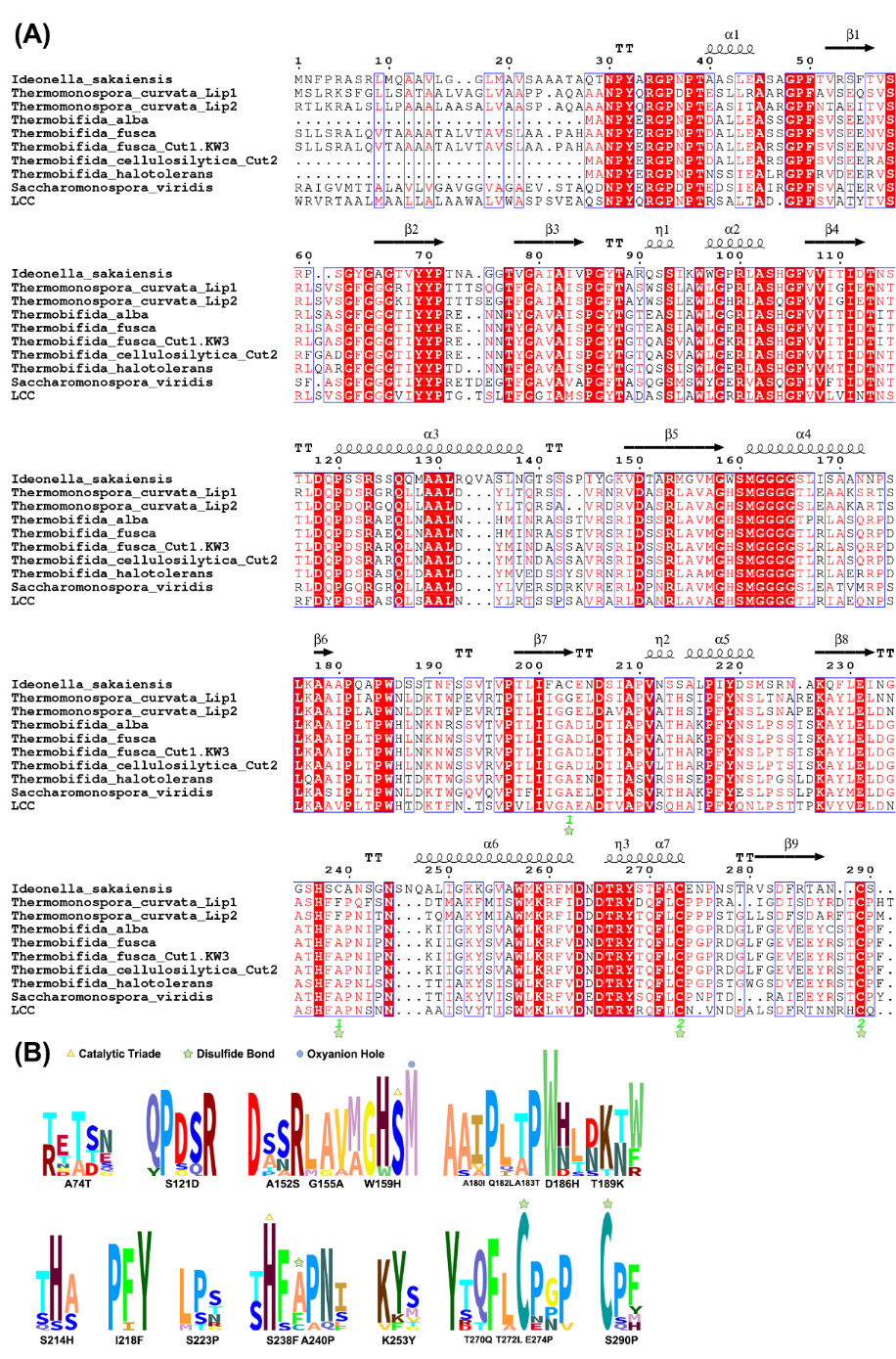

**Figure S1.** Amino acid sequence alignment of PET hydrolases. (A) An alignment of the PET hydrolase sequences used in the Consensus analysis. Aligned sequences with accession numbers are Ideonella_sakaiensis (*Is*PETase, *Ideonella sakaiensis*, A0A0K8P6T7), Thermomonospora_curvata_Lip1 (lipase, *Thermomonospora curvata*, WP_012851645), Thermomonospora_curvata_Lip2 (lipase, *Thermomonospora curvata*, WP_012850775), Thermobifida_alba (cutinase, *Thermobifida alba*, ADV92525), Thermobifida_fusca (lipase, *Thermobifida fusca*, WP_011291330), Thermobifida_fusca_Cut1.KW3 (cutinase, *Thermobifida fusca*, CBY05529), Thermobifida_cellulosilytica_Cut2 (cutinase 2, *Thermobifida cellulosilytica*, ADV92527), Thermobifida_halotolerans (serine hydrolase, *Thermobifida halotolerans*, AFA45122), Saccharomonospora_viridis (lipase, *Saccharomonospora viridis*, WP_015787089), LCC (cutinase, uncultured bacteria isolated from metagenomics analysis, AEV21261). (B) Hidden Markov model (HMM) of mutations with consensus score > 2.0. The amino acid alignment from panel A was used to calculate an HMM profile. The HMM was consequently visualized as a logo with information content above the background (Skylign; http://skylign.org).

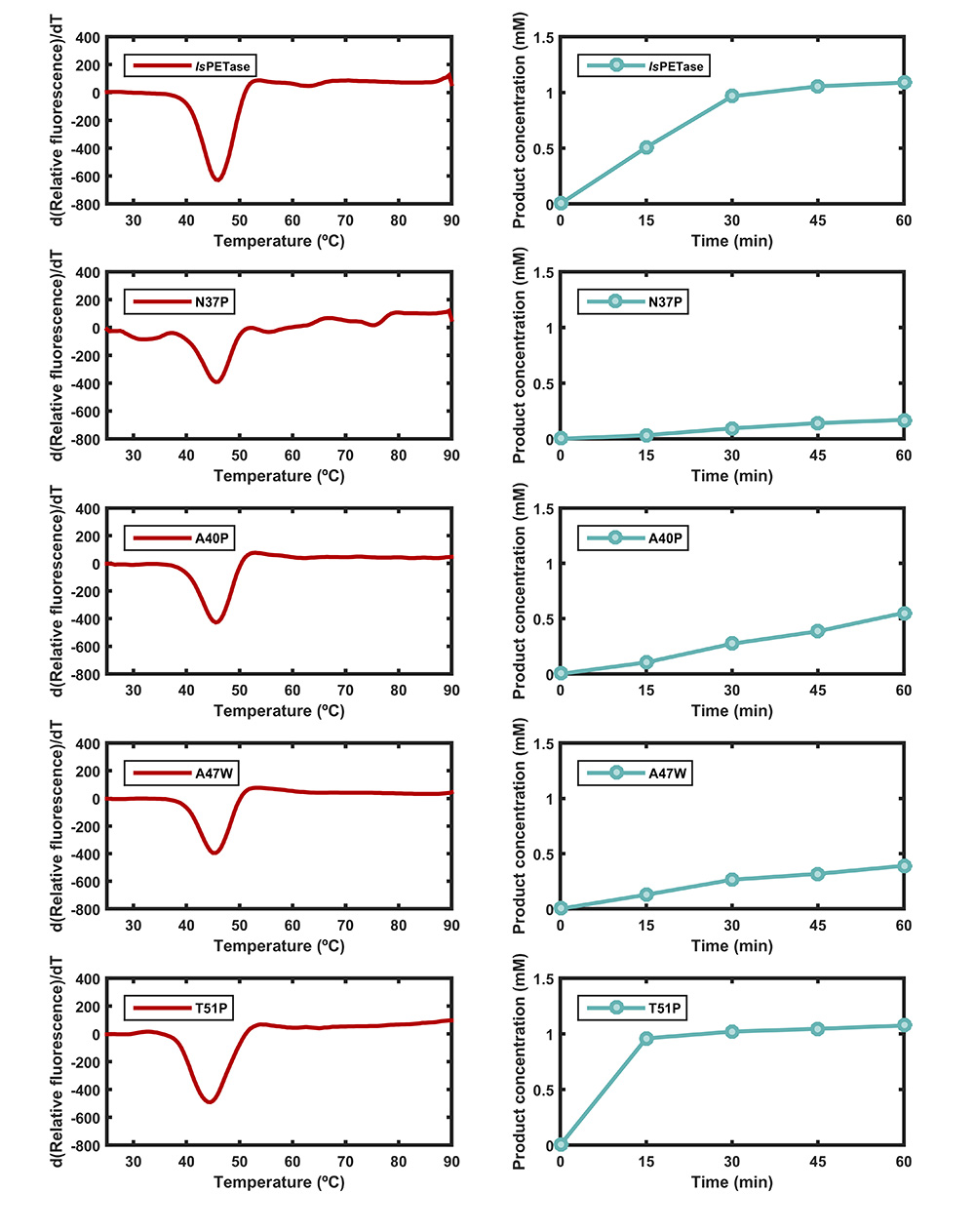

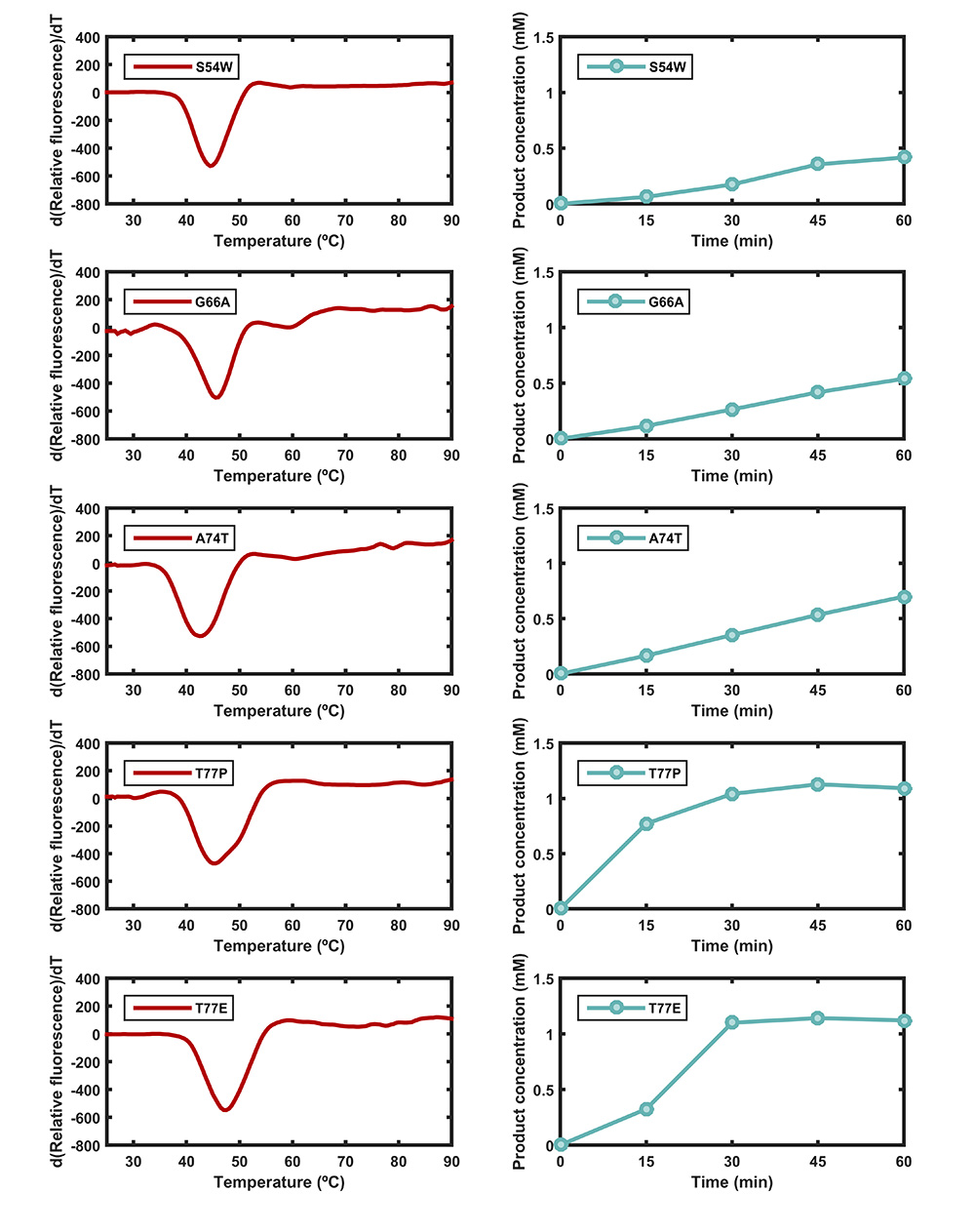

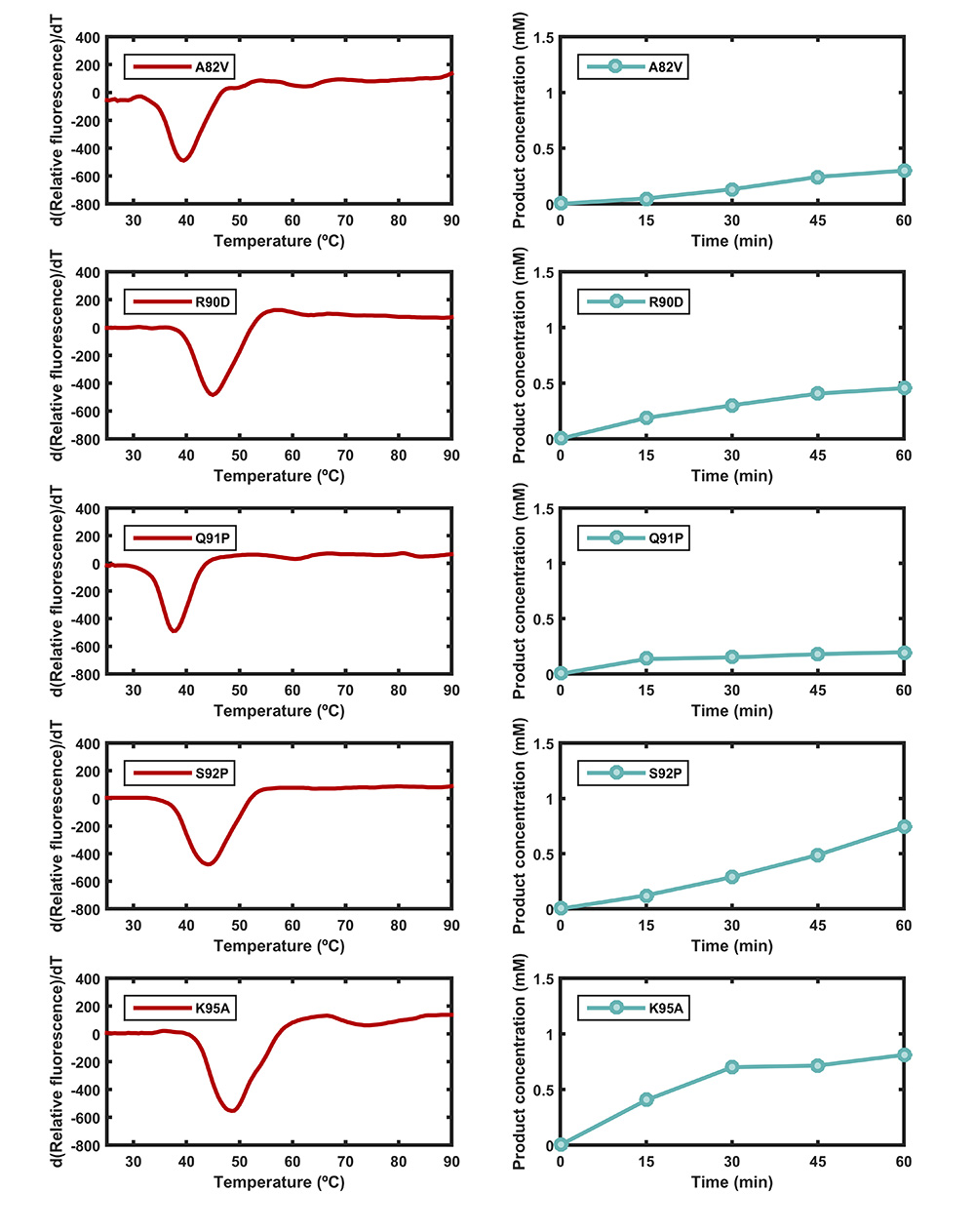

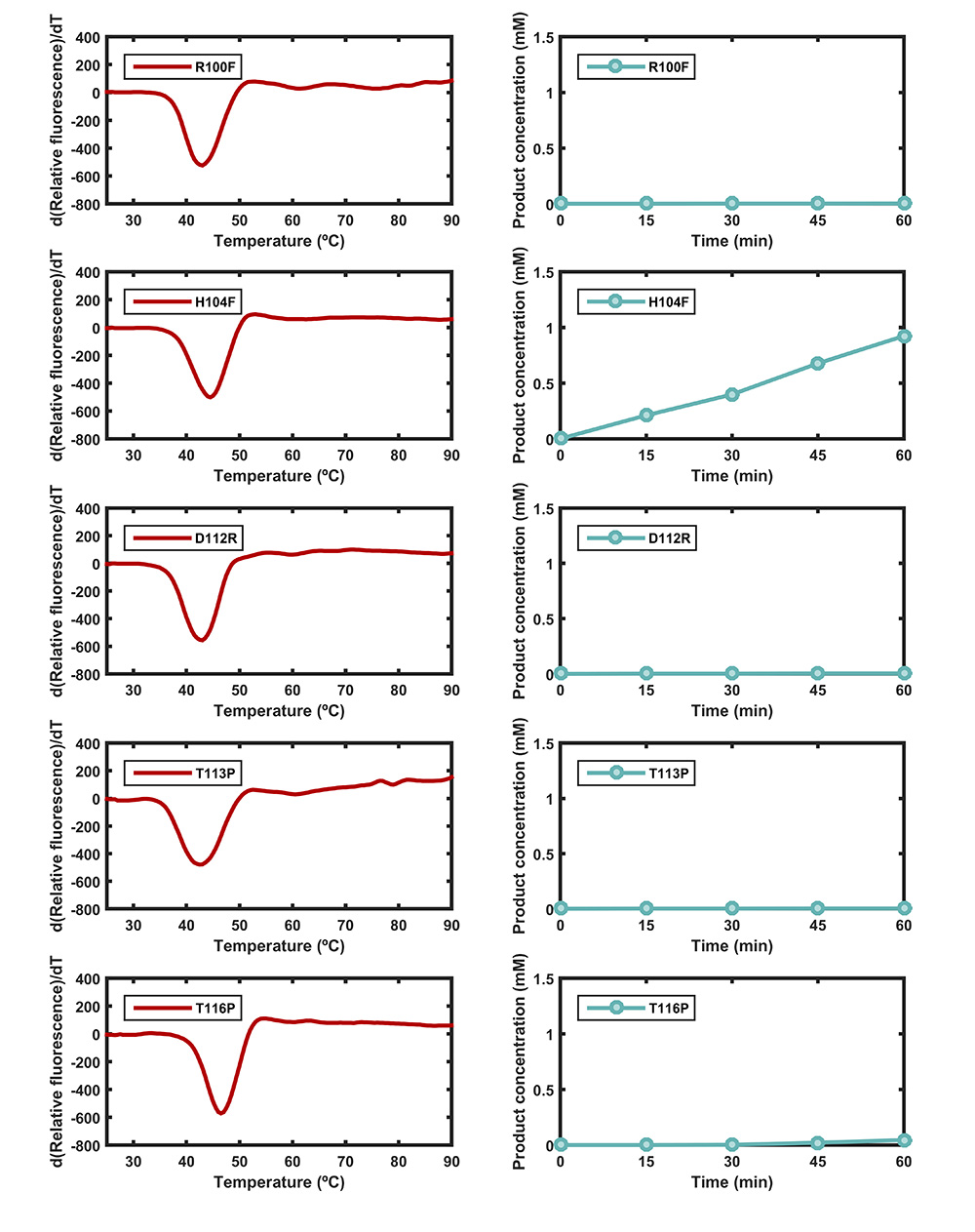

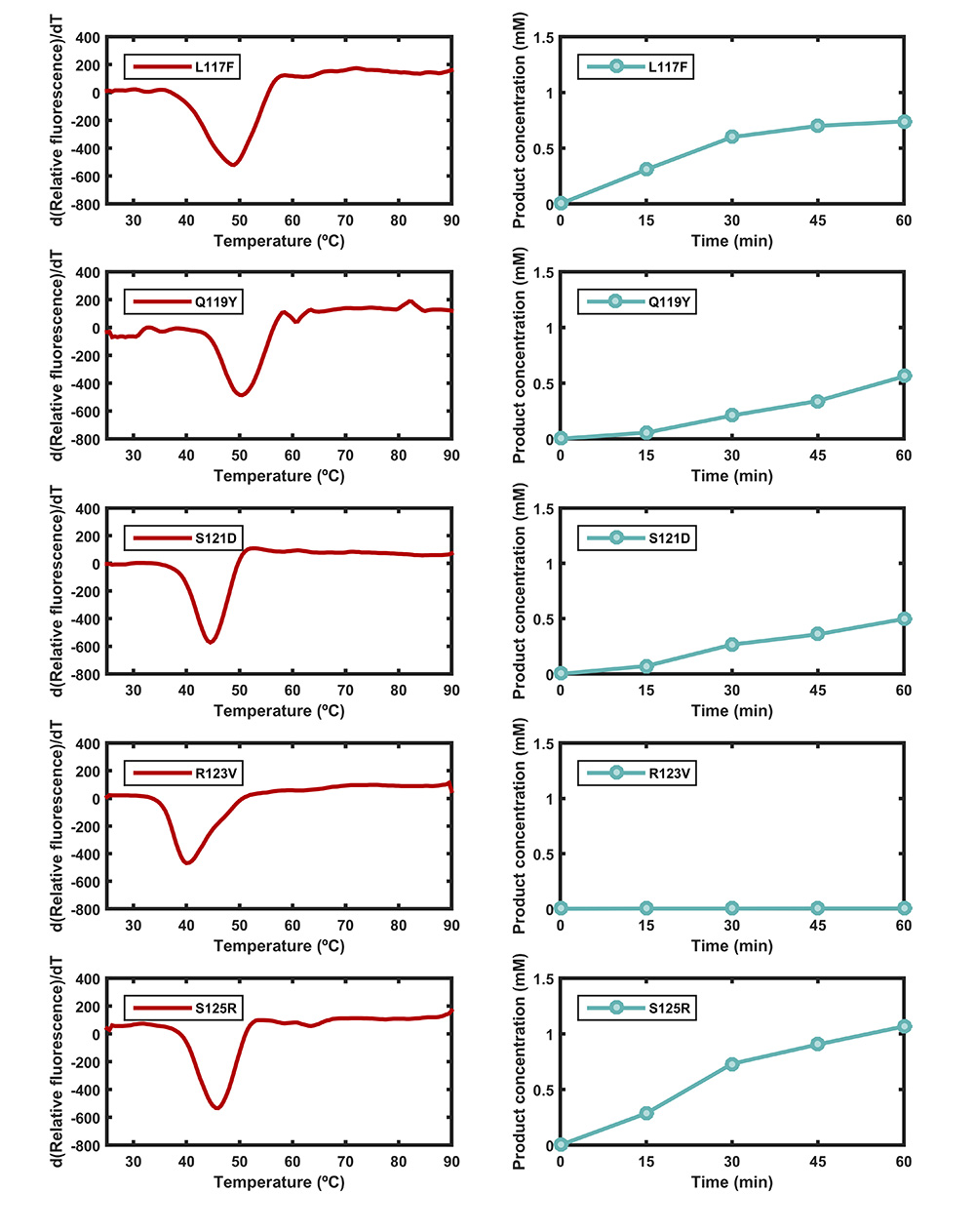

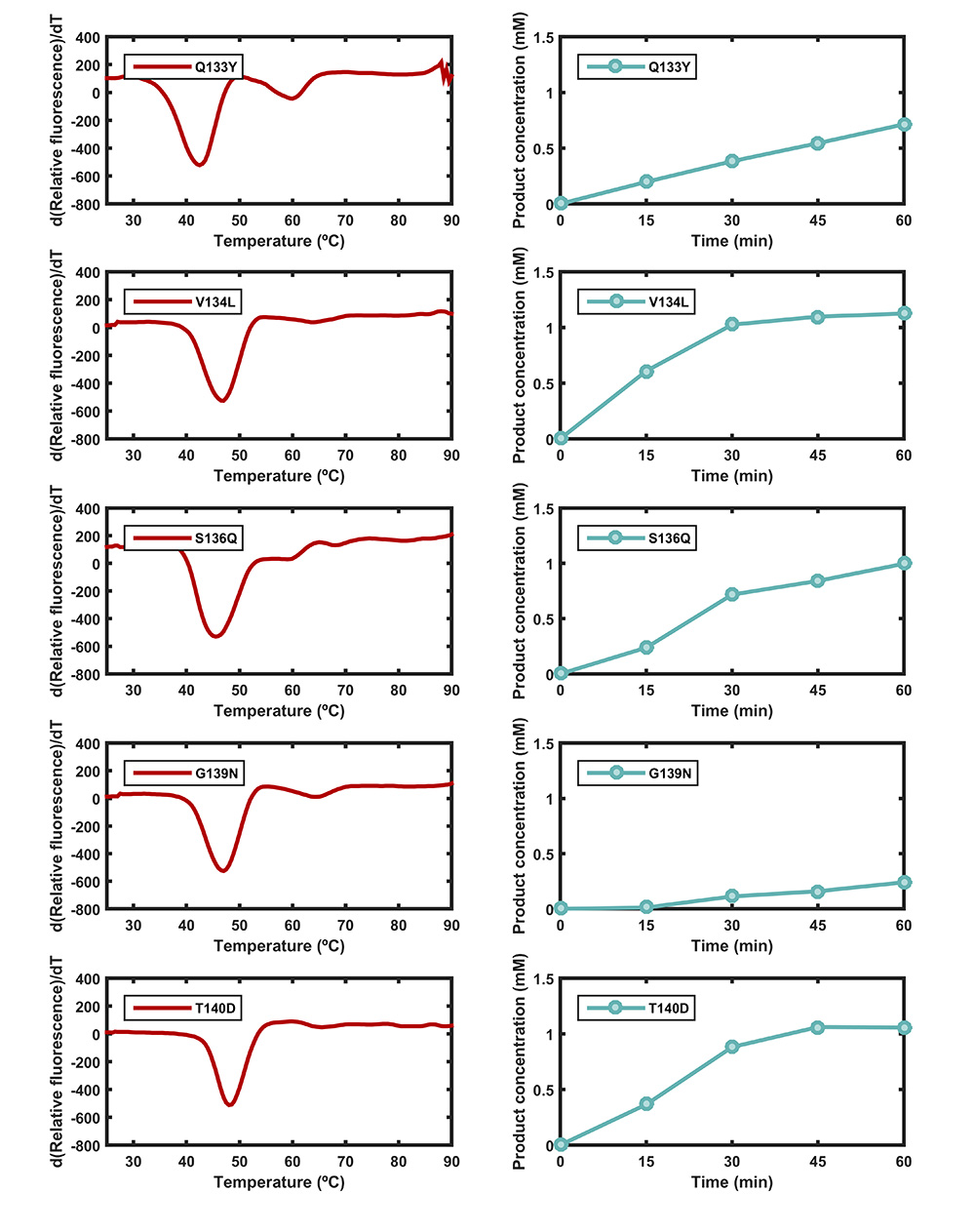

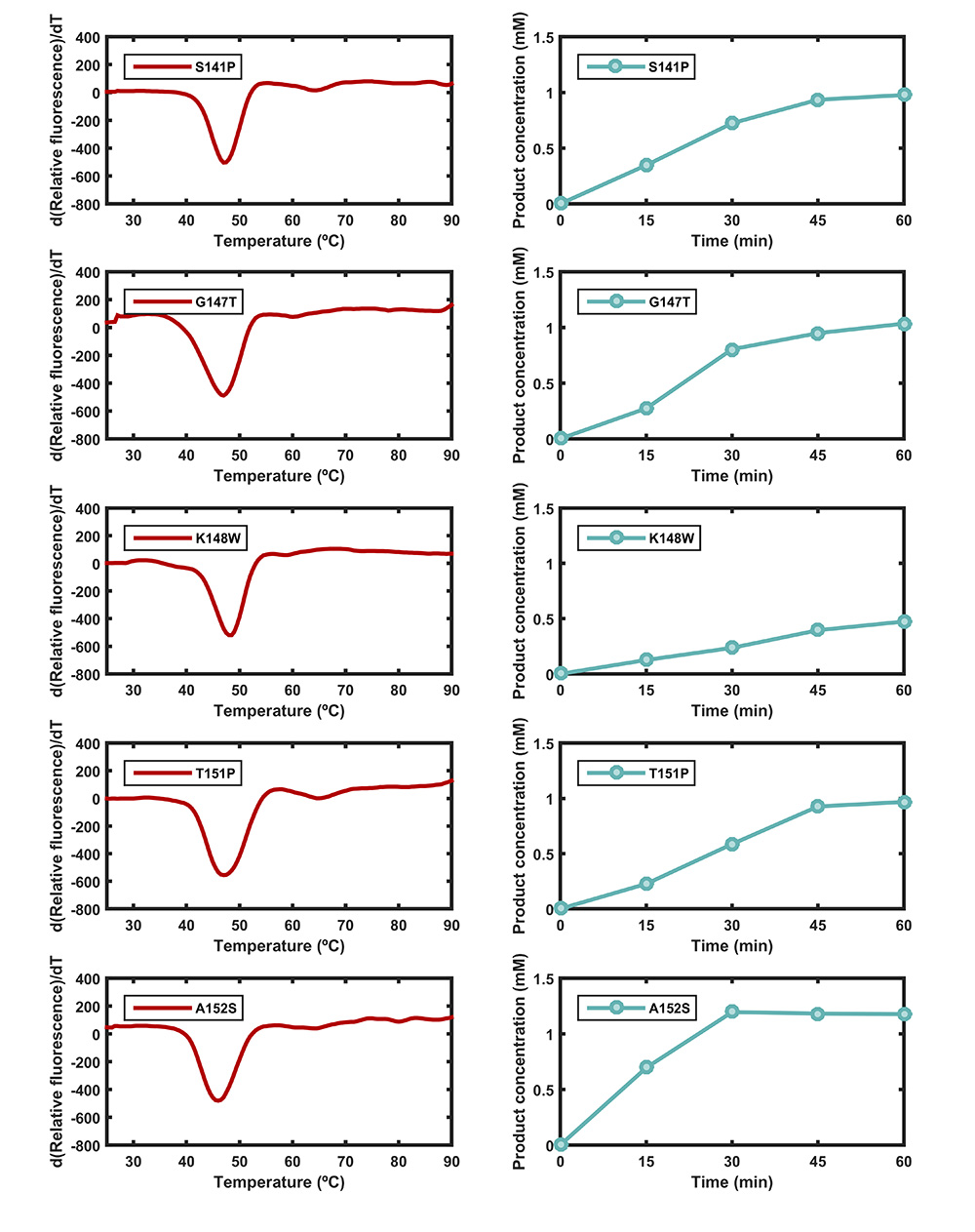

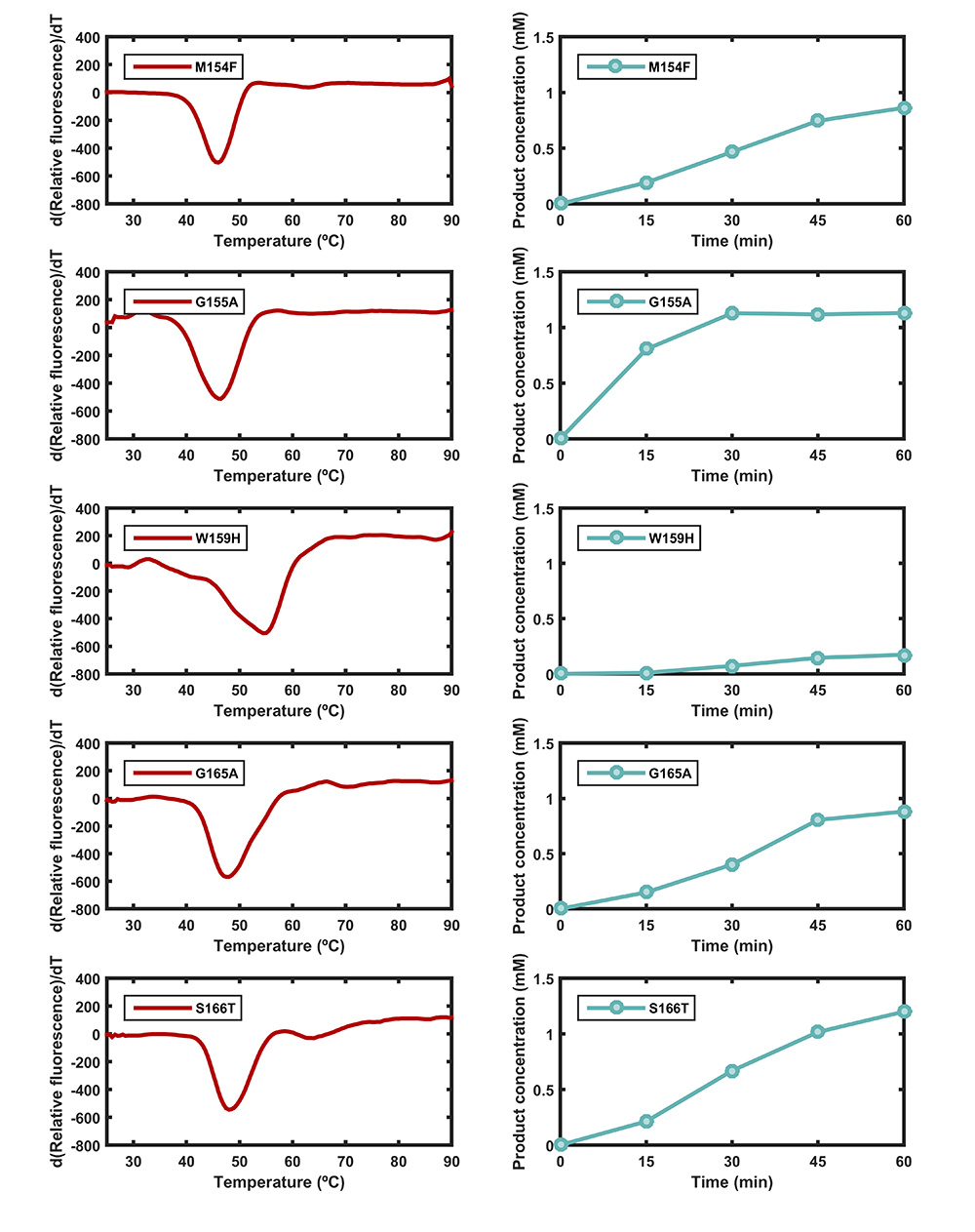

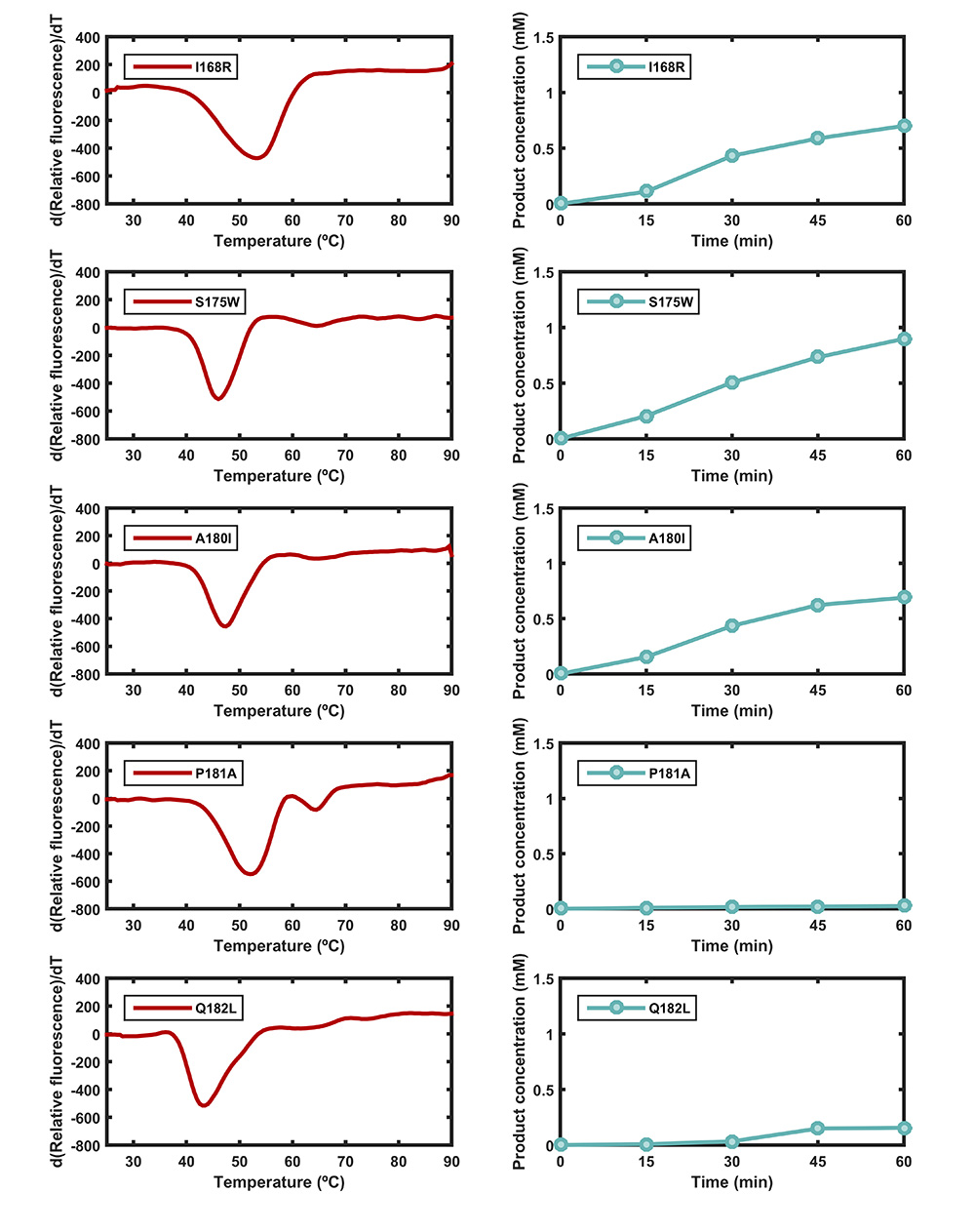

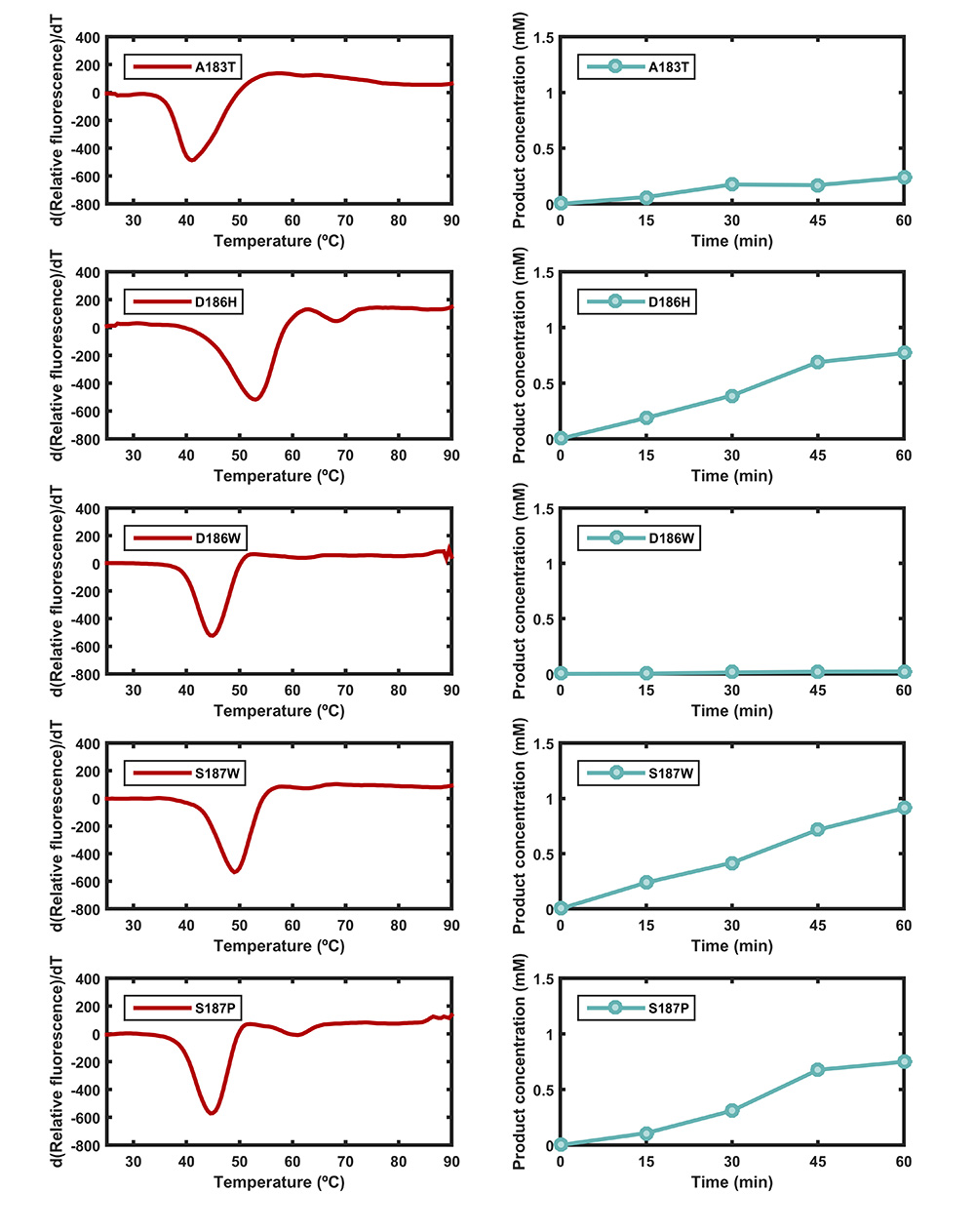

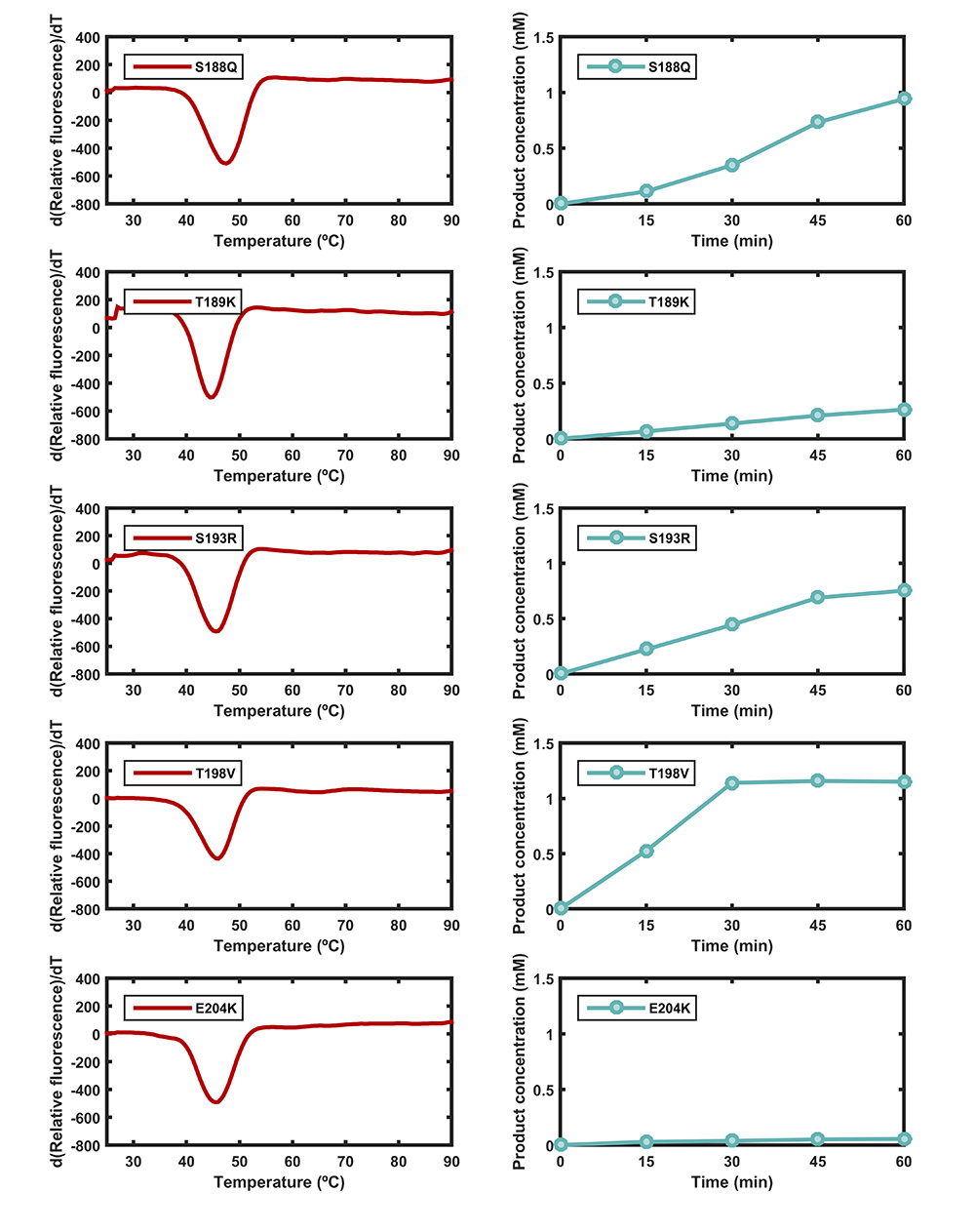

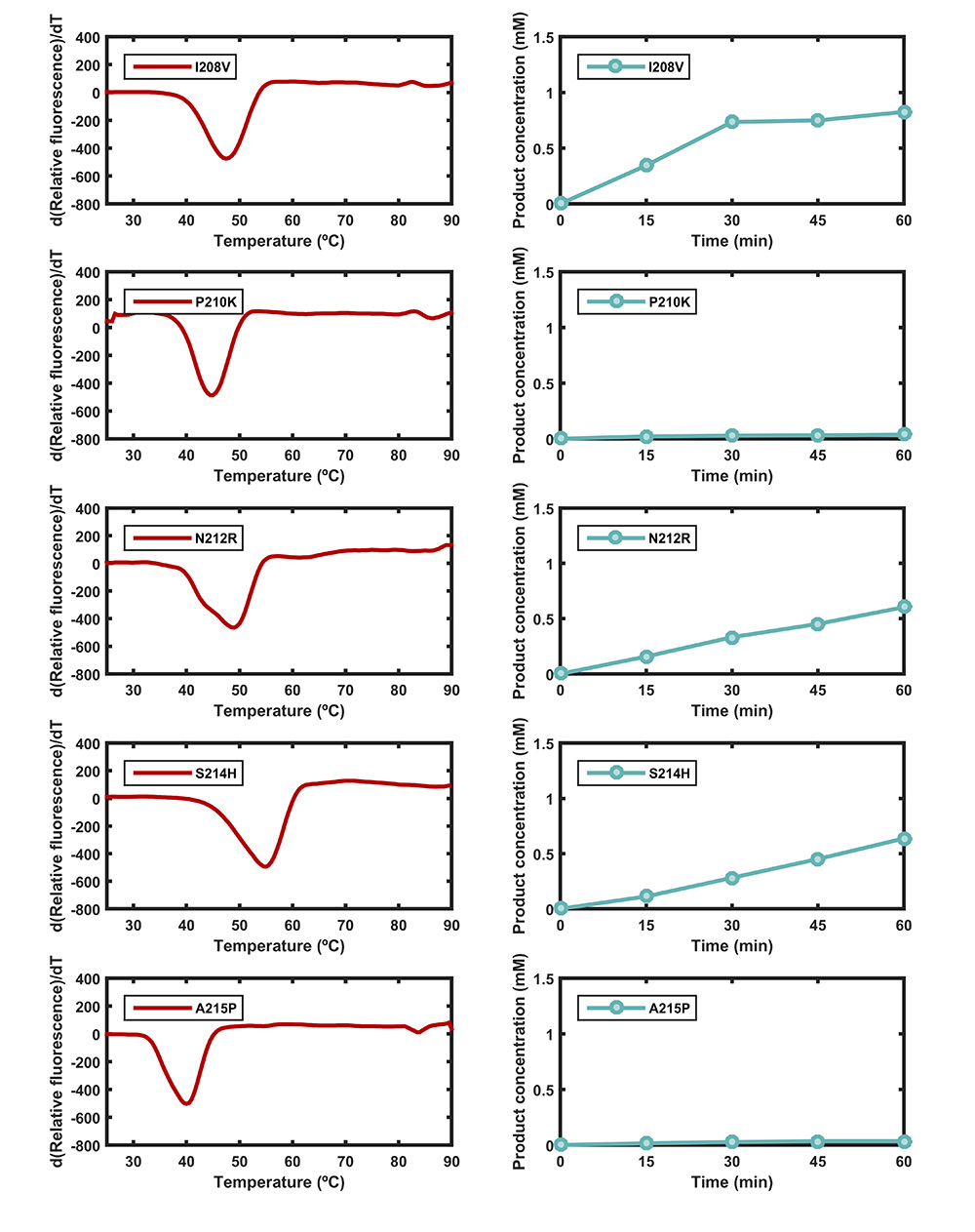

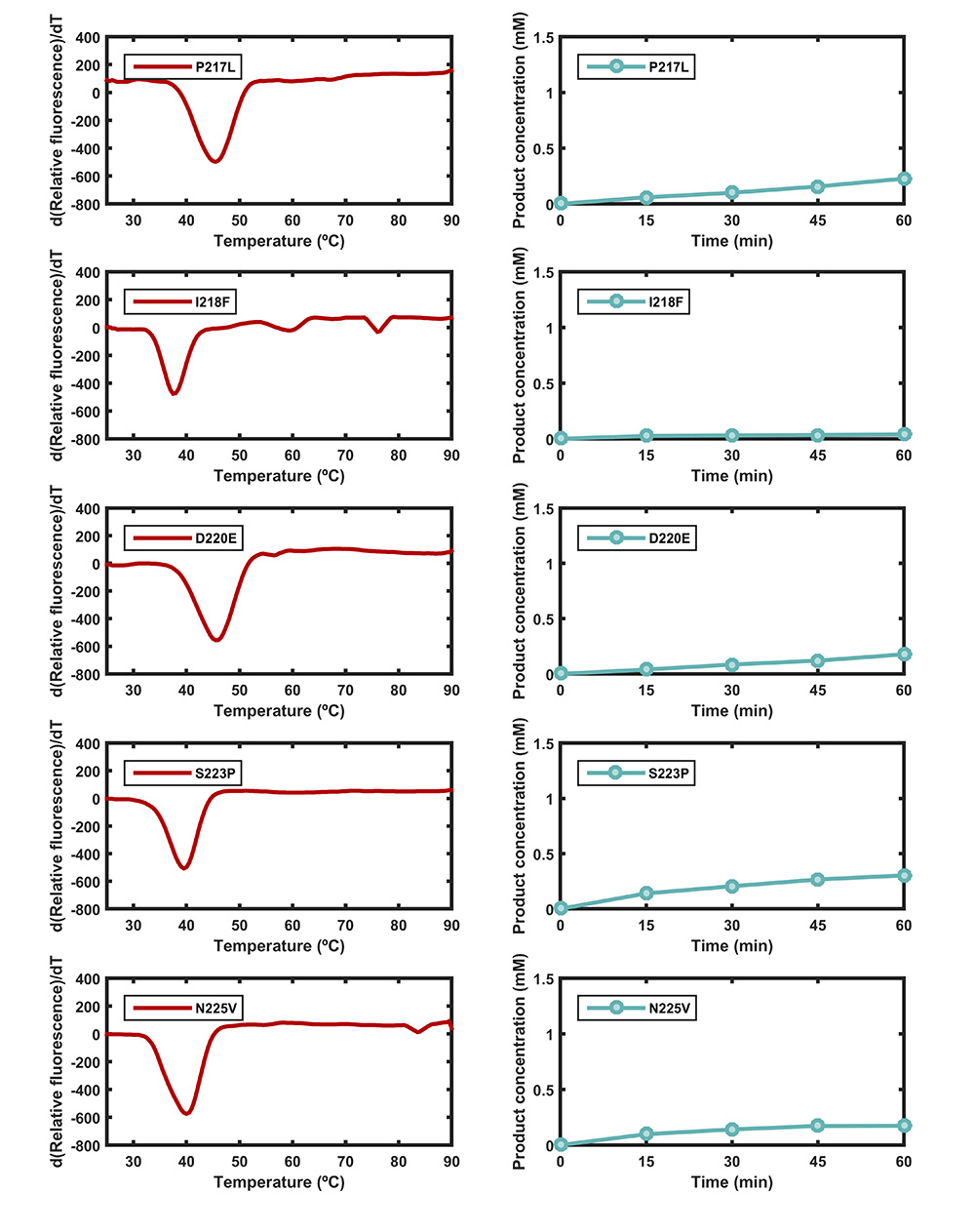

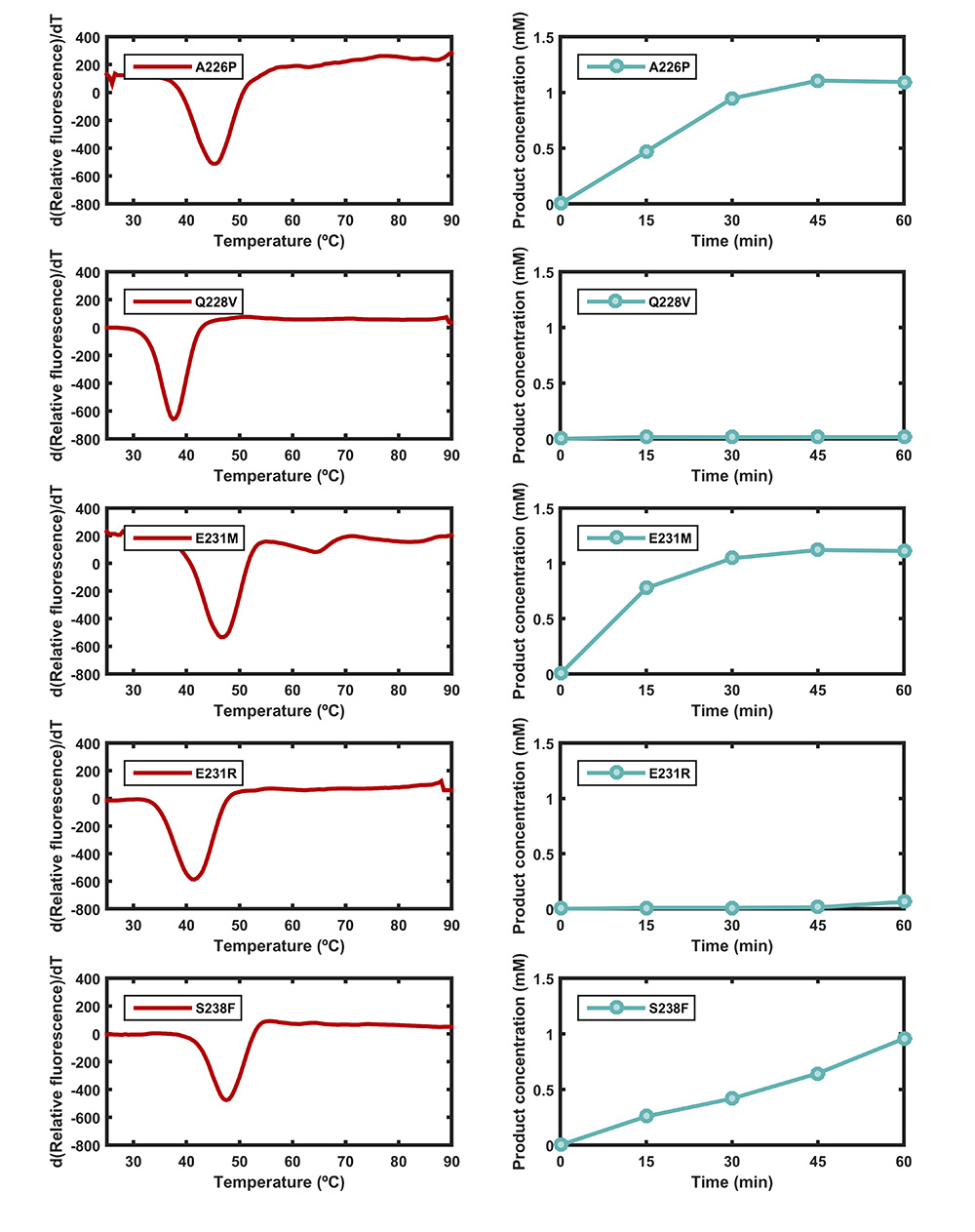

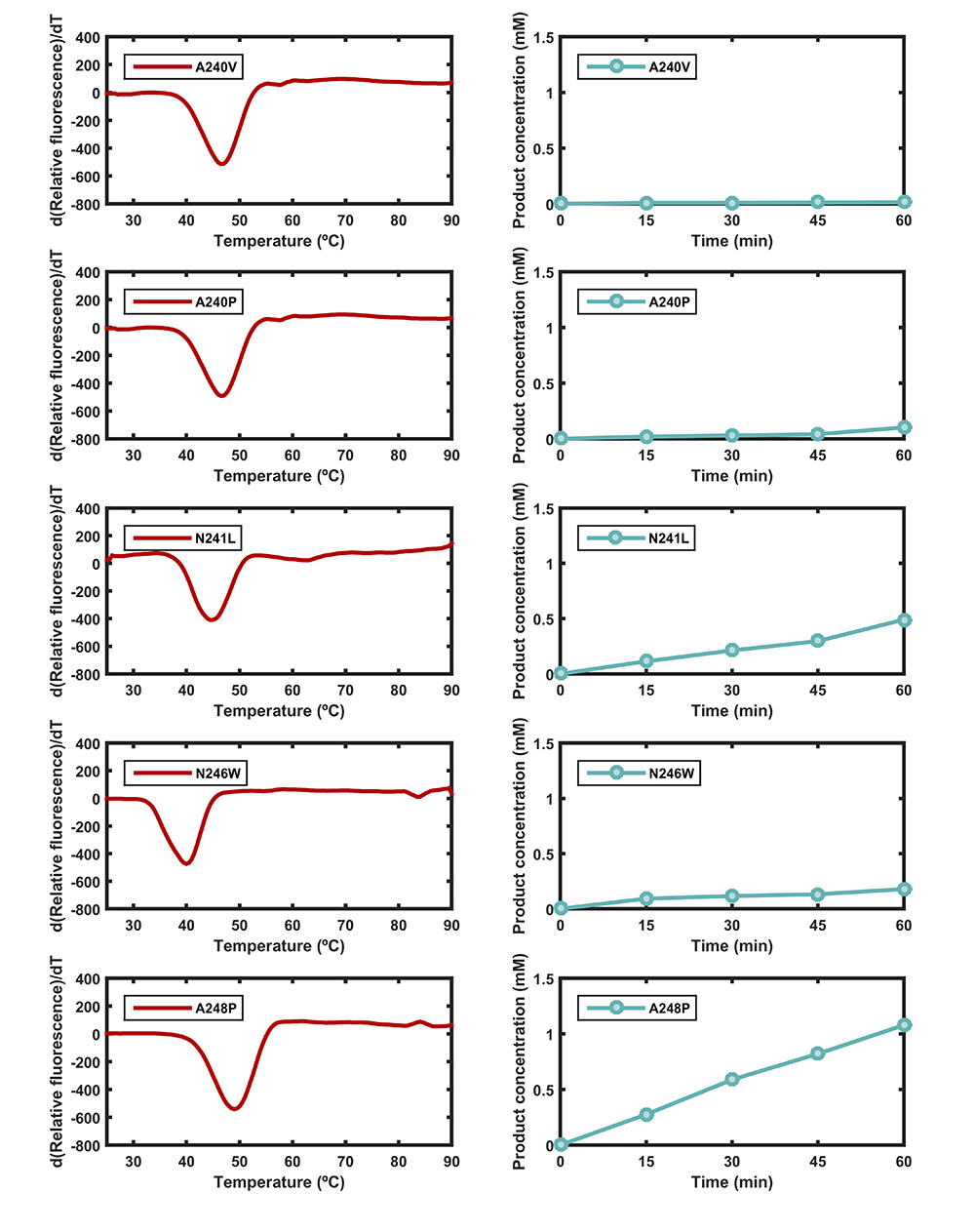

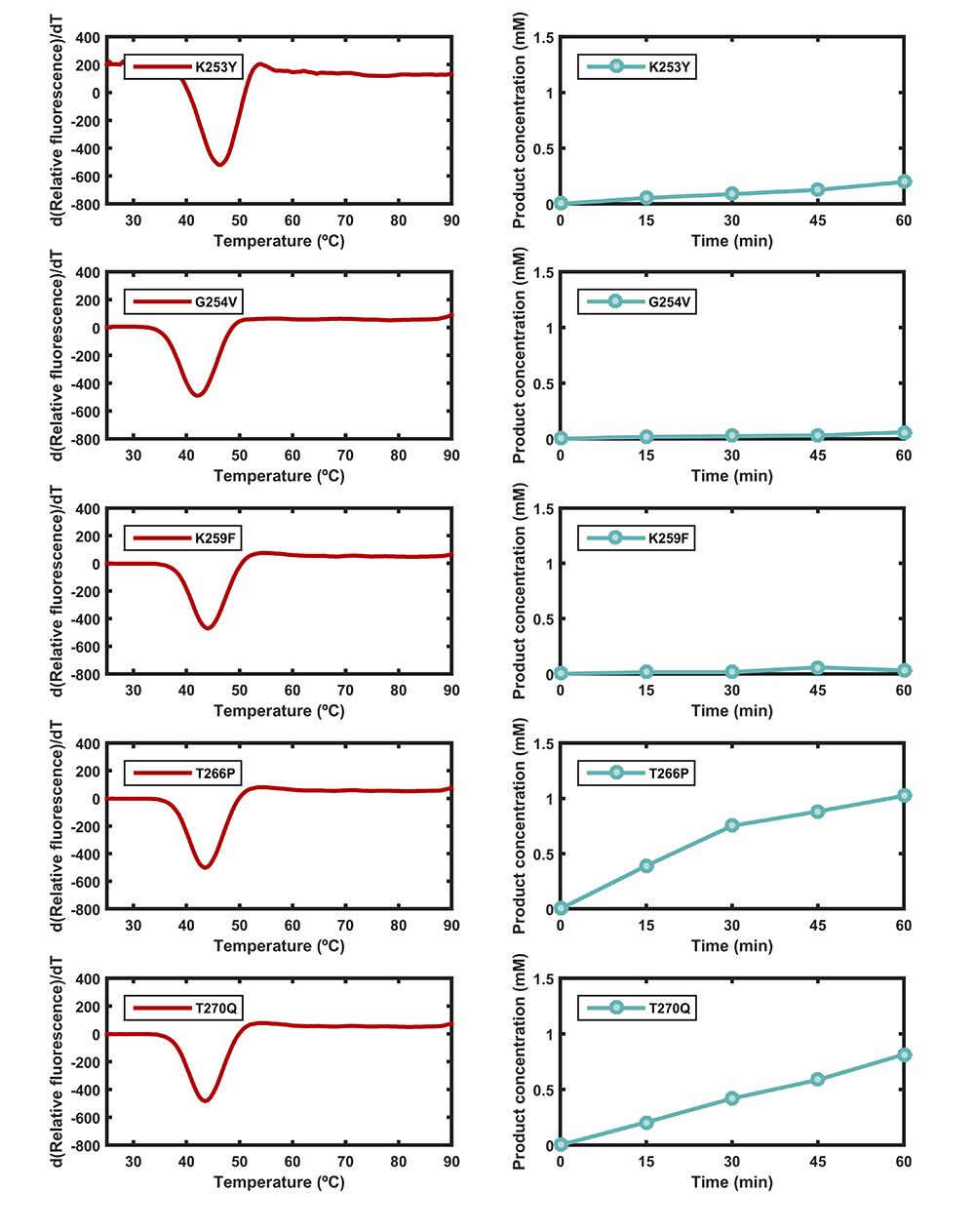

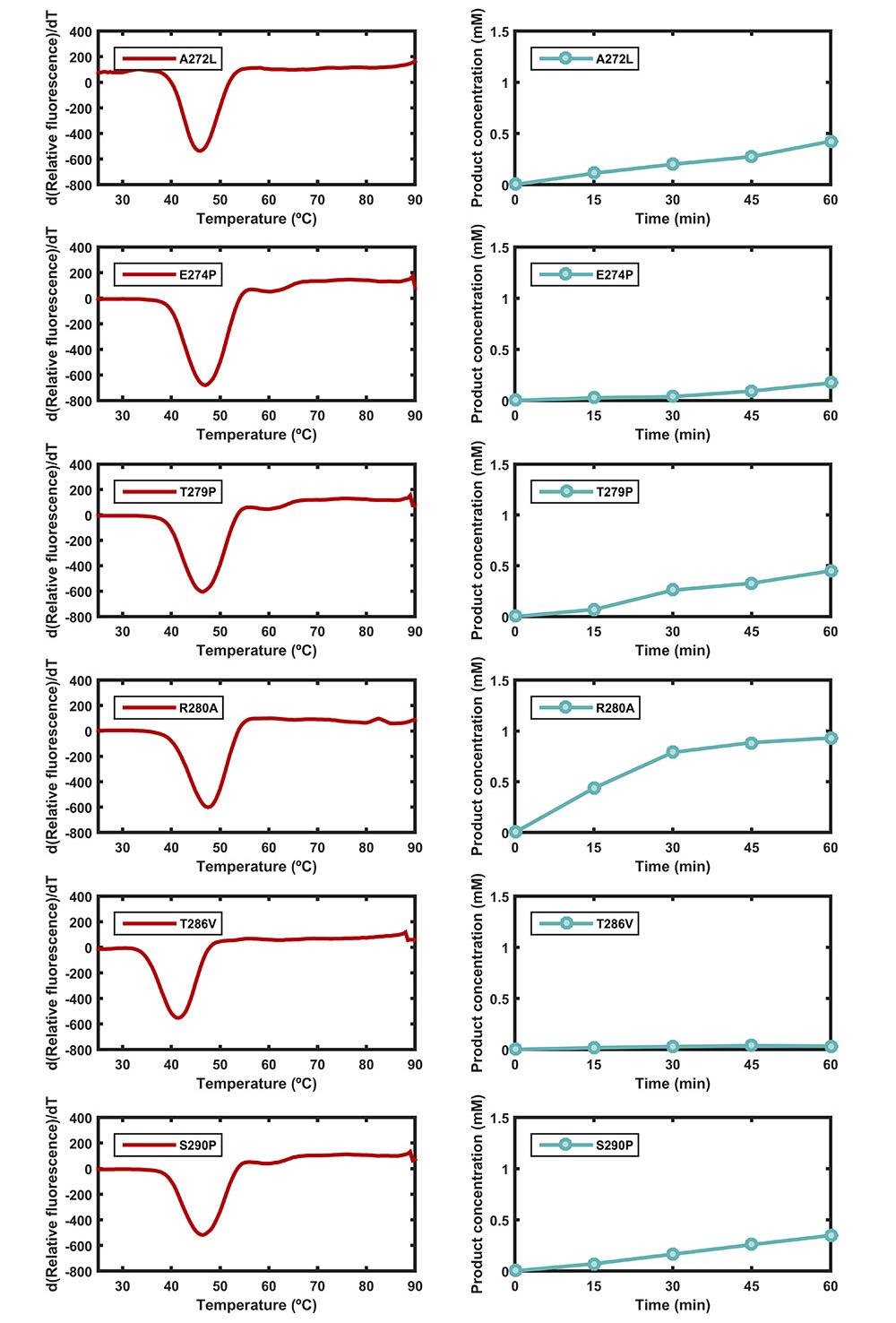
 **Figure S2.** Left panel: Apparent melting temperatures (*T*_M_) of *Is*PETase and 85 selected variants measured by the thermofluor method. Right panel: PET degradation activity of *Is*PETase and 85 selected variants. PET nanoparticles (200 µL) were incubated with 10 μL of enzyme (stock concentration 0.1 mg/mL) in 290 μL of 50 mM glycine-NaOH buffer (pH 9.0) at 37 ºC for 1 h.

|  | Apparent melting temperature (ºC) |
| --- | --- |
| IsPETase | 46 |
| T77E | 47.5 |
| K95A | 48.5 |
| L117F | 49 |
| Q119Y | 50.5 |
| T140D | 48 |
| K148W | 48 |
| W159H | 54.5 |
| G165A | 47.5 |
| S166T | 48 |
| I168R | 53.5 |
| A180I | 47.5 |
| P181A | 52 |
| D186H | 53 |
| S187W | 49 |
| S188Q | 47.5 |
| I208V | 47.5 |
| N212R | 49 |
| S214H | 55 |
| S238F | 47.5 |
| A248P | 49 |
| R280A | 47.5 |

**Figure S3.** *T*_M_ values of *Is*PETase and 21 beneficial variants

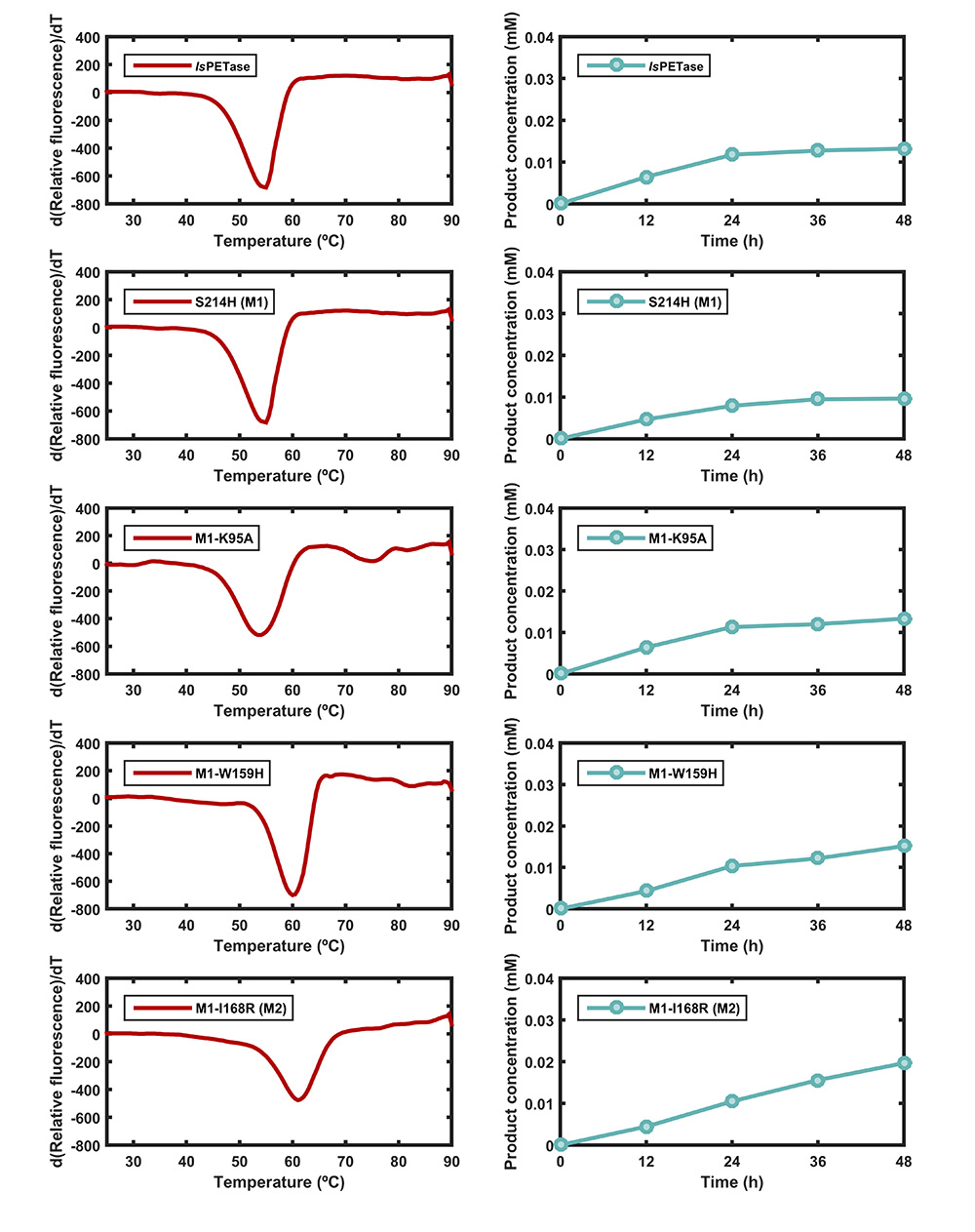

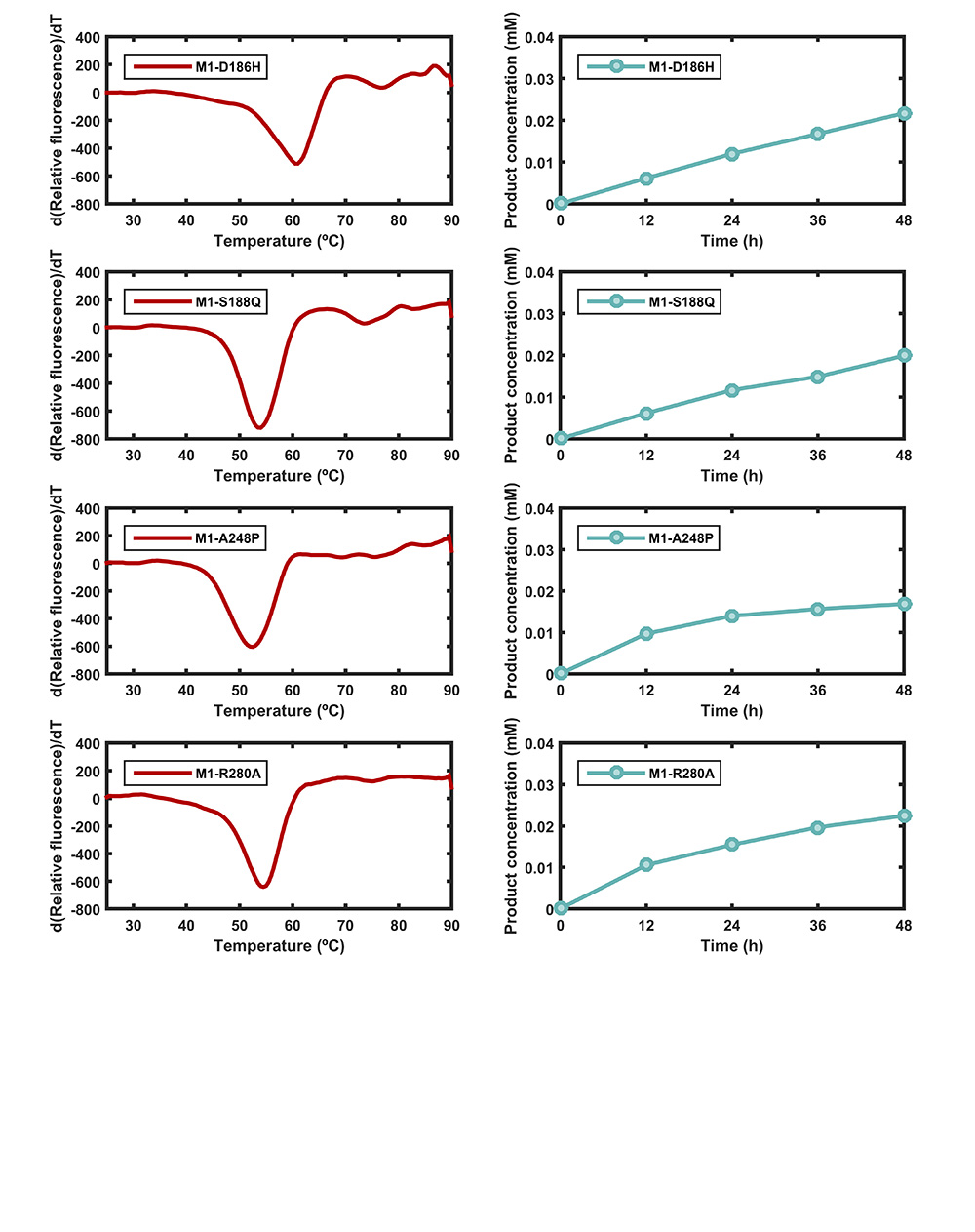

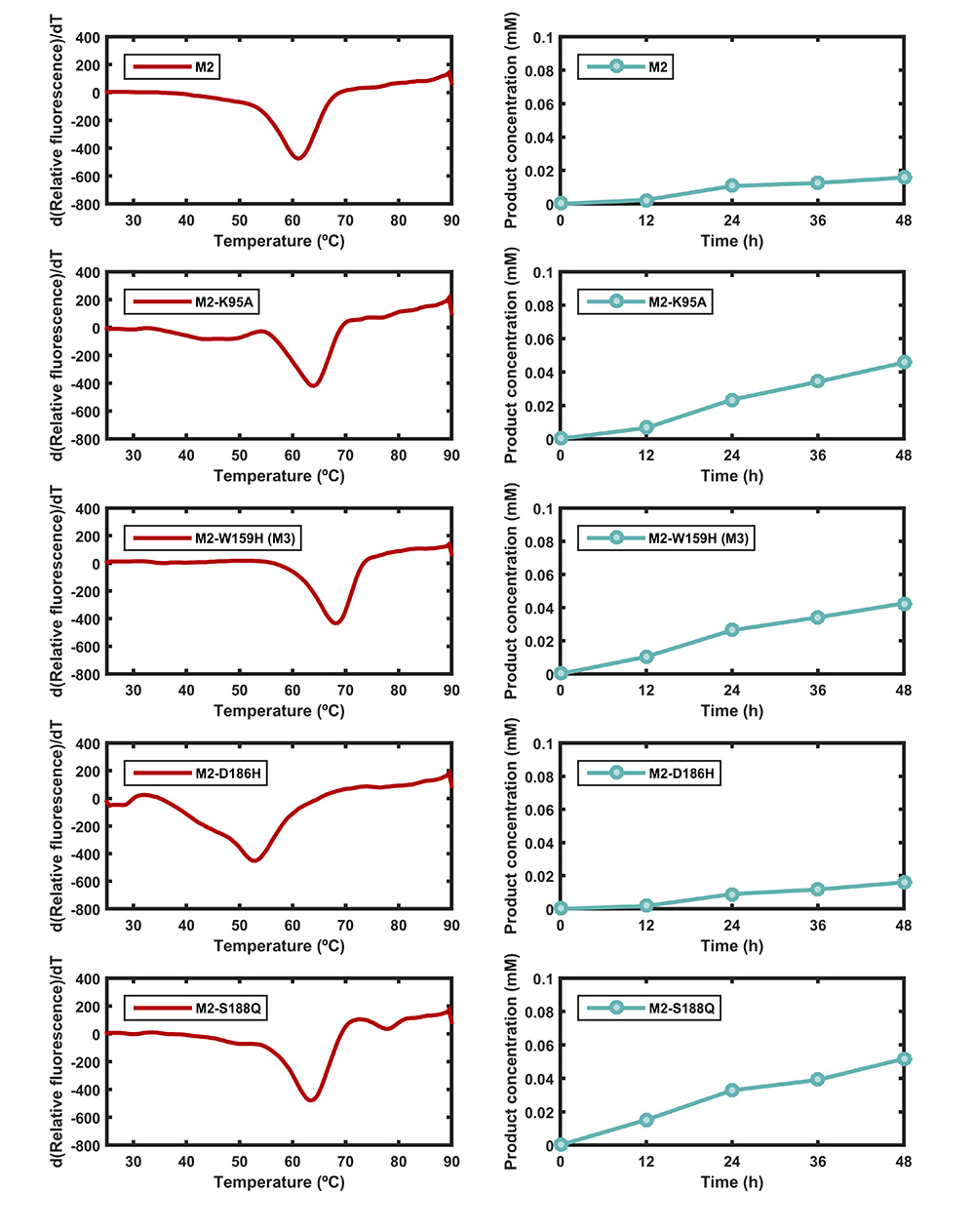

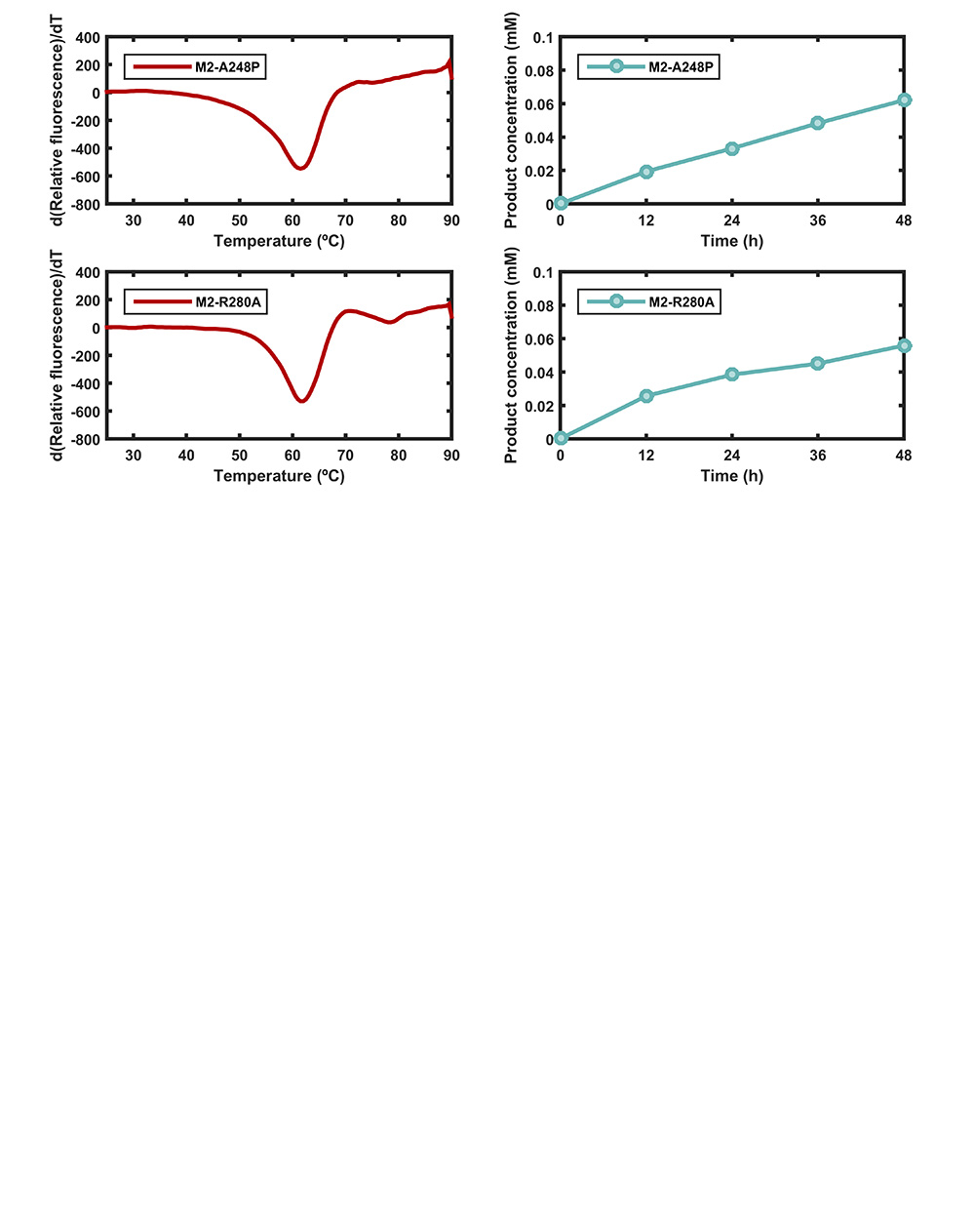

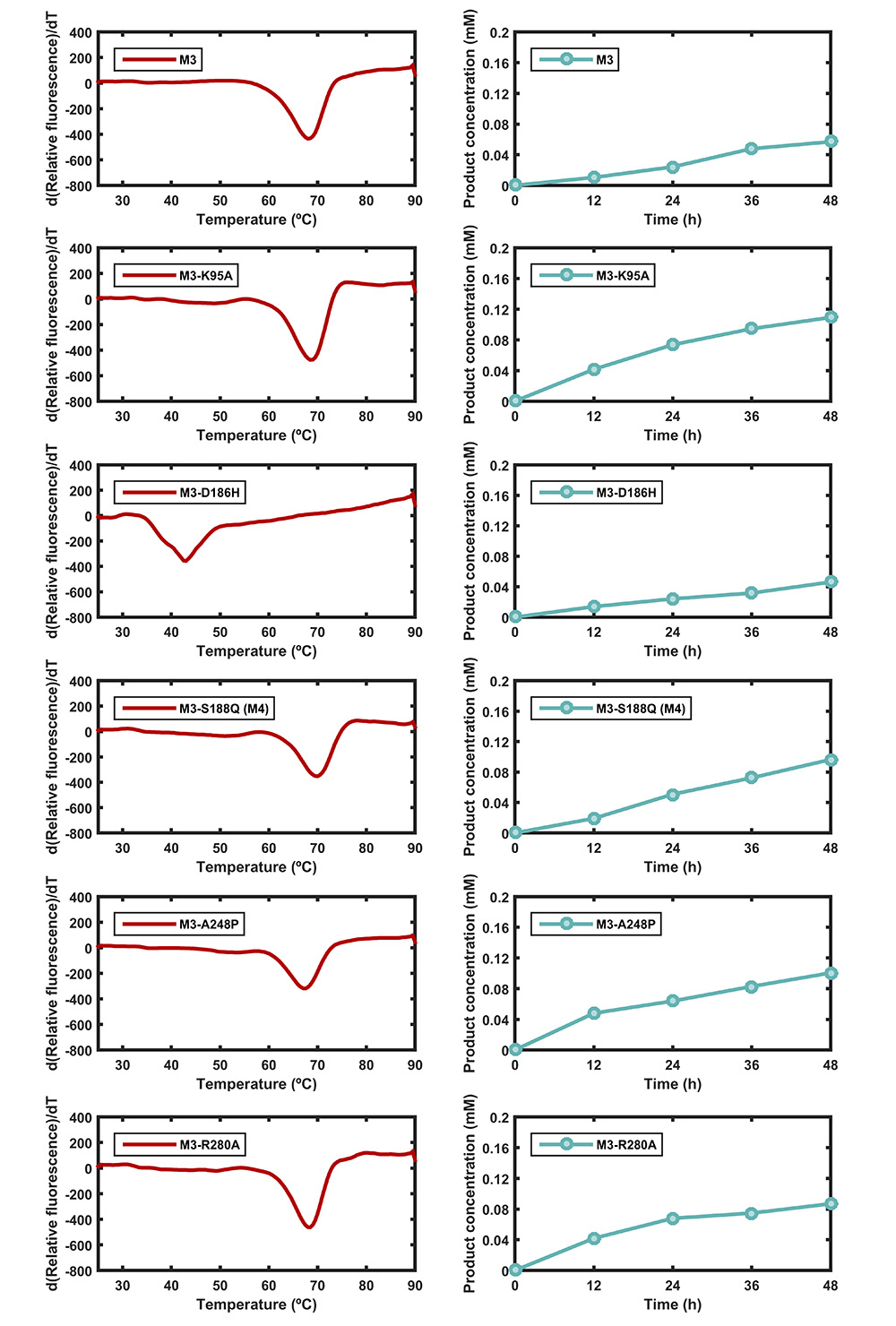

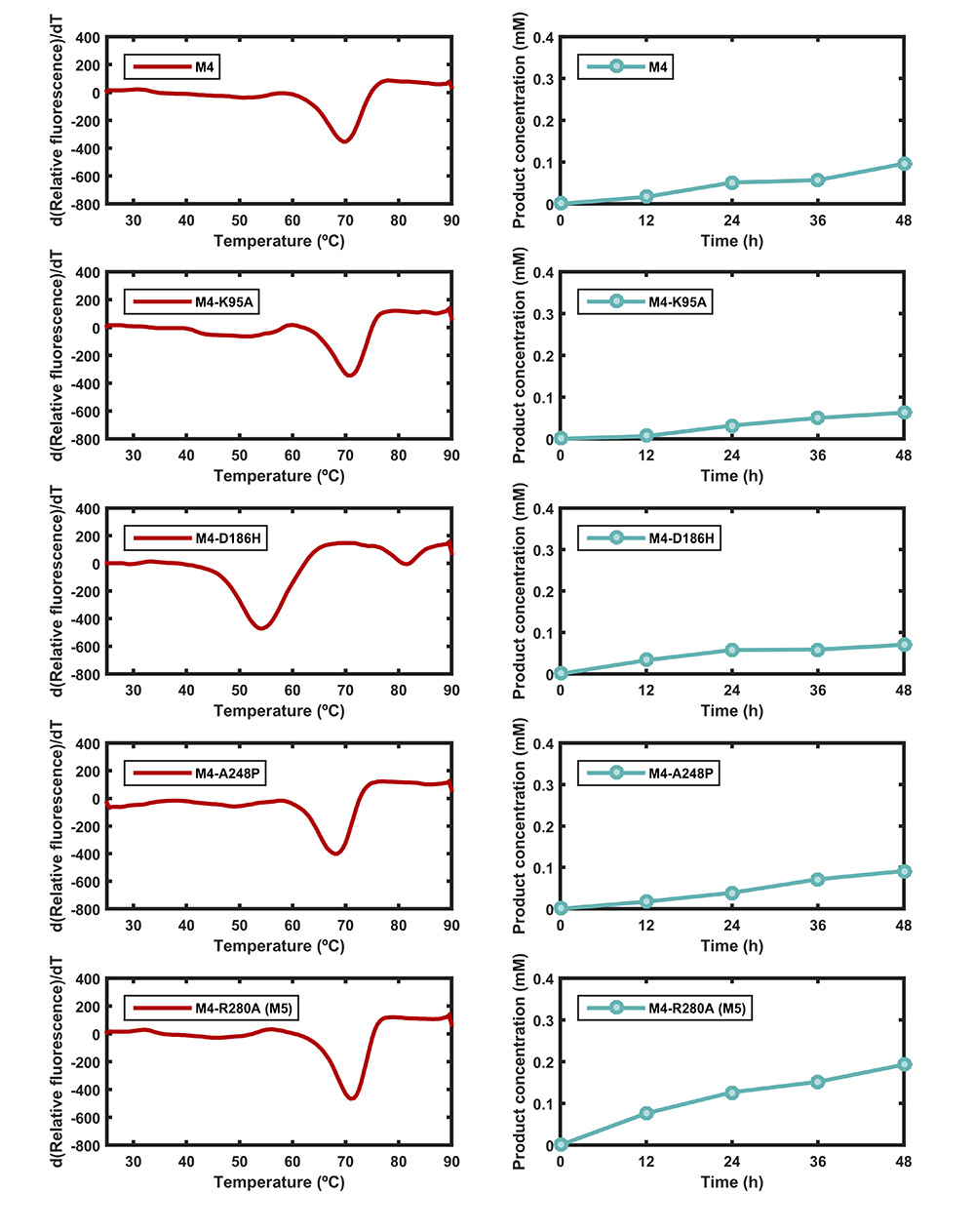

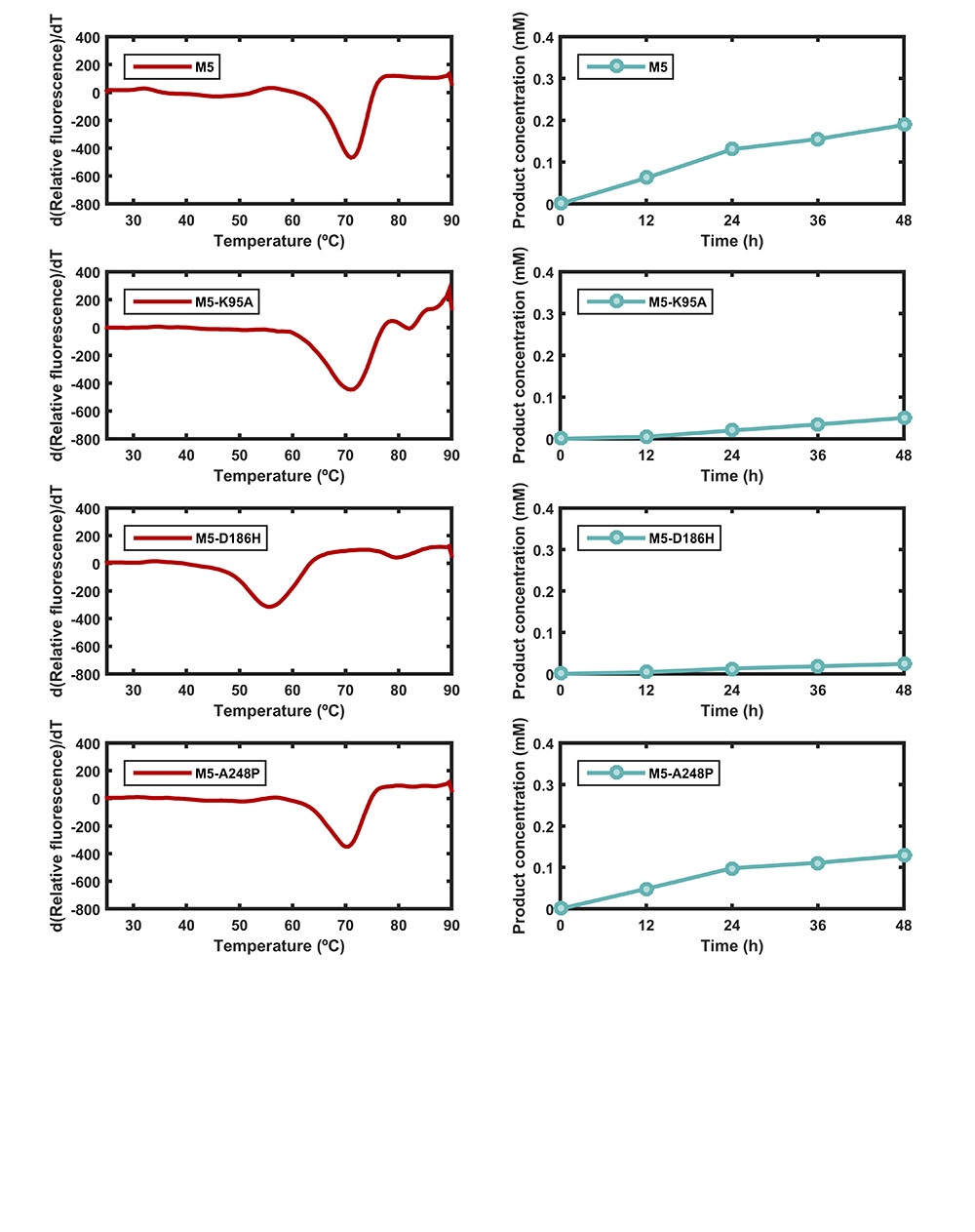

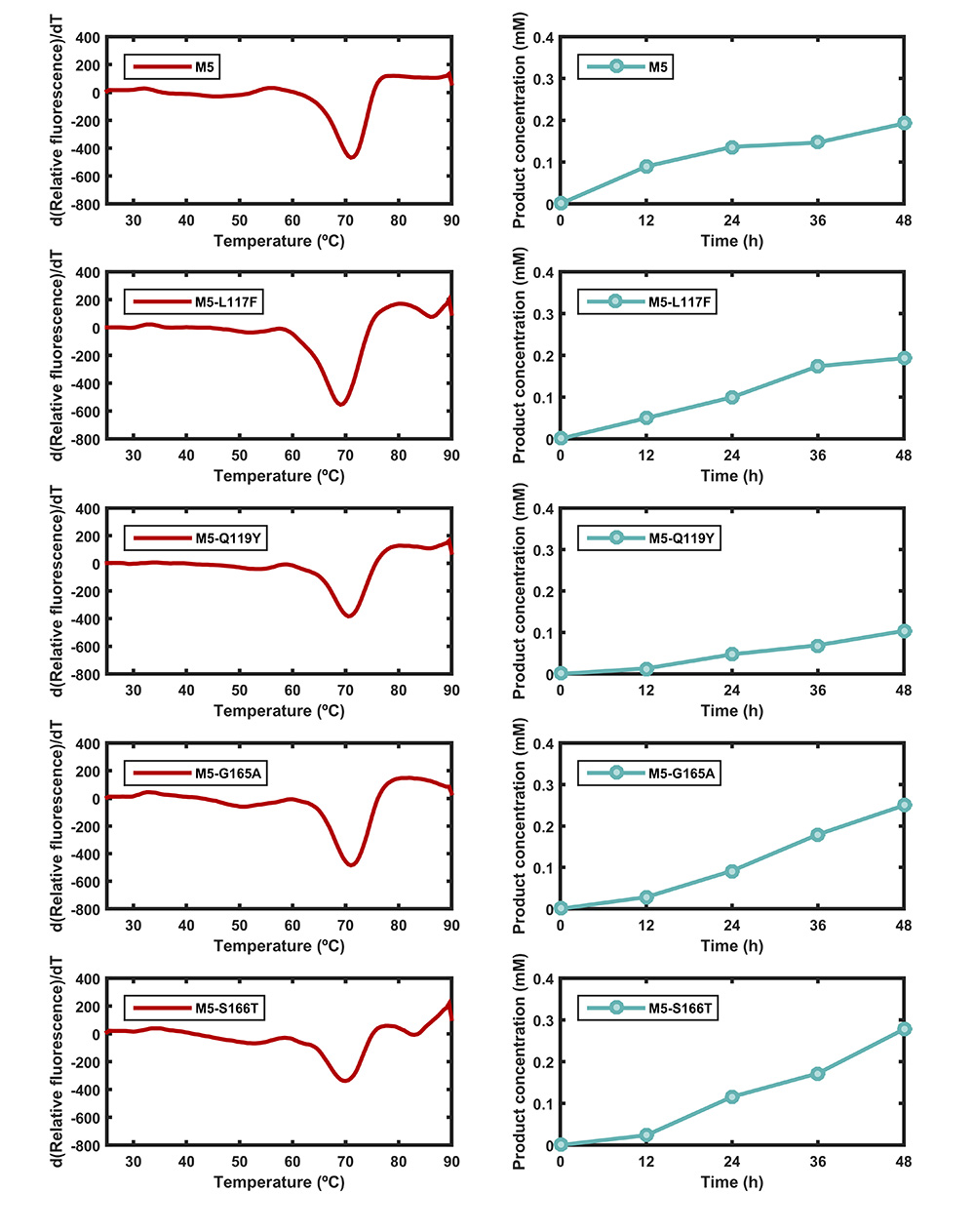

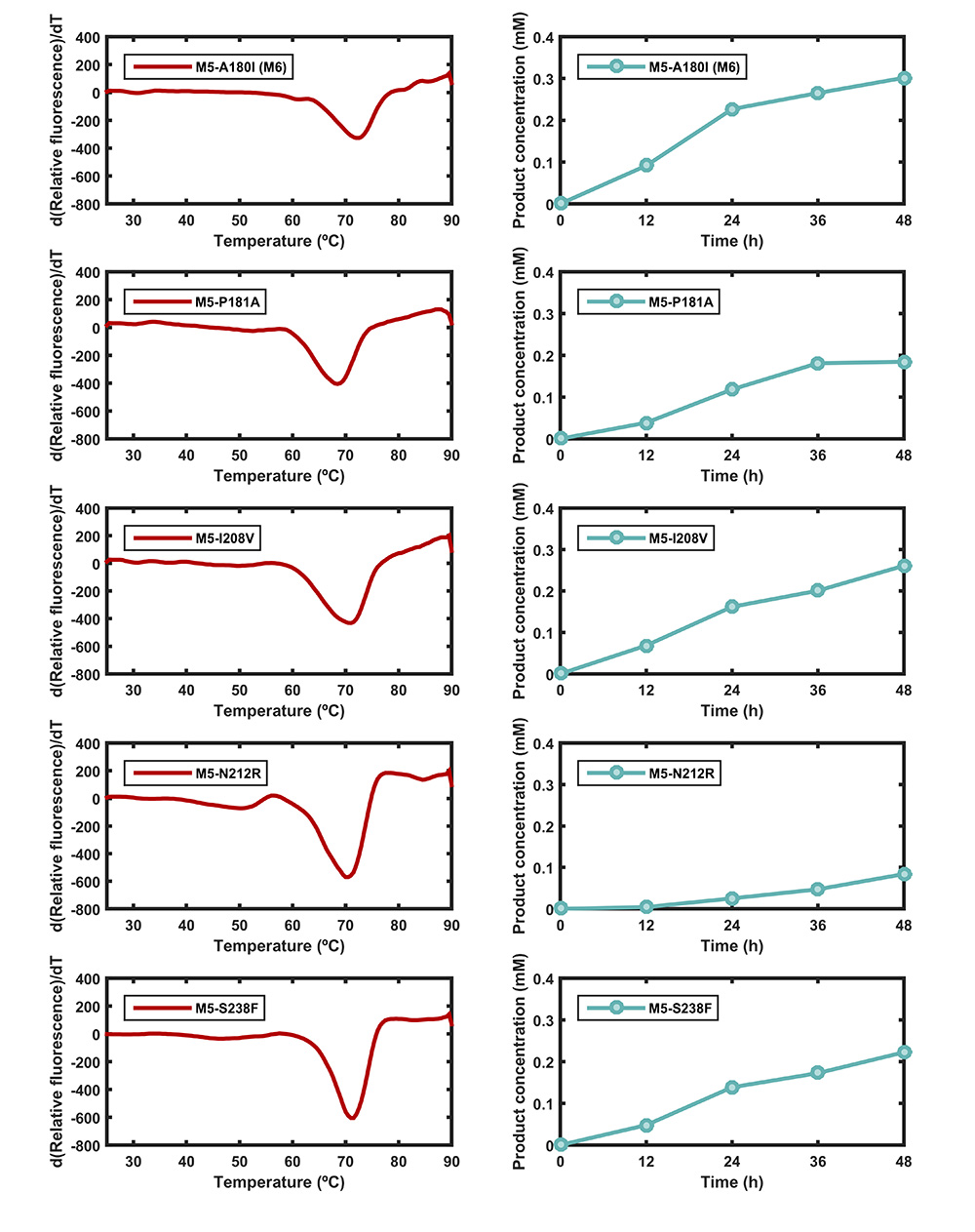

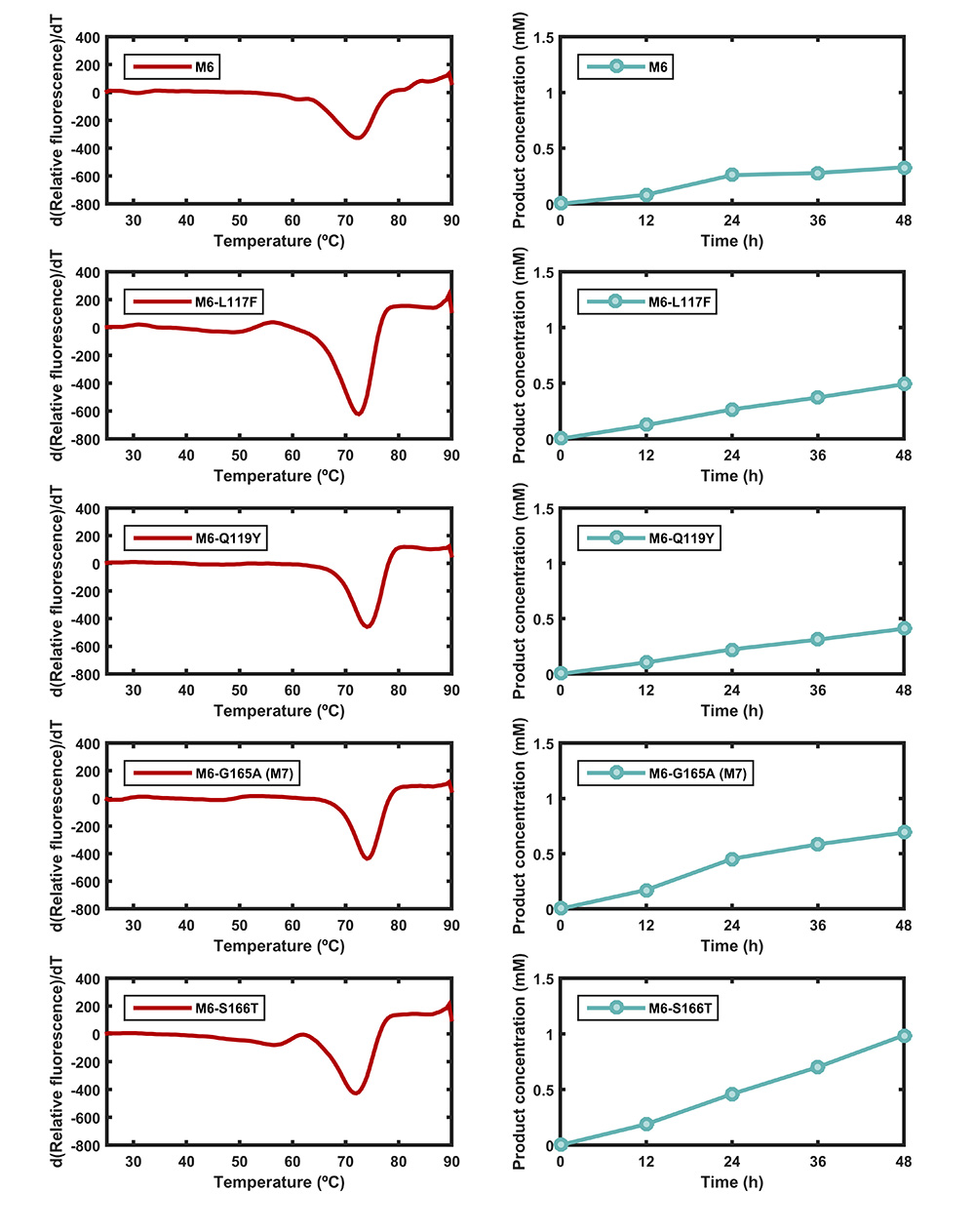

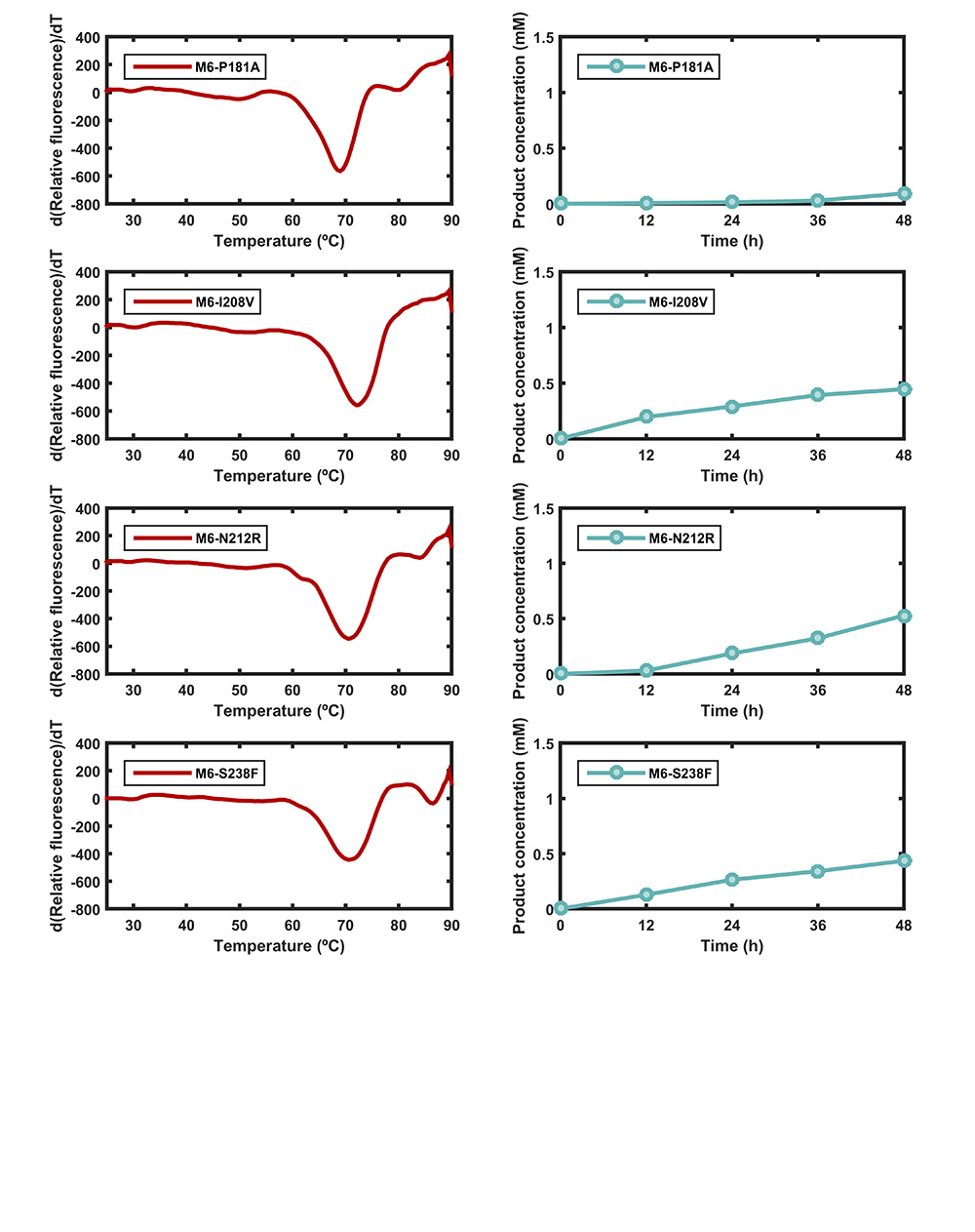

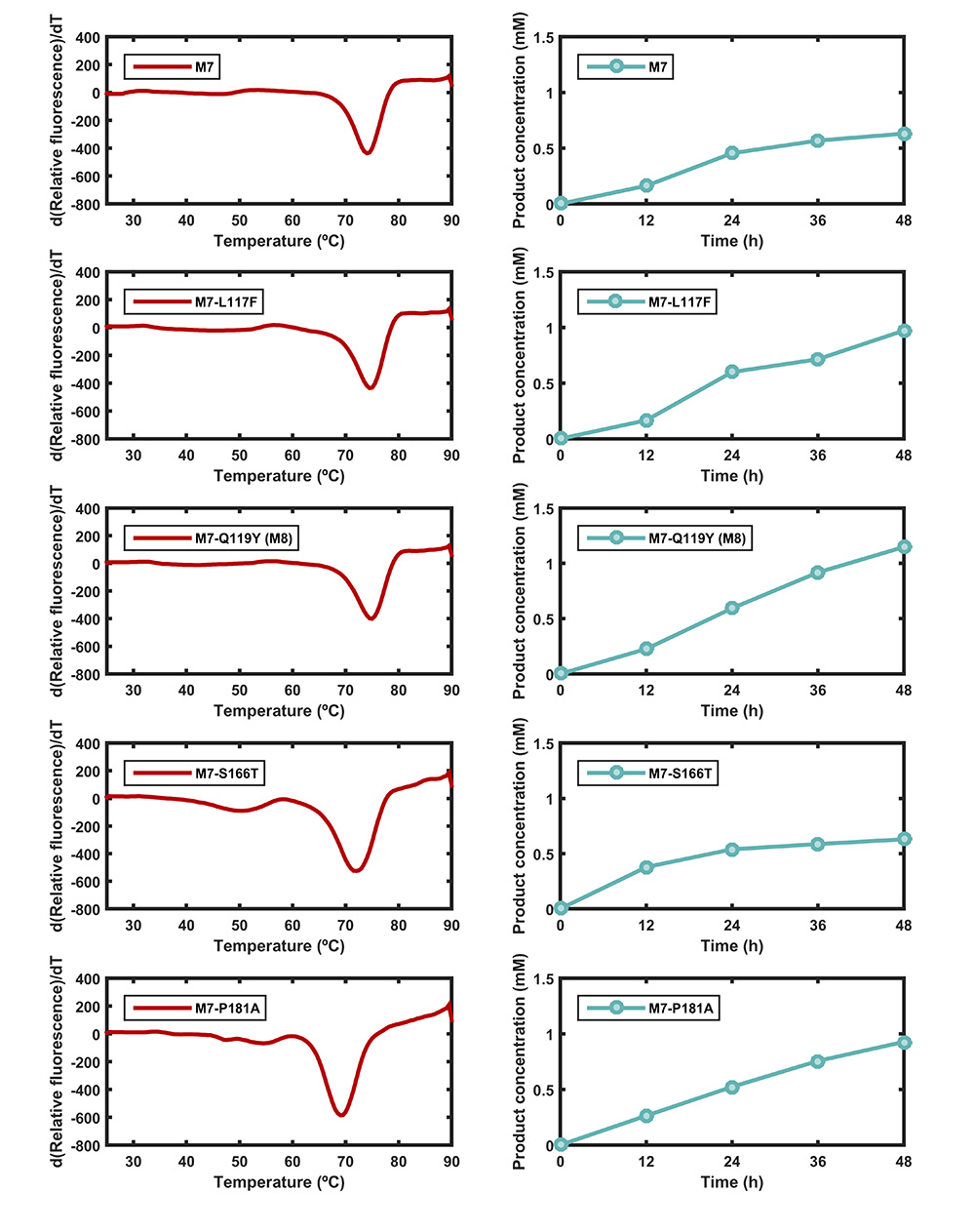

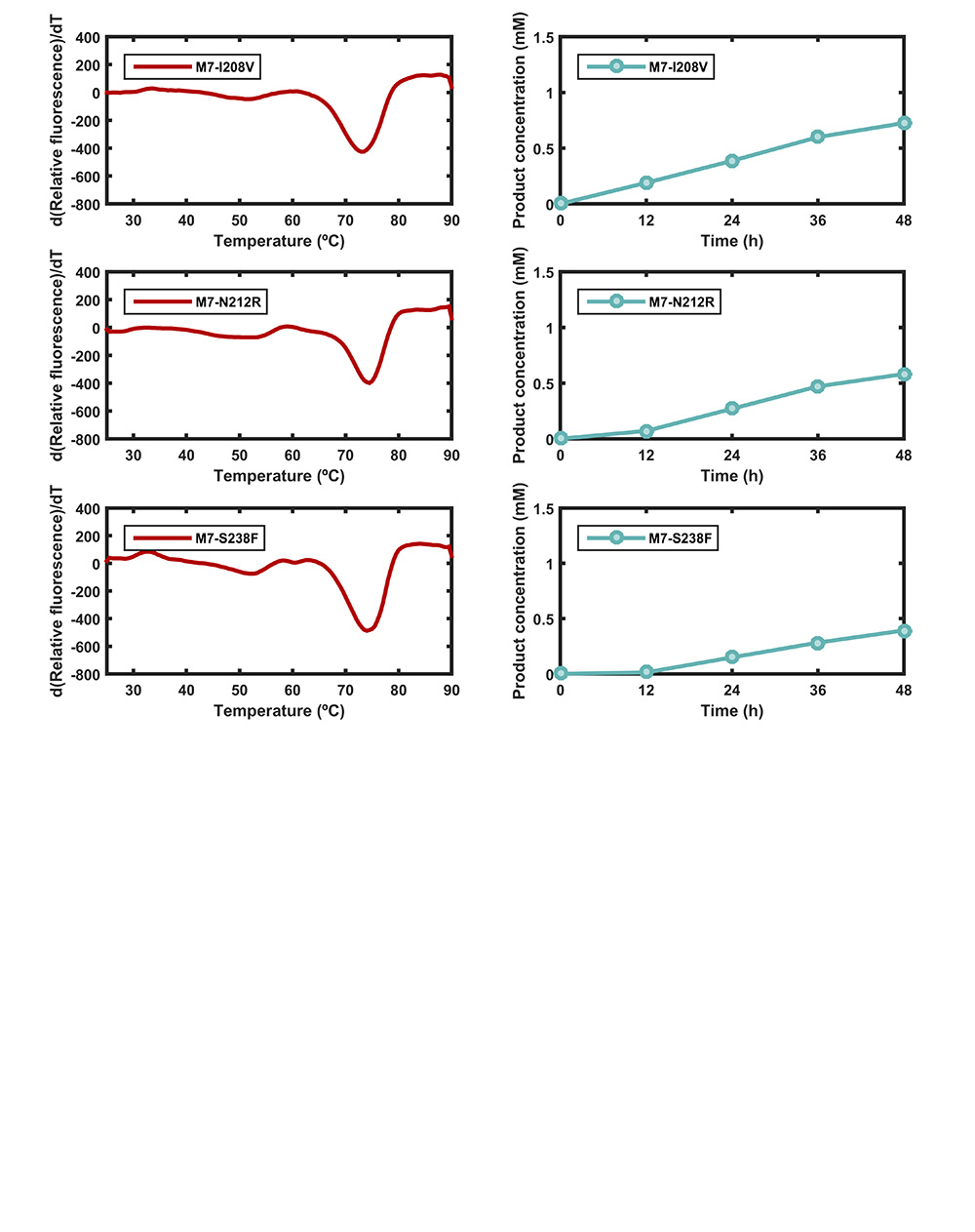

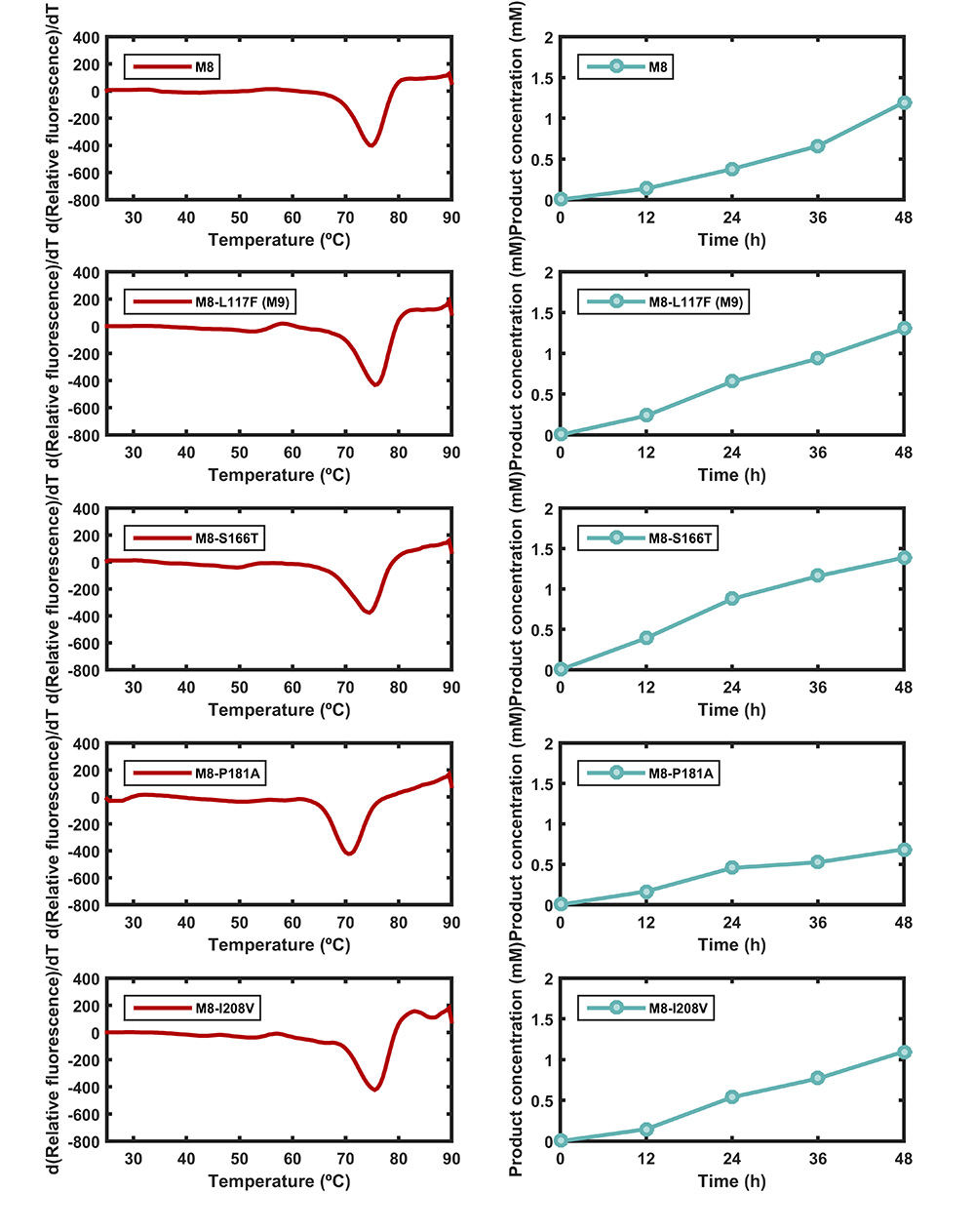

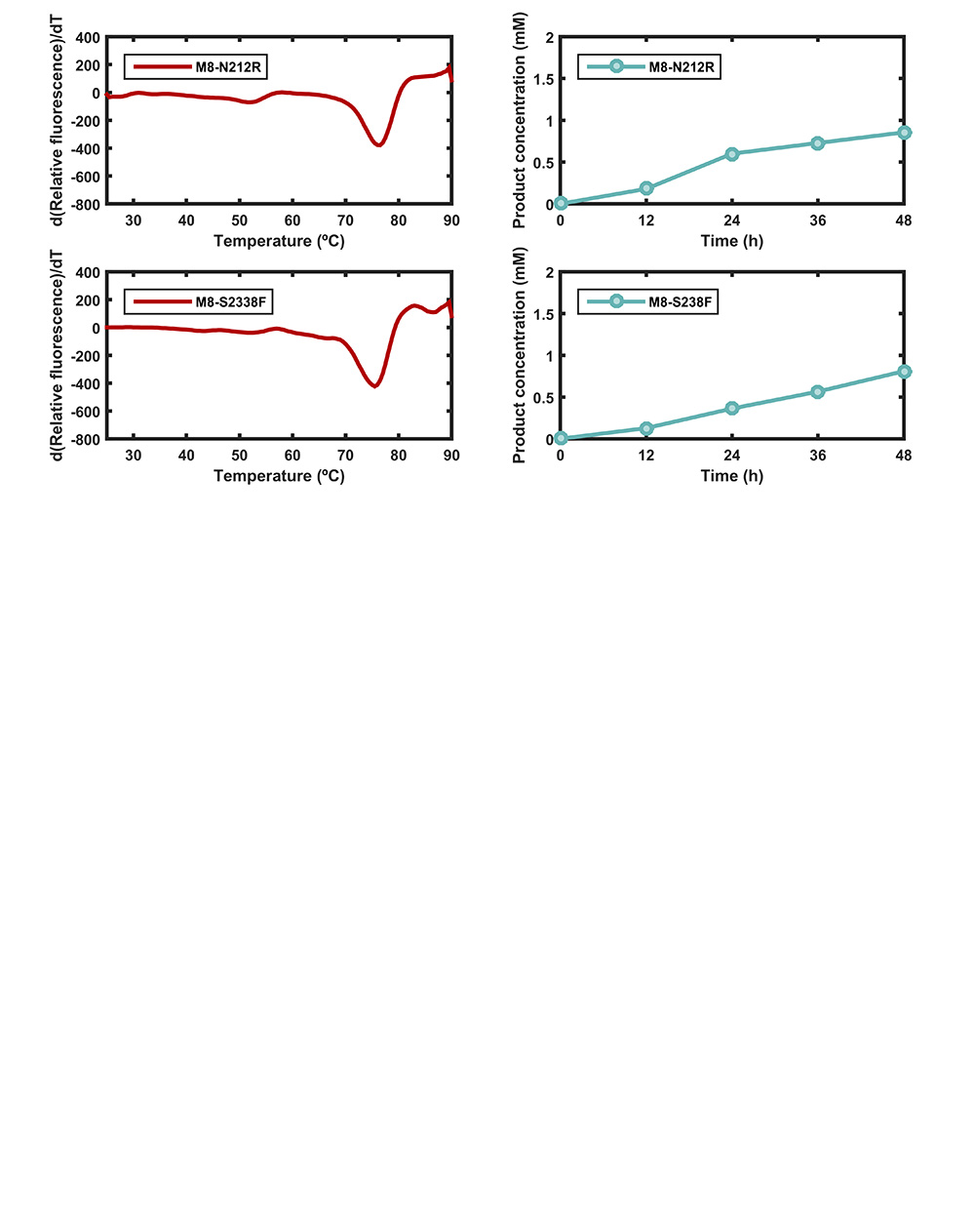

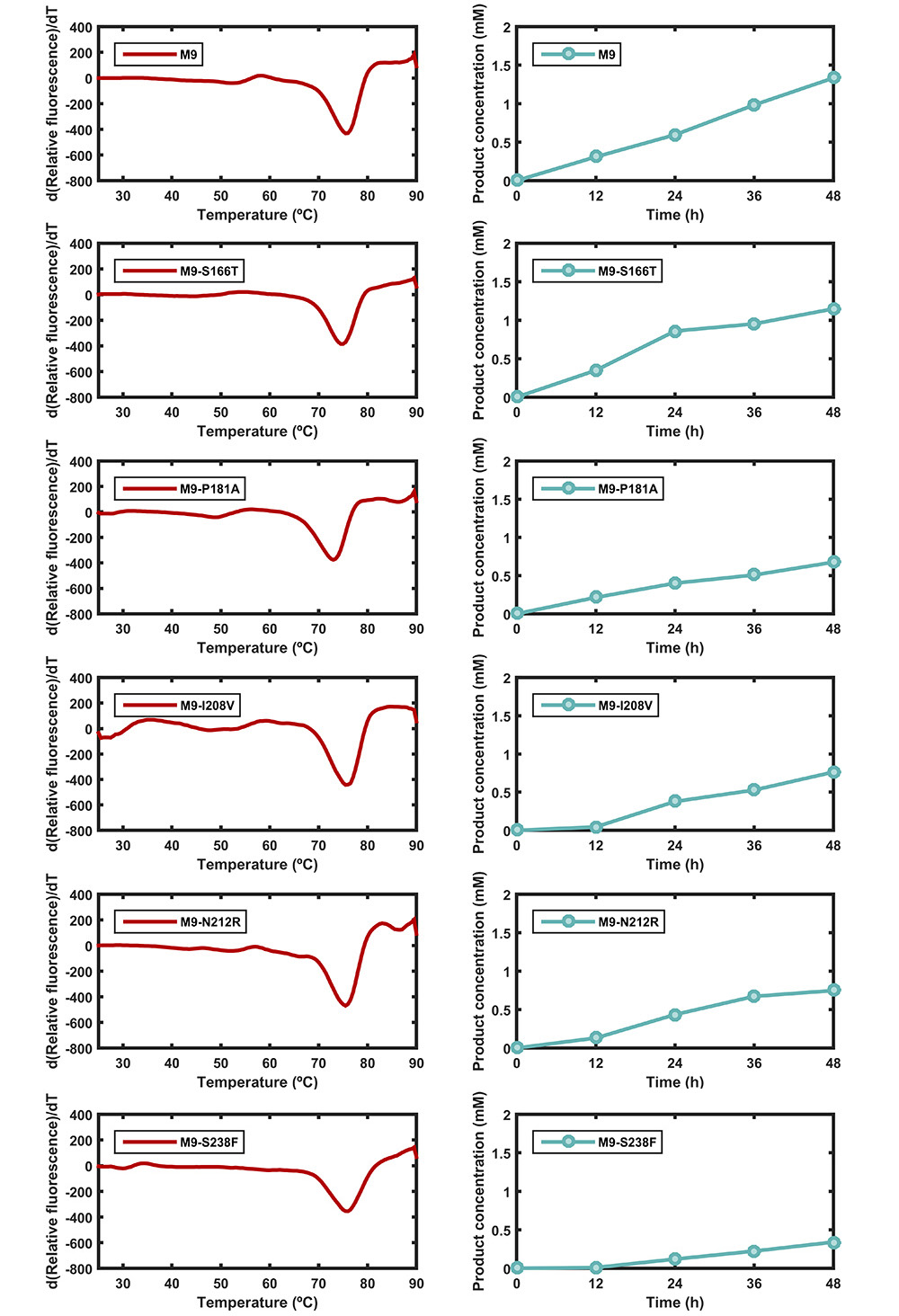

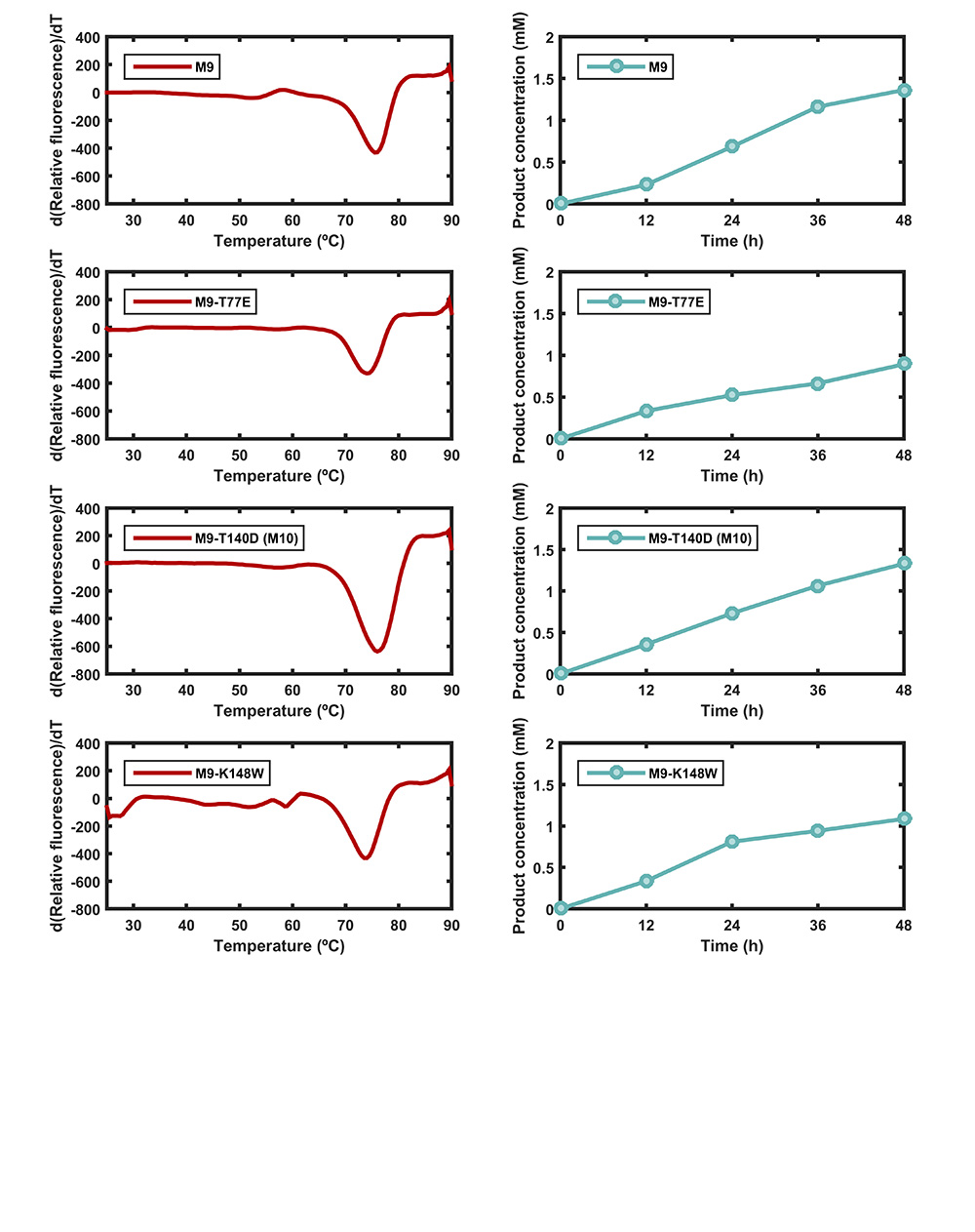

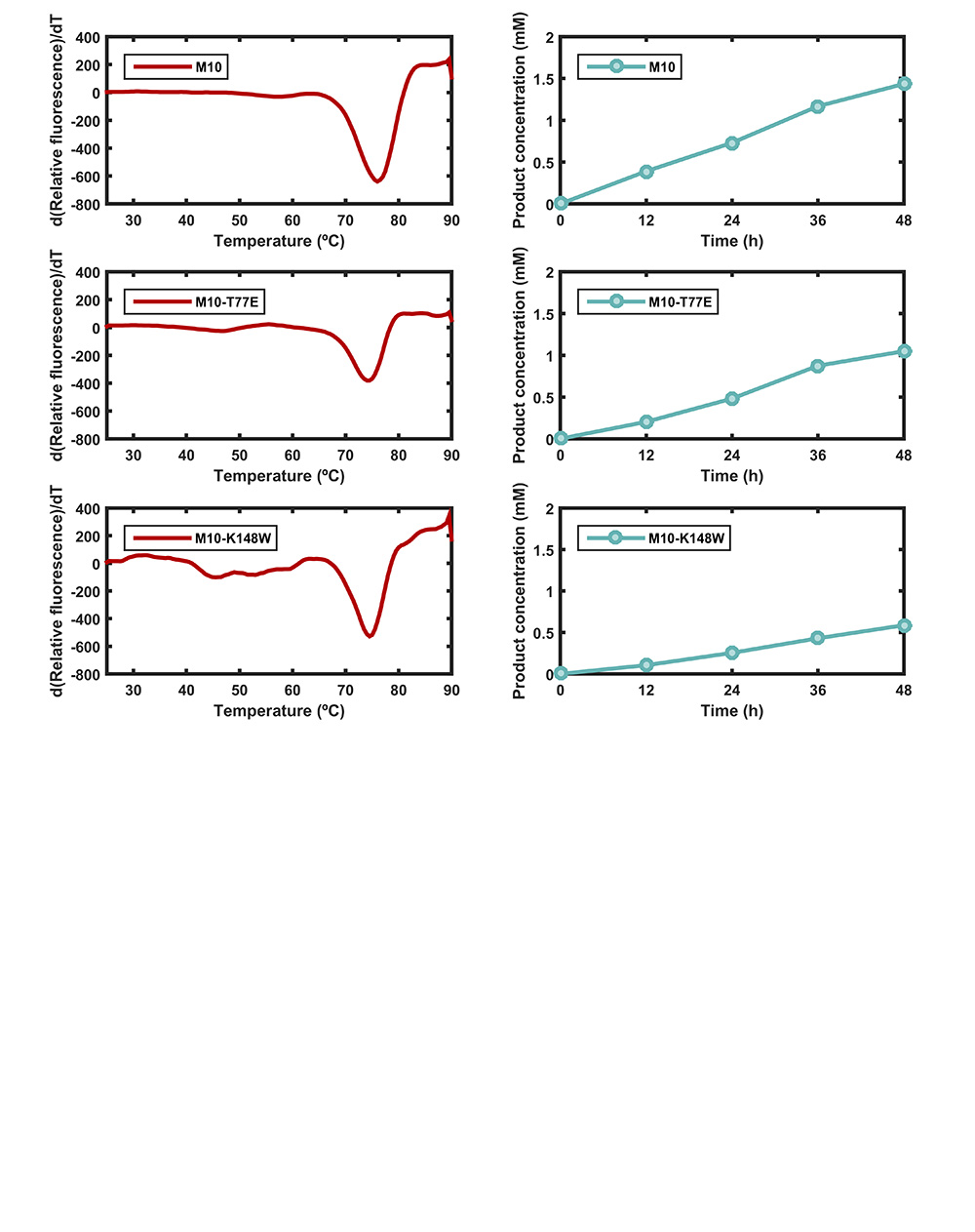

**Figure S4.** Left panel: *T*_M_ values of *Is*PETase and 66 variants during greedy accumulation steps. Right panel: PET degradation activity of *Is*PETase and 66 variants. PET film was incubated with 10 μL of enzyme (stock concentration 0.5 mg/mL) in 490 μL of 50 mM glycine-NaOH buffer (pH 9.0) at 37 ºC for 48 h. The product concentration is the sum of BHET, MHET and TPA.

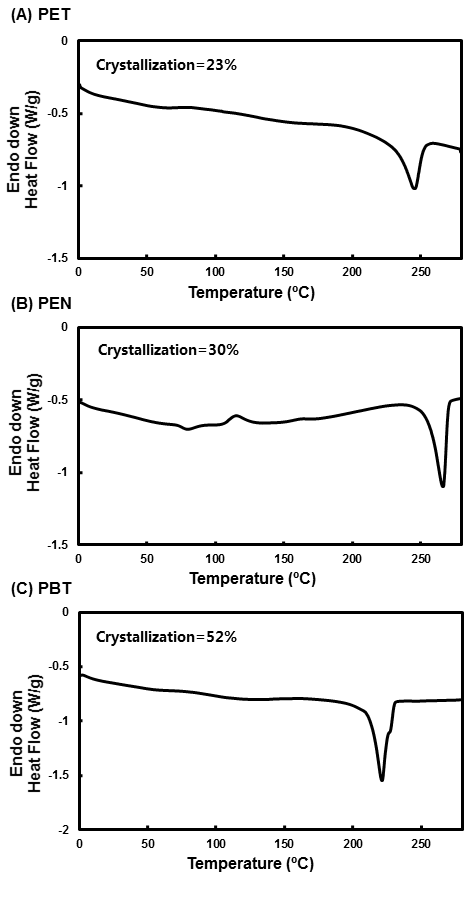

**Figure S5.** DSC for crystallinities of PET, PEN and PBY films. Crystallinities were calculated based on the heat of fusion of crystallites. The calculated crystallinities of PET film (A), PEN film (B) and PBT film (C) were 23%, 30% and 52%, respectively.

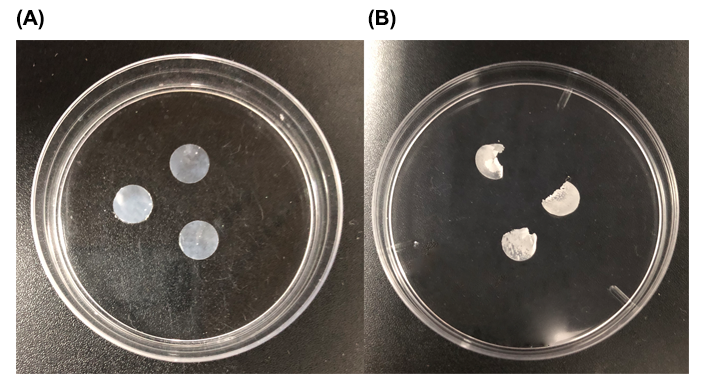

**Figure S6.** PET films before and after incubation with 10 μL of DuraPETase (stock concentration 0.5 mg/mL) in 25 μL of MeOH and 465 μL of 50 mM glycine-NaOH buffer (pH 9.0) at 37 ºC for 10 days.

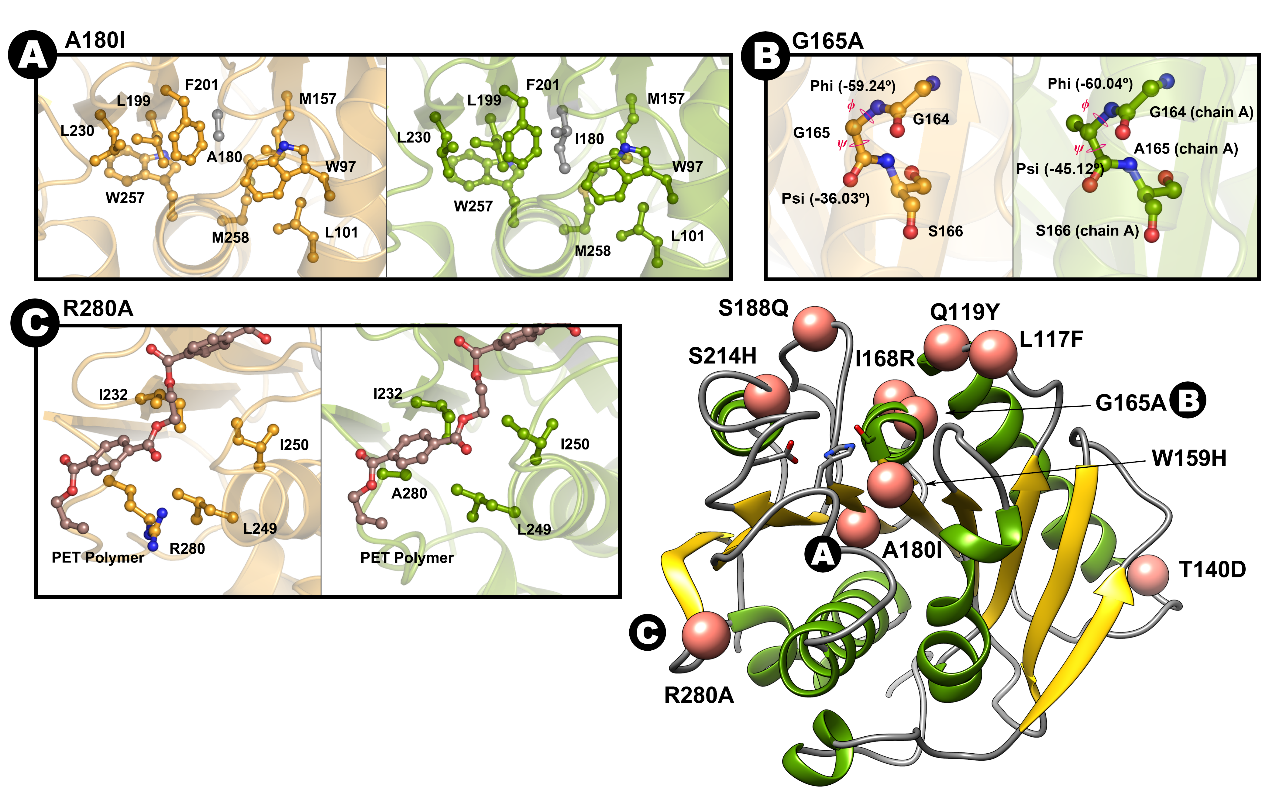

**Figure S7.** Location and structural effects of the stabilizing mutations of (A) A180I, (B) G165A, and (C) R280A. *Is*PETase and DuraPETase are shown in bright orange and green, respectively, in these images. Key residues proximal to the stabilizing mutations are presented as ball and stick representations. For the A180I mutation, the extended hydrophobic side chain filled the deeply buried hydrophobic core consisting of W97, L101, M157, L199, F201, L230, W257 and M258 to facilitate the interior hydrophobic interactions. Similar to the A180I mutation, R280A improved the hydrophobic interactions with I232, L249 and I250. Moreover, Kim et al.^2^ suggested that the increased activity of R280A was caused by the extended subsite IIc with a nonprotruding cleft. For the G165A mutation located at the α4 helix, the substitution of glycine with alanine facilitated the stabilization of α4 helix. The backbone dihedrals were rearranged from ϕ = -59.24º and ψ = -36.03º in the wild type to ϕ = -60.04º and ψ = -45.12º in the mutant chain A and ϕ = 61.15º and ψ = -43.71º in chain B.

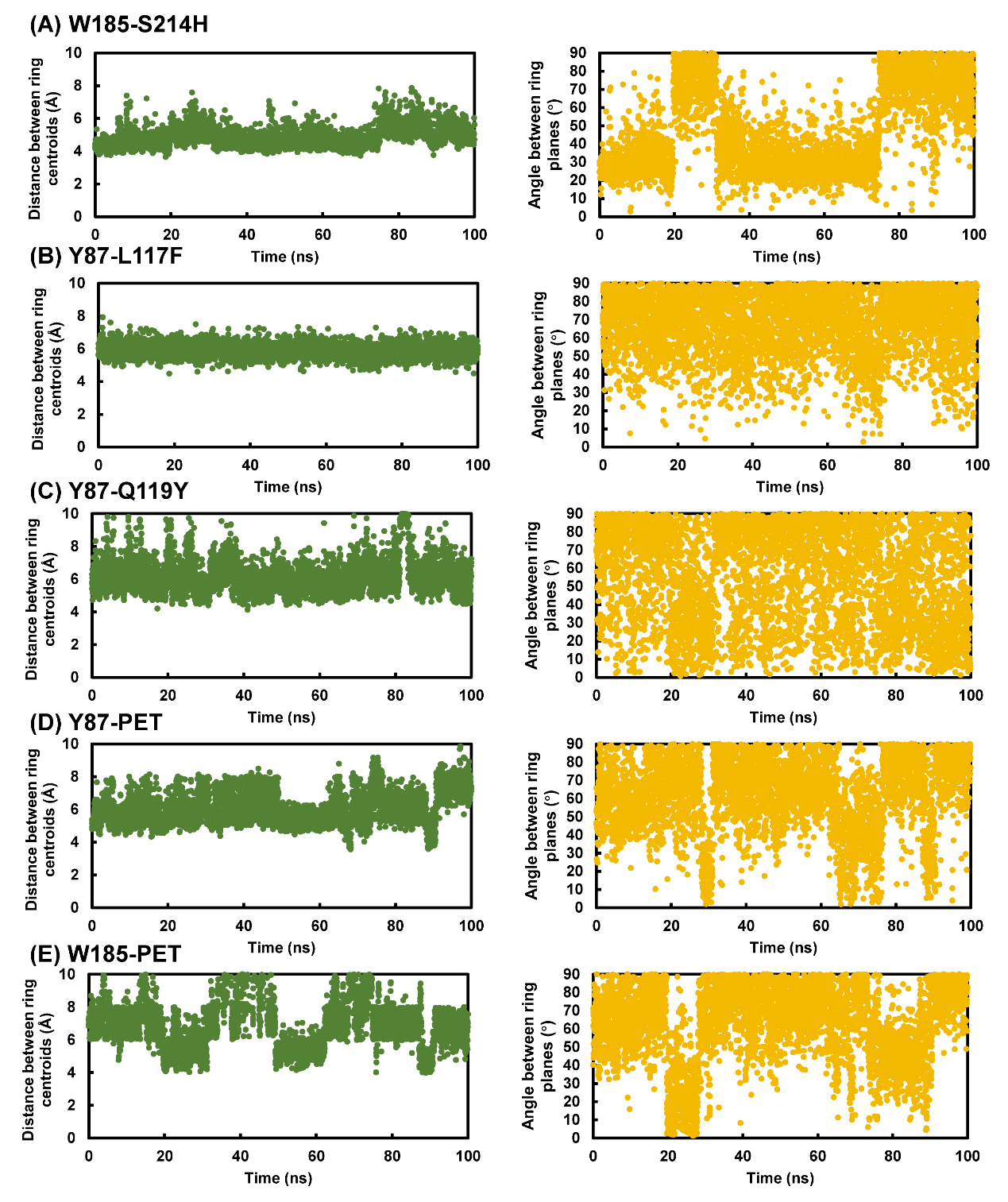

**Figure S8.** Time evolutions of the occupancies of π-π interactions during MD simulations. A π-π interaction is defined as an interaction between two aromatic rings in which the angle between the ring planes is less than 30° or between 60° and 120° and the distance between the ring centroids is less than 7 Å.
